# Transforming Esogastric Cancer Surgery Integrating SpiderMass Mass Spectrometry with Clinical and Microbiome Data for Margin Delineation and Prognosis

**DOI:** 10.1101/2025.10.13.682035

**Authors:** Léa Ledoux, Yanis Zirem, Alexandre Goosen, Florence Renaud, Charlotte Dufour, Julie Viezant, Cédric Lion, Guillaume Piessen, Michel Salzet, Isabelle Fournier

## Abstract

Esophageal-gastric cancers (EC) represent a significant global health concern, with esophageal cancer ranking seventh in terms of incidence and mortality worldwide. Gastric cancer is especially concerning, with an estimated one million new cases and 800,000 deaths annually. Late diagnoses often lead to poor outcomes, requiring critical interventions such as radical surgical resection with clear margins, in conjunction with chemotherapy, or radiotherapy to prevent recurrences and enhance survival. Thus, EC represents a significant clinical challenge, especially given the difficulty in achieving precise surgical margins in aggressive subtypes like poorly cohesive carcinoma (PCC). Moreover, pathological intraoperative margin assessment encounters significant issues, especially for PCCs, due to lacks of sensitivity for microscopic infiltration, potentially leading to recurrence and poorer patient outcomes. We address these critical limitations by integrating SpiderMass, an ambient mass spectrometry (MS) technology, with clinical metadata and microbiome profiling couple along with Machine learning. We demonstrate SpiderMass capability in real-time molecular margin delineation and identify distinct lipidomic and microbiome signatures correlating with tissue type and prognosis. Our integrative approach provides a more precise and biologically informative intraoperative diagnostic tool, significantly enhancing surgical decision-makin, to improve patient outcomes and extend survival.

## Introduction

The second most common cause of death in the world in 2022, with almost 20 million new cases and 10 million deaths, is cancer [1]. Gastric and esophageal cancers affect nearly 1.5 million people each year worldwide, composed of 968,784 new cases of gastric cancer and 511,054 of esophagus. The mortality rate of esophageal-gastric cancer (EC) is alarmingly high, with close to 1.1 million deaths per year, making it the fifth leading cause of death for gastric cancer (660,175 deaths) and the seventh for esophagus cancer (445,391 deaths). Both esophageal and gastric cancers are more prevalent in males than females, with 64.8% of gastric and 71.5% of esophagus cases in males [1]. Esophagus and gastric cancer share some risk and epidemiological factors due to their anatomic proximity. Adenocarcinoma and squamous cell carcinoma are the two most common subtypes of esophagus and esophageal- gastric cancers. Adenocarcinoma (AC) typically occurs in the lower esophagus or esophagogastric junction (EGJ), prevalent in the Western population (Northern Europe and North America). Risk factors include smoking, excess body weight, gastroesophageal reflux disease and esophageal intestinal metaplasia [2]. On the other hand, squamous-cell carcinoma (SCC), which is the most common type of esophageal carcinoma in the East Asia, South Central Asia and South Africa is found in the two-third upper part of the esophagus with a multifactorial aetiology including HPV as well as alcohol consumption, smoking, and their synergy [3]. Gastric cancer (GC) has several known etiological factors, including *H. pylori* infection as well as obesity and gastro-esophageal reflux, excessive alcohol intake, high salt consumption, and low fruit and vegetable intake. Excess body weight and gastro-esophageal reflux are also responsible for EGJ and the lower third of esophageal adenocarcinoma, as shown by their shared similarities in molecular features. Recent evidence shows a rising incidence of non-EGJ GC in individuals under 50, especially in low-incidence countries, where *H. pylori* prevalence is low. This may be linked to gastric microbiome dysbiosis, influenced by modern lifestyle and increasing autoimmune disorders [4]. 10% of GC shows familial aggregation, and only 3% are demonstrated with genetic predisposition. According to the 1965 Lauren classification, 95% of GC are gastric adenocarcinoma (GAC), which is classified between intestinal gastric cancer (IGC) and diffuse-type gastric cancer (DGC), with about 50% of GAC for 30% of DGC [5]. GAC with >25% of intestinal or diffuse types is classified as mixed-type adenocarcinoma and represents approximately 10% of cases. DGC and IGC are very different in their aetiology, histological, and molecular features. IGC is characterised by cohesive neoplastic cells organised in well-differentiated glandular structures, while DGC does not show or only little gland formation and diffusely infiltrates the gastric wall. If IGC incidence has largely decreased thanks to *H. pylori* treatment and improved dietary habits, DGC has been steadily increasing over the years. DGC, including poorly cohesive cell carcinoma (PCC) and signet-ring cell carcinoma (SRC) is more prevalent in younger and females while IGC is associated with older and male patients [6]. There have been several changes in the EC classification specifically in the definition of DGC and PCC. PCC is composed of isolated or small islets of neoplastic cells, but the SRC subtype is characterized by cells presenting a central optically clear, globoid droplet of cytoplasmic mucin with an eccentrically placed nucleus. In the consensus definition, DGC is classified as SRC if >90% of cells present the specific SRC type. Alternatively, if <90% of SRC-cell type is present but still >10%, the tumour is considered as PCC with SRC component and if <10% SRC-cell type is present then it is classified as PCC-NOS (non-other specified) [7].

Unfortunately, GC is most often asymptomatic with nonspecific, latent symptoms at the early stages, and about 65% of patients are diagnosed at an advanced stage (stage III or IV) resulting in a poor prognosis. The 5-year overall survival remains less than 50%, even with intensive perioperative treatment [8]. Indeed, for patients with advanced GC or GOJC, the survival rate at 5-years is only 6% [9]. Studies have shown that early and accurate detection of this cancer could increase the five-year survival rate to around 90% [10]. Endoscopy plays an essential role in diagnosing EC and relies on forceps biopsies; 5 to 8 biopsies are generally necessary to reach a sufficient representation of the tumour and get its histological and/or molecular characterisation. However, recent studies have shown that the detection accuracy of conventional endoscopy is only 79% [11]. Although neoadjuvant and/or adjuvant therapies such as chemoradiotherapy, chemotherapy or targeted therapies, including immunotherapy, can improve the survival of EC patients to some extent, radical surgery remains the main option for treating non-metastatic EC cancers [12]. Only patients with advanced metastatic unresectable tumours will exclusively receive first-line conventional chemotherapy with platinum-fluoropyrimidine doublet followed targeted therapies with anti-HER2 or PDL1 according to the molecular features found by the IHC or ISH [12]. Curative resection with microscopically negative margins has been accepted as the most effective treatment based on surgical philosophy, as even minimal numbers of remaining cancer cells could favour recurrences [13]. Indeed, it is widely recognised that complete removal of the tumour is associated with better prospects of cure for most solid cancers. On the other hand, positive margins [14] are associated with an increased risk of local recurrence and reduced overall survival in many types of cancer, and radical tumour-free margins (R0) should be achieved.

Total gastrectomy doesn’t lead to better patient outcomes, though and distal gastrectomy is associated with better outcomes and fewer morbidities, demonstrating the importance of reaching the right balance between R0 and organ sparing [15]. To avoid leaving positive margins, a frozen section intraoperative analysis of the longitudinal and/or circumferential may be necessary. This pathological analysis is carried out as close as possible to the operating theatre from a frozen, stained section of the resected material that is examined by the pathologist during the surgery. The results of this extemporaneous pathological assessment of the margins are then communicated to the surgeon to guide future decisions. However, this exam generally requires half to 1h and shows limited reliability due to sampling error since only a fraction of the tissues are analysed and to lower quality of the exams that can be conducted by comparison to post-surgery with the use of IHC. This leads to discrepancies between the post-surgical pathological assessment and the intraoperative one with only 67% sensitivity and 93% accuracy on proximal margins even if frozen sections can be used and significantly higher error rates in the cases of the PCCs subtypes for which particular vigilance is required [16]. The importance of reaching R0 resection margins and finding the loco-regional extension of the tumour (positive sentinel lymph node, metastasis in peritoneal cavity) and the difficulties met with the gold standard practices emphasise the need for novel technologies to be developed to enable in-situ and real-time analysis. Various imaging technologies are currently under development and evaluation for surgical guidance. Intraoperative MRI and ultrasound have proven their interest for a while, but more recently, spectroscopy imaging has also shown promising results considering acquisition time (i.e., 10 min) and accuracy. They include hyperspectral imaging (HIS), which shows accuracies lying between 81 and 94%, Raman imaging with Stimulated Raman Scattering (SRS), fluorescence imaging, optical coherence tomography (OCT) and optoacoustic [17]. Over the past decade, Mass Spectrometry (MS) has also appeared as an interesting technology for assistance with surgery. Despite MS has been historically developed to analyse extracts, the introduction of Ambient Ionization Mass Spectrometry (AIMS) has promoted the creation of a few new technologies enabling *in vivo* analysis and dedicated to intraoperative assessment [18]. If MS is larger than other spectroscopic or acoustic systems, it has the clear advantage of showing a very accurate description of the molecular content of the analysed tissues [19]. Our group has worked since 2011 on the development of a laser-based technology for intraoperative *in vivo* real-time analysis SpiderMass, which is non-invasive [20]. It has been developed to help surgeons in the operating theatre with decision-making based on real-time diagnosis thanks to non-target molecular analysis [20,21]. The MS molecular profiles collected with the technology are highly specific to the cell population or sub-population phenotypes, and combining SpiderMass with machine learning (ML) offers a possible interpretation of the MS spectra for the technology to be used in the surgery context. SpiderMass cell phenotyping is yet based on metabolites and lipids, and the performance of the technology has been demonstrated for various pathologies *ex vivo,* including skin diseases and different cancers, among which glioblastoma [22], ovarian fimbria pre-neoplastic lesions [23], head and neck cancer [24] and sarcoma [25]. The technology was also proof-of-concept *in vivo* on patients with skin pathologies received at a dermatological referral centre (NCT04472546) and on dog patients with sarcoma in the veterinary room [26], demonstrating its innocuity and performance in the real environment. More recently, we have augmented the technology with imaging capabilities by connection to a high-accuracy robotic arm [27] to enable possible interpretation of a complete tissue area, whether suspect ones, in-situ margins after resections or resected tissues. Additionally, we have also assessed the capacity of the SpiderMass to discriminate the immune cells from the cancer cells and to discriminate between the different immune cell populations to use the technology to characterise and study the tumour microenvironment (TME), creating an immunescore based on SpiderMass data to move forward real-time prognosis [28]. Interestingly, the capacity to detect specific cell populations which are infiltrated within the tissue from SpiderMass images demonstrates the potential of the technology to address complex EC cancers such as the DGCs, and particularly the PCCs with application to surgical margin definition and evaluation of bacterial communities. Indeed, SpiderMass has also the ability to detect and classify bacteria as we previously described [29]. In this context, this technology can also be used to track microbiota linked to EC.

Microbiota is known to play a key role, and modification in its composition is associated with dysbiosis and contributes to the development of neoplasia. Significant enrichment of *Streptococcus* and a decrease in the abundance of *Lactobacillus* are found in the tumour while in healthy controls, the microbial community is characterised by the enrichment of several genera, including *Peptostreptococcus*, *Prevotella*, *Actinobacillus*, *Gemella* and *Rothia* [30] establishing a link between disturbances in the microbiota and cancer. With the development of molecular biology and metagenomics, scientists have a more complete understanding of gastrointestinal bacteria, and it is now accepted that microbial dysbiosis can promote the onset of gastric cancer through various mechanisms [31], indicating that other bacteria than *H. pylori* also play an important role in gastric cancer. Yet if studies have identified differences in gastric microbiota composition and function between gastric cancer patients and control groups, there is no consensus on the specific microbial taxa involved in gastric cancer pathogenesis. Additionally, bacterial heterogeneity at the scale of the tumour microenvironment or across different GC subtypes remains unclear [32]. Some studies point to the involvement of different bacterial communities in DGC by comparison to IGC since treatment of *H. pylori* have led to a decrease in IGC whilst DGC is steadily increasing [33]. Finding bacterial community differences for DGCs including PCCs paves the way for better management of these cancers which show poor prognosis. Thus, there is growing recognition that microbiome composition within gastric tissues might influence tumorigenesis and clinical outcomes. However, interpreting microbiome data in cancer studies is complex due to potential confounders, including the impact of antibiotic use and proton pump inhibitor (PPI) therapy, which can alter gastric microbiota independently of cancer. Consequently, novel diagnostic tools integrating molecular pathology and microbiome insights are urgently required. SpiderMass, an advanced ambient MS technique, provides real-time, high-resolution molecular analysis with minimal sample preparation, representing a promising solution. The present study evaluates SpiderMass’s clinical potential and introduces a novel microbial quantification strategy (bacterioscore), enhancing intraoperative diagnostics and prognostication. in the management of gastric cancer (**Fig. 1**).

**Figure 1.**
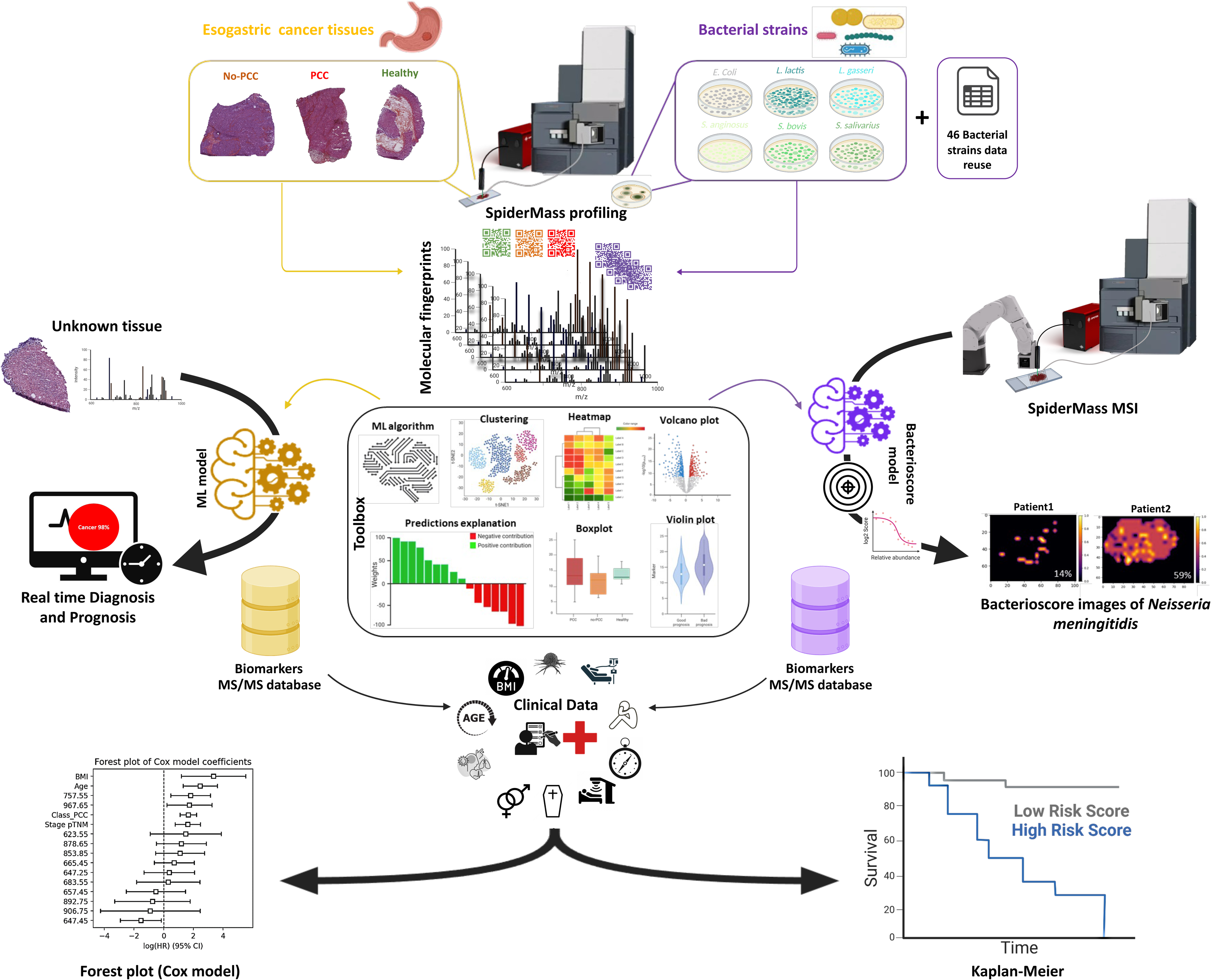
Methodological Workflow based on SpiderMass technology used for esogastric cancer. Esogastric tissue samples (PCC, non-PCC, healthy) and selected bacterial strains are analyzed using SpiderMass technology. Spectral data are processed with ML and dimensionality reduction techniques (e.g., UMAP) to develop classification models. Once trained, the models are applied blindly to new datasets to assess prediction performance. Statistical analyses including Cox proportional hazards models and Kaplan–Meier survival curves are used to evaluate prognostic factors. Additionally, microbial data are used to generate a Bacterioscore, contributing to patient risk stratification and supporting personalized treatment strategies.

## Results

### Cohort characteristics

The total EC cohort included 108 patients (80 males, 27 females) with an average age of 63 years (range: 24–86) for a total of 164 tissue samples, of which 77 correspond to healthy and 87 to adenocarcinomas distributed over 60 no-PCC and 27 PCC subtypes (**Table S1-S2** and **Fig. S1-3**). Tumour locations were distributed as follows: 55 in the stomach, 39 in the esophagogastric junction (EGJ), and 13 in the esophagus. Gastric reflux was observed in 21 patients. BMI analysis classified patients as lean (5), normal (49), overweight (30), moderately obese (17), highly obese (1), and morbidly obese (1). Only 60 of the 107 patients received neo-adjuvant chemotherapy to reduce the size of the tumour prior to surgery, and 10 underwent a more specific radiochemotherapy scheme. To greatly reduce the risk of recurrence or metastasis, 76 patients underwent adjuvant chemotherapy after surgical resection of the tumour. Among the 107 patients in this cohort, 49 suffered a relapse. So far, 70 patients have passed away, while 37 are still alive. Median overall survival (OS) was 40 months (range: 1 day–160 months).

An additional pilot cohort of mixed EC tissues was used for margin delineation assessment. This cohort comprises 5 tissue samples, each containing both healthy and cancerous regions, obtained from 5 patients - 2 women and 3 men - with an average age of 55 years. Among them, 3 had tumours located in the stomach while 2 had tumours in the EGJ. All patients had a BMI ranging from 20 to 27, indicative of normal body weight, with only one patient experiencing gastric reflux. All patients received adjuvant treatment, with only 4 also undergoing neoadjuvant therapy. Among the 5 patients, 2 experienced disease relapses, and so far, only 2 have deceased. The median OS for this mixed cohort is 64 months (range: 13 to 130 months).

### Assessment of EC using SpiderMass-based diagnosis

#### Automated diagnosis of EC by blind predictions including EC subtyping

SpiderMass high capabilities for cell phenotyping relies on using all the measured markers and their relative intensities in the MS spectra as a barcoding system (**Fig. 2A**). This necessitates training through ML approach to associate a cell class with a specific profile. Thus, to assess the diagnosis performances of the SpiderMass technology for EC, an ML-based classification model was trained, and its performance was evaluated to discriminate cancer versus non-cancer cells. Indeed, the EC cohort was divided into two groups, one for training and the second for validation, composed of 132 and 32 tissues, respectively. The classification model was built from the training dataset using the RidgeClassifier ML algorithm, due to its robust performance in high-dimensional data handling and interpretability, to achieve optimal classification, resulting in a 100% accuracy for both MS ion modes. Cross-validations (CV), either 5 or 20-fold, indicated strong accuracy (>90%) in distinguishing cancer from normal tissue. Using 20-fold CV to ensure the model’s robustness and generalizability, the accuracy of the built model was 93% and 90% in negative and positive ion mode (**Table 1 and Fig. 2B-S4**).

**Figure 2.**
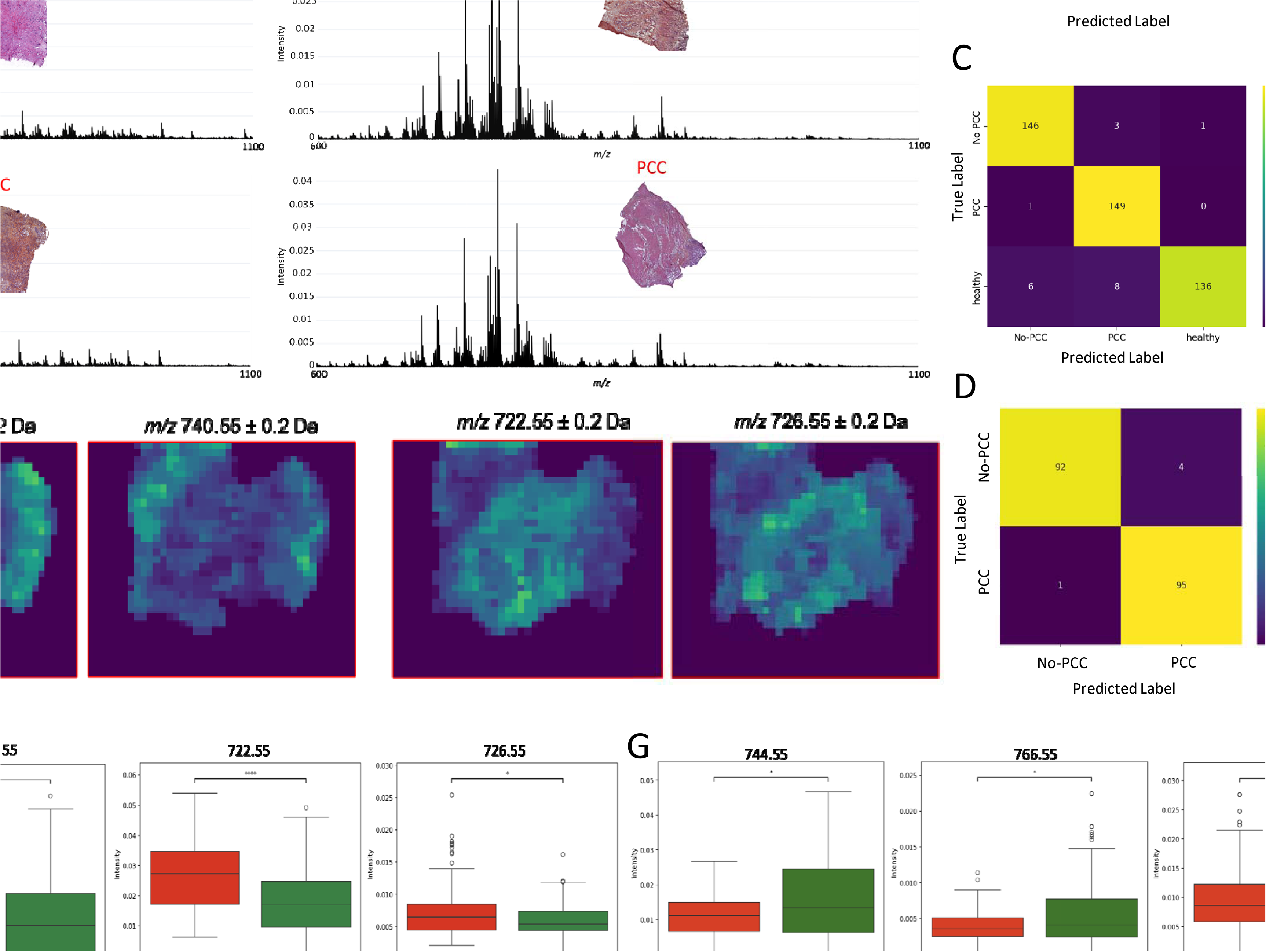
Automated molecular-based diagnosis of esogastric cancer. **(A)** Mean spectra of each type of esogastric tissues; healthy, non-PCC and PCC in negative (left) and positive ion mode (right). **(B-C-D)** Classification report and confusion matrix of RidgeClassifier after 20- fold cross-validation, obtained for the classification model cancer vs healthy in negative ion mode, non-PCC, PCC and healthy in positive ion mode and PCC vs non-PCC in positive ion mode. **(E)** Distribution of 4 biomarkers in mixed EC tissue imaged by SpiderMass. As depicted, the arrangement of these specific ions distinctly outlines the healthy and cancerous areas in the tissue sample, thereby confirming the trustworthiness of these ions as reliable biomarkers. **(F-G)** Corresponding boxplot of 8 confident biomarkers specific of cancer and healthy tissue in negative and positive ion mode. **(H-I)** Corresponding boxplot of 8 confident biomarkers specific of PCC and non-PCC subtypes in negative and positive ion mode. Related to Tables S1-S2-S3 and Fig. S1-S2-S3-S4-S5-S6.

**Table 1.**
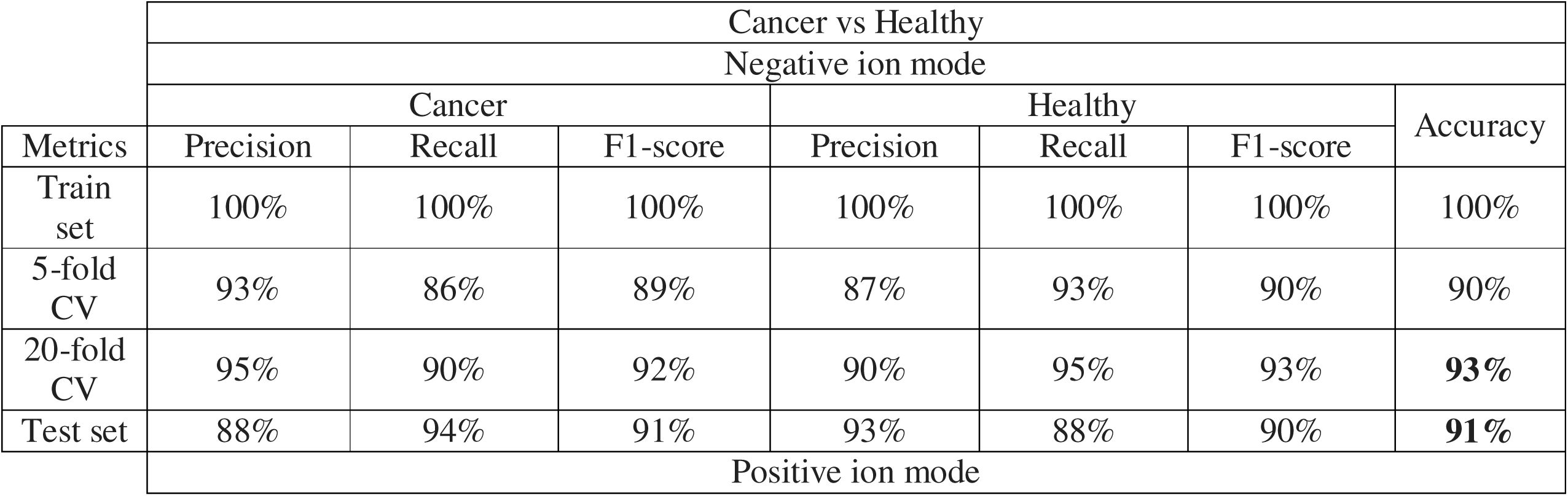
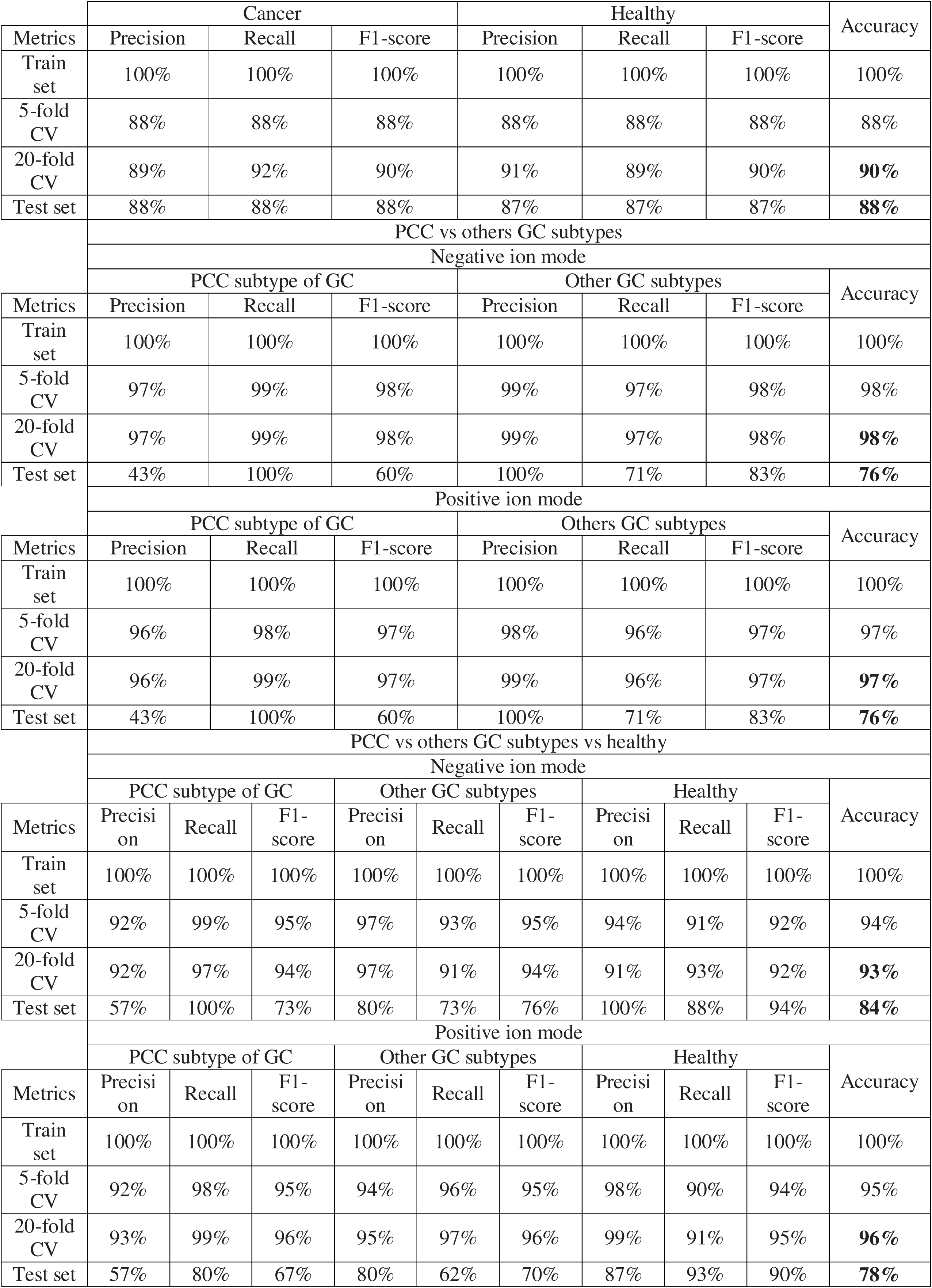
Overview of the results of the classification models. Classification models were built based on the association of SpiderMass spectra with the class of the analyzed tissue. Models were built using Ridge classifier and further cross-validated using 5- and 20-folds cross-validation. Classifications were obtained from both positive and negative ion mode SpiderMass data. In addition, a test set were used to evaluate the models in blind.

To further assess the performance of the classification model in both ion modes, blind predictions were conducted on the independent validation group. The tissues were subjected to SpiderMass analysis in a blinded manner and compared against the pathologist annotations. In negative ion mode, only 4 out of 32 analytical spots were misclassified, resulting in a commendable accuracy rate of 88%. Conversely, in positive ion mode, only 5 out of 32 were inaccurately classified, yielding an accuracy rate of 84%. It’s worth noting that there were specific instances where misclassifications could be justified. For example, there was a case where a tissue initially labelled as healthy in the FREGAT database was subsequently deemed cancerous by the pathologist. SpiderMass prediction, which indicated the tissue as healthy, aligned with the initial data, casting doubt on the pathologist’s annotation. Taking this exceptional case into account, SpiderMass blind predictions allowed for the accurate classification of 29 out of 32 in negative ion mode, resulting in a correct prediction rate of 91%. A slightly better prediction rate of 88% was obtained in positive ion mode (**Fig. S4**).

In the second phase, we checked the performance of the technology to specifically diagnose the presence of PCC versus other types of esogastric cancers (referenced here as non-PCCs) knowing to the difficulties raised by this subtype for intraoperative analysis and considering its aggressive nature. We first built, using the same supervised learning approach, a model based on three classes with healthy tissues, non-PCCs and PCCs esogastric cancer subtypes. The classification model achieves a respectable accuracy of 93% and 96% after 20-fold CV for negative and positive ion modes, respectively (**Table 1 and Fig. 2C -S5**). Although the blind results were relatively low, they remained respectable, with correct classification rates of 84% and 78% (**Fig. S5**). This was compared to a second classification model with only two classes built to discriminate between PCCs and other subtypes (non-PCCs). This model showed higher accuracy, reaching correct classification rates of 97% and 98% for both ion modes (**Table 1 and Fig. 2D -S6**). This result highlights a dual strategy that appears to be effective, i.e., a dual classification model that first determines whether the tissue is cancerous or healthy with an accuracy of 93%, and then if the tissue is cancerous, the second model determines the subtype with an accuracy of 98%. However, blind diagnosis allowed to well classify 13 of the 17 unknown tissues, giving an accuracy of 76% (**Fig. S6**).

#### Annotation of EC no-PCC and PCC lipids

Our interest extended more specifically to the identification of lipids linked to the distinction between cancerous and healthy esogastric tissues to enhance our understanding of the biological mechanisms responsible for this differentiation. The use of a supervised approach combining LIME algorithm and unsupervised statistical analysis [28,34] allowed us to generate a comprehensive list of confident discriminative lipids. In total, 50 of them were identified, associated either with tumours or healthy tissues (**Table S3**). To provide an interpretation of the built classification models, lipids were analysed by MS/MS. In negative ion mode, out of the 32 discriminants lipids, 24 ions exhibit an increased expression in tumour tissue, primarily consisting of phosphatidylserines (PS), phosphatidylethanolamines (PE), and ceramides. This is notably observed for *m/z* 698.55 Cer (d18:2_24:0 (2OH)), *m/z* 718.55 (PE 34:0), and *m/z* 788.55 (PS (18:1_18:0)) (**Fig. 2F**). In the MS positive ion mode, of the 28 reliable lipids, different phospholipids are present in different abundances in cancer or healthy tissues. Indeed, the ions *m/z* 712.65 to Cer (t20:0_24:0 (2OH)), *m/z* 724.55 (PC 32:5) and *m/z* 734.55 (PE 36:7) are highly expressed in cancerous tissues while the ions at *m/z* 744.55 (PE 36:2) and m/z 766.55 (PC O-36:5) are overexpressed in healthy tissue (**Fig. 2G**). In contrast, for healthy tissues, eight ions exhibit specificity, including certain phosphatidylinositols (PI) and phosphatidic acids (PA). For instance, the PI (18:1_16:0) at *m/z* 835.55 and the lipid PA (18:1_18:2) stand out, with the latter being notably specific in its negatively protonated form (*m/z* 697.45) and in its chloride-adduct form (*m/z* 733.45). These markers can also be confirmed by locating the ions in a mixed EC tissue using SpiderMass imaging. As illustrated in **Fig. 2E**, the distribution of these selected ions well delineates the healthy and cancerous regions within the tissue sample, thereby affirming the reliability of these ions as confident biomarkers.

It was also possible to identify with confidence lipid markers of both non-PCC and PCC subtypes, some that were already found as specific of cancerous tissue in the first model but also new ones. In negative mode, 19 reliable biomarkers were found, with some over- expressed in non-PCC subtype cancer tissues, such as the ions at *m/z* 671.45 corresponding to PA 34:2, *m/z* 673.45 identified as PA 34:1 and *m/z* 762.55 as PE (38:6); and others in PCC subtype cancer tissues, as the ions *m/z* 678.45 attributed to PE 32:6 and *m/z* 687.55 to PA O- 36:1 (**Fig. 2H and Table S3**). Of note, some markers are cross-correlated between the 2-class model cancer versus non-cancer and the 3-class model including the subtyping. This is the case for the ions at *m/z* 697.45 identified as PA (18:1_18:2) and *m/z* 835.55 identified as PI (16:0_18:1), which are found to be markers of healthy tissues as well as non-PCC subtypes. Interestingly, some molecular markers are shared between the healthy and the non-PCC subtypes which is less aggressive and has better survival. Of particular interest is the *m/z* 819.55 ion specific for cancerous tissues, which we also know from previous experiments to be a marker of M2-type macrophages [28] and is over-expressed in PCC tissues. This links the aggressive PCC subtype to M2-type macrophages, which have an anti-inflammatory and pro-tumour profile [35]. Similarly, the lymphocyte-specific ion *m/z* 738.55 corresponding to a PE (16:0_20:4) and the M1 macrophage-specific ion *m/z* 762.55 to a PE 38:6 [28] were found to be over-expressed in non-PCC subtype cancer tissues. In the positive mode, 18 reliable lipids are identified for the discrimination. Some ions, such as the *m/z* 724.55 identified as the PC 32:5 and the *m/z* 780.55 corresponding to PC 36:5 are over-expressed in PCC tissues while the non-PCC subtype is characterised by the presence of ceramides such as the ion *m/z* 650.65 (Cer 42:1) and PEs (e.g.; *m/z* 702.55 PE O-34:2 and *m/z* 792.55 PE 40:6) (**Fig. 2I and Table S3**).

### Margin delineation based on supervised classification model

To leverage SpiderMass as a margin delineation tool, we tested the technology on a pilot cohort of 5 mixed esogastric tissues containing healthy (peritumoral) and cancerous regions (**Fig. 3A -S7**). To simulate the process of intraoperative margin delineation, a single line of analysis traversing all areas across the tissue was performed (**Fig. 3B**). For the interpretation, we used the 2-class models previously built and validated. The spot size analysed by the system is 500 µm in diameter, corresponding to a spatial resolution in good adequation with the needs of the surgeons. Thus, each analytical spot contains slightly more than 1000 cells. As we expect some areas to be constituted by a mixed population of cancer and non-cancer cells, we need an approach that goes beyond binary classification, providing insights into the proportion of healthy and cancerous cells at the level of each spot. This was achieved by incorporating the probabilities calculated by the model for predicting each class at each spot, providing more detailed insights into how the tissue types are classified in a binary classification framework. As shown in **Fig. 3C-E**, a binary classification is insufficient to depict the reality of the tissue margins, considering that surgeons are looking for the “R0” status. In a binary classification, many spots containing a high percentage of cancer cells (e.g., more than 25%) could fall in the class healthy, leading to a false negative result. Results obtained for tissues 68-2 and 66-2 seem to agree with the histopathological annotations, with a margin starting at shot n°7 and n°3 respectively (**Fig. S7**). Tissue 8-2 exemplifies the margin delineation capability of SpiderMass, particularly within complex tissues. In this tissue, a cancerous zone is situated centrally, surrounded by healthy mucosa. Through percentage classification, we well discern two distinct margins within this tissue while the central shot is entirely predicted as cancerous and the shots at the ends are predicted as healthy. Histopathological annotation of tissue 82 suggests 2 different regions with a clear delineation between healthy and cancer almost in the middle. Binary classification alone already compromises the precise identification of margin boundaries, and the percentage-based classification further validates this concern. For instance, in shot n°3, initially deemed healthy based on histopathological annotation, our analysis reveals a composition of 55% cancerous cells. This finding suggests that despite the histopathological classification, this area may warrant consideration within the surgeon’s resection zone. It was validated by MALDI-MSI segmentation, in which we were able to see that the delimitation between the healthy and cancerous zones is not as clear-cut as pathologists describe it (**Fig. 3F-G**). Tissue 103 serves as a compelling demonstration of SpiderMass margin delineation accuracy. In this tissue, the shots n°3 and 4 are predicted to be located at the margin and not in a fully healthy part. Clearly meaningful when observing the MALDI-MSI segmentation, where the margin is defined as higher compared to the histopathological annotation (**Fig. 3F-G**). All these results were confirmed by t-SNE analysis and by MALDI-IHC of Ki67 (**Fig. 3H-I**). In addition, as a negative control, the same procedure was also performed on a two cancerous tissue samples (n°75 and n°112) and two healthy tissue samples (n°74 and n°49) not used in the training of the 2-class model (**Fig. 3 -S7**). As expected, the model classifies the various shots as either healthy or cancerous, excluding margin regions. Specifically, each cancerous spot is classified with no more than 4% of healthy class (with a mean of 98.6% cancer), while healthy spots contain no more than 24% cancerous class (with a mean of 86% healthy). By predicting the ratio of the cancer versus the non-cancer cells this methodology ensures to get a more accurate view on the tissues and the margins and demonstrate that some tissue annotated as healthy by the pathologist do not present molecular fingerprint of completely healthy tissue though. Hence, it is clear that the molecular signature of cancerous tissue must include highly specific markers, as evidenced by the fact that cancerous samples show less than 4% healthy class. In contrast, the molecular signature of healthy tissue seems to be influenced by variations in molecular expression, which explains the relatively high percentages of cancer detected (ranging from at least 2% to approximately 24%). Additionally, the healthy tissues used in this study come from cancerous patients, which may not be as "healthy" as tissue from individuals without cancer. To delve deeper into the analysis, we examined the margin for the presence of specific ions. Interestingly, the margin did not contain unique ions of its own. Instead, we observed a reduction in the expression of ions that were found specific to healthy and cancerous tissues (**Fig. 3J-K -S8-S9**). For instance, ion *m/z* 733.45, which is overexpressed in healthy tissue, is less abundant in the margin region but still more prevalent there than in the cancerous zone. Similarly, ion *m/z* 796.55, predominantly overexpressed in cancerous tissue, shows a gradual reduction in expression across the transition zones: from the cancerous zone to the margin area adjacent to the cancerous tissue (Peri-cancer 50-75), then to the margin near the healthy tissue (Peri-healthy 25-50), and finally to the healthy zone. Thus, SpiderMass identified regions of subtle infiltration, validated independently through MALDI-MS imaging and Ki-67 immunohistochemical staining, confirming its precision and reliability. This percentage-based classification offers surgeons nuanced, actionable insights into margin status, thus potentially reducing recurrence. These different examples show the potential of SpiderMass, an ambient mass spectrometry source, as a future promising tool for real-time margin delineation within the operating room environment. Its ability to accurately discern tissue compositions, even within complex structures, suggests its viability as an asset for improving surgical outcomes through precise margin identification during procedures.

**Figure 3.**
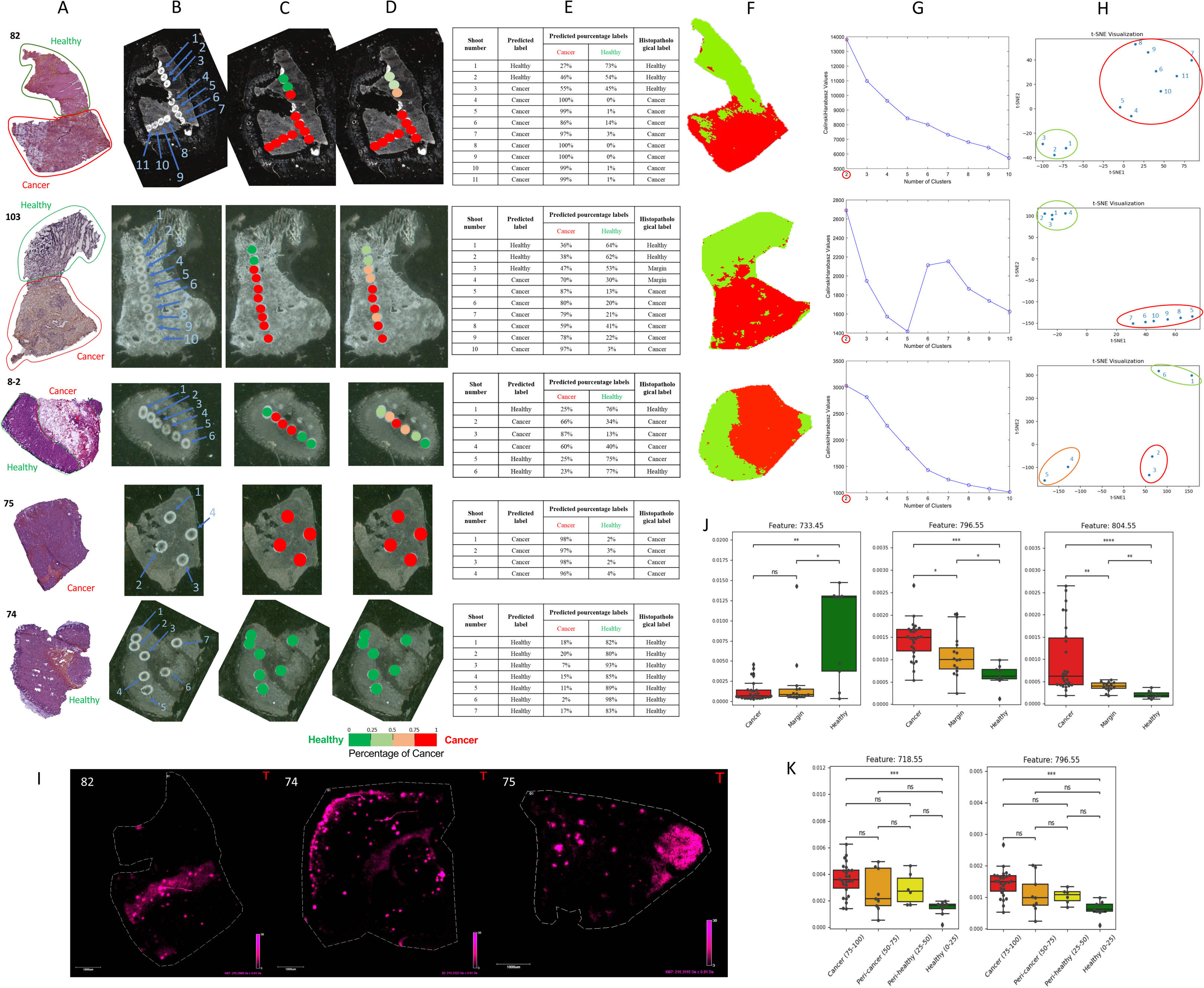
Margin delineation using SpiderMass technology of 3 mixed tissues and 2 negative controls. **(A)** Histopathological annotations based on H&E staining. **(B)** Localization and number of SpiderMass shots, along all the tissue. **(C-D)** Corresponding predicted labels of each shot thanks to two colour-coded legends: binary and percentages, respectively. **(E)** Summarise of all results obtained for each tissue, allowing the comparison of the different techniques. **(F-G)** MALDI-MSI segmentation (2 clusters) of 3 mixte tissues with their corresponding Silhouette plot. **(H)** t-SNE analysis for margin delineation of mixed tissues. **(I)** 1-plex MALDI-IHC on Ki67 biomarker in three tissue sections (left to right: mixte, healthy and cancerous tissues). The display scale (arbitrary peak intensity units) is as follows (minimum intensity/full intensity threshold): 3/30 (Ki67). **(J)** Boxplots of 3 ions overexpressed in healthy or cancerous tissue but here between healthy, margin and cancerous areas. **(K)** Boxplots of 2 ions overexpressed in cancerous tissue but here between healthy, margin and cancerous areas. Related to Fig. S7-S8-S9.

### Prediction of patient outcome by SpiderMass

Using SpiderMass lipid fingerprints, we were also interested in investigating his potential to predict the patient outcome. First, a classification model was built, separating our cohort in < or > 2 years of overall survival (OS). Both in positive and negative ion mode, a correct classification rate of 77% and 76% after 20-fold CV were obtained, respectively. Furthermore, 89% and 74% accuracies were obtained for the blind set (**Fig. 4A**). When examining confident markers to discriminate the OS, 11 ions and 13 ions were identified as indicative of shorter or longer survival, respectively, in positive and negative ion modes (**Fig. 4B** and **Table S4**). Of the identified lipids, many PEs were found to exhibit an overexpression in tissues from patients with an overall survival exceeding 2 years, including ions with *m/z* 736.55 (PE 36:5) and *m/z* 764.55 (PE 38:5). In contrast, PAs showed higher expression in tissues from patients with an overall survival of less than 2 years, as exemplified by the ion *m/z* 701.55 (PA 36:1). We then studied the association of SpiderMass molecular data with the rest of the clinical data gathered for all the patient cohort. For clinical data, the survival curves (Kaplan-Meier analysis) correlate the OS with different factors but mainly as known for esogastric cancer patients with the age, tissue type (PCC vs. non-PCC) and stage pTNM (**Fig. 4C**). For instance, PCC patients exhibit shorter survival, while those under the age of 70 tend to live longer. On the other hand, factors like ASA score, BMI, gastric reflux, neoadjuvant/adjuvant use, gender, tumour location and recurrence show no significant associations (**Fig. S10**). To deepen the analysis, a Cox proportional hazards model incorporating all available metadata was then constructed, complementing the Kaplan-Meier approach, which is limited to two-group comparisons. Several variables, including BMI, age, tissue type and pTNM stage, exhibited p-values < 0.02, indicating their statistical significance to survival (**Fig. 4D -S10-S11A**). For example, a one-level increase in stage pTNM is associated with a 57% increase in the risk of death and a one-year increase in age is associated with a 4% increase in risk. More interestingly, when combining these metadata with SpiderMass lipid markers (found previously as discriminant based on the classification model), the model highlights the importance, not only of the clinical factors but of the specific biomarkers as well, in the patient OS. In the negative ion mode, three ions are related to a negative impact on the OS while two others are associated with long survival (**Fig. 4E -S11B**). Indeed, PE_40:5 (*m/z* 792.55) and PE_36:5 (*m/z* 736.55) are identified as good prognosis factors. PE_40:5, with a hazard ratio of 0.02 (p < 0.005), corresponds to a 98% decrease in mortality risk, while PE_36:5, with a hazard ratio of 0.19 (p = 0.05), is linked to an 81% decrease in mortality risk. Contrarily, PA 36:1 (HR = 22.58, p = 0.003), PE 36:1 (HR = 11.14, p = 0.05), and PE 38:4 (HR = 9.07, p < 0.005) are associated with poorer outcomes, each linked to a substantial increase in the risk of death. These findings are in line with those already found using the classification model distinguishing between patients with survival of < or > 2 years, except for the PE 38:4 marker. In positive ion mode, only one biomarker was found to be associated to the OS. Indeed, PA 44:6 (*m/z* 805.55), with a hazard ratio of 664.97 (p = 0.01), is associated with a dramatic increase in the risk of death, suggesting that higher levels of PA 44:6 are linked to an approximately 99.85% increase in the risk of mortality (**Fig. 4F -S11C**). These combined analyses (both clinical and lipids data) confirmed the discovery of pertinent and reliable lipid biomarkers associated with poorer or improved survival. Thus, Incorporating SpiderMass-derived lipid biomarkers with conventional clinical parameters significantly enhanced prognostic predictions. Patients exhibiting adverse lipid profiles might benefit from intensified postoperative surveillance and adjuvant therapies, highlighting the potential clinical impact of molecular profiling in personalized treatment strategies.

**Figure 4.**
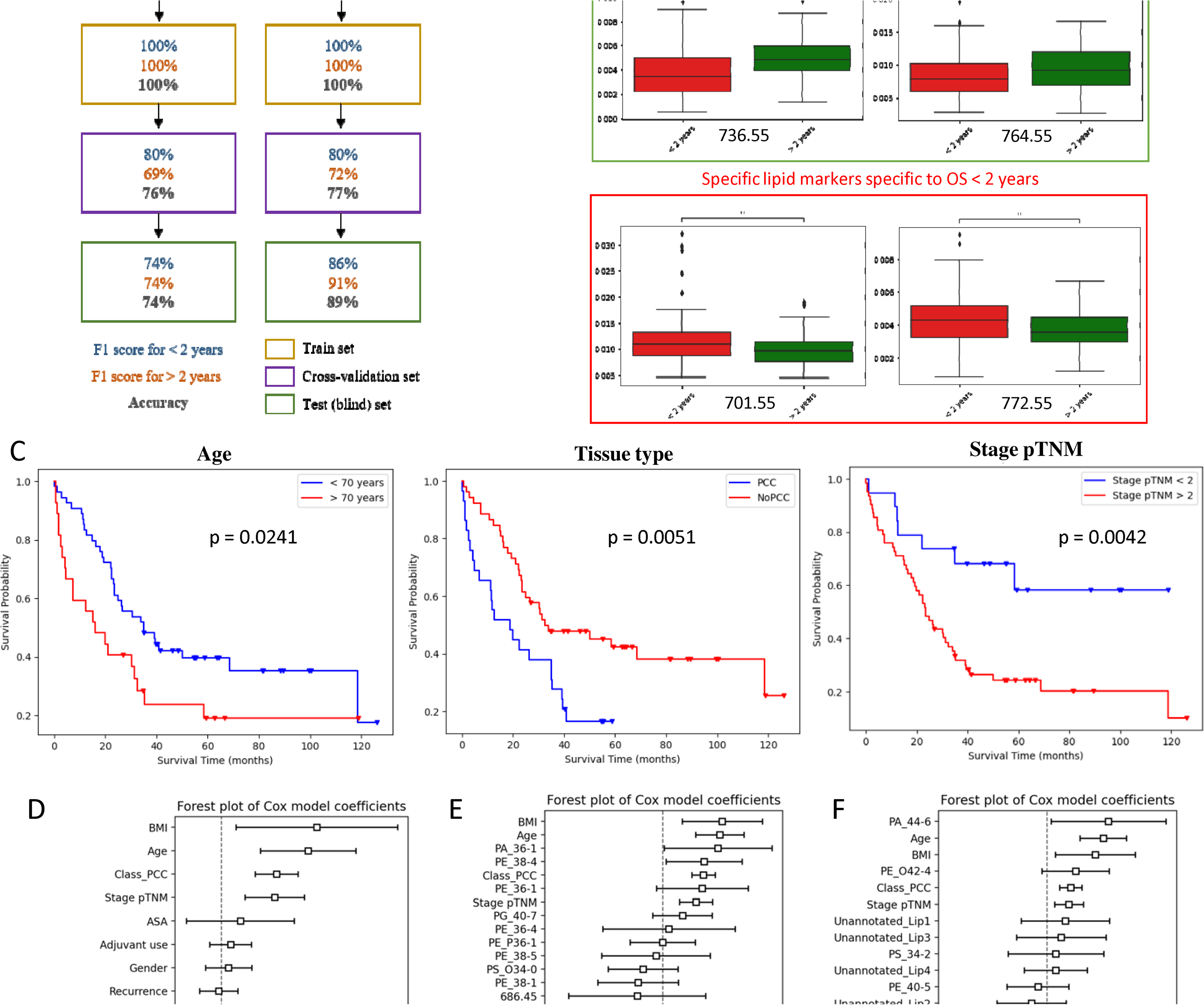
Prediction of patient prognosis based on SpiderMass lipid fingerprints. (A) Schematic report of training, cross-validation and blind test of the classification model for overall survival in negative and positive ion mode. (B) Corresponding boxplot of 4 confident biomarkers specific to OS < or > 2 years. (C) Survival curves (Kaplan Meier analysis) of all patients according to age, tissue type and Stage pTNM. (D-E-F) Forest plot of Cox model coefficients made with all clinical data available, by adding lipid biomarkers in negative mode and in positive ion mode respectively. Related to Table S4 and Fig. S10-S11.

### Diagnosis and prognosis of esogastric cancer tissues by SpiderMass bacterioscoring

#### Lipids MS spectra classification of bacterial strains

We first tested the ability of the SpiderMass technology to discriminate between different bacteria previously shown to the associated with EC. We thus analysed with the SpiderMass technology, directly from Petri dish after one-day incubation, different bacterial strains including *E. coli*, *L. lactis*, *L. gasseri*, *S. anginosus*, *S. salivarius* and *S. bovis* to search for lipids specific to each of these bacterial strains. We also incorporated data from the study by Chen *et al*. [36], which analyzed a significantly larger number of bacterial strains. In addition to the 6 bacterial strains in our original dataset, we included 46 additional strains of interest, resulting in a total of 52 bacterial strains (**Fig. S12 and Supplementary data 1**). All these bacterial strains were very well differentiated by their lipid molecular profiles using a supervised multivariate statistical analysis. Indeed, the built classification model using LinearSVC reached 94% accuracy after 20-fold CV in negative ion mode. An even better classification model was obtained when applying oversampling, using SMOTE algorithm, to the dataset to equilibrate the classes. Indeed, we obtained a model with 96% of accuracy with Random Forest after CV (**Fig. 5A - S13**). When using supervised and unsupervised marker discovery, 82 lipid biomarkers were found in total specifics to a particular species, phylum, genus or order (**Fig. 5B -S14 and Supplementary data 2**).

**Figure 5.**
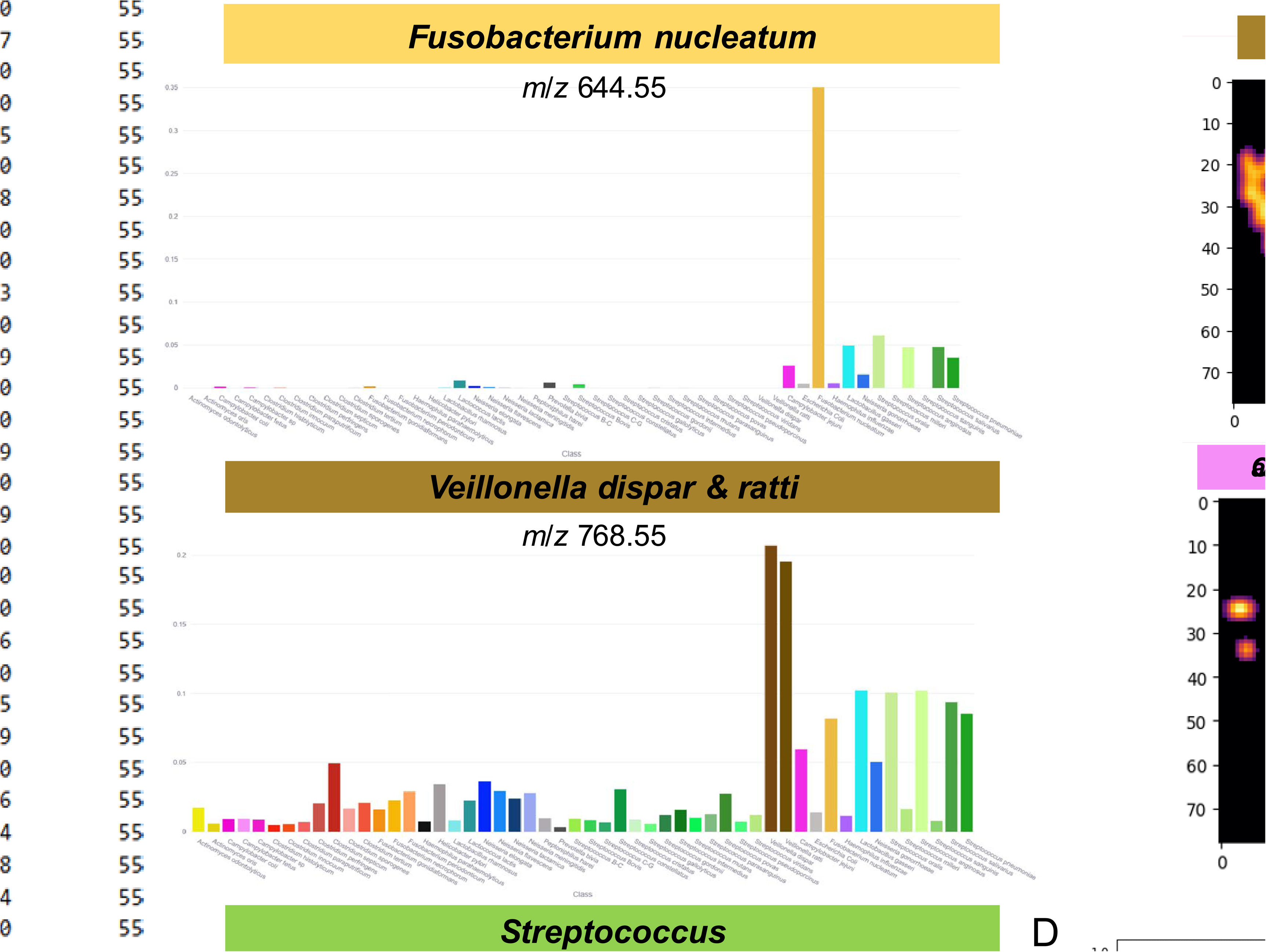
Development of the bacterioscore. **(A)** Classification report of bacterial strains models after oversampling, after 20-fold cross-validation in negative ion mode. **(B)** Mean intensity of 4 lipid biomarkers examples in all 52 bacterial strains. **(C)** TIC Map and bacterioscore images of 5 bacterial strain in one tumoral tissue example. **(D)** Normalized Log abundance Sum of each bacterial strain in one tumoral tissue example. Related to Fig. S12-13-14-15-16.

### Correlating EC subtypes with SpiderMass bacterioscores at the image level

We also investigated the potential of identifying bacterial populations within EC tissues as a feature for intraoperative diagnosis or prognosis assessment. Specifically, we aimed to analyze how a bacterial index, called ‘bacterioscore’ and based on the data generated by analysing the individual bacterial strains, reflect bacterial distribution across different tissue types. The dataset included 13 EC tissues, among which eight were cancerous, distributed over 4 PCCs and 4 non-PCCs. The bacterioscoring model, trained using Random Forest (96% of accuracy), was applied to this 13 SpiderMass images to estimate the confident scores of presences for each bacterial strain by predicting for each pixel the percentage of ratio of each of the bacteria. The model was calibrated using isotonic regression, ensuring that predicted probabilities better reflect true bacterial presence. To enhance interpretability, the scores were transformed into log-abundance values, providing a more nuanced representation of bacterial presence across tissue sections. The mean abundance was then computed for each image, revealing significant differences between cancerous and non-cancerous tissues (**Fig 5C-D**). Out of the 52 bacterial strains, 45 were detected in the tissues. First, using a Venn diagram, certain bacterial strains were identified as being present exclusively in specific tissue types— for example, *Streptococcus viridans* in non-PCC tissues, and *Peptoniphilus harei* and *Clostridium sporogenes* in healthy tissues (**Fig. 6A**). However, these strains were not significantly associated with their respective tissue types, as they were detected in only one out of the four or five samples per group. To compare more precisely these predicted abundances between patients, the mean abundances of detected bacteria in samples were compared using heatmap clustering with an ANOVA test. Using this strategy, we identified two bacterial strains that are overexpressed in healthy tissues with a p-value of 0.01, as shown in **Fig. 6B-D -S15A**. *L. gasseri* appears to be highly specific to healthy tissues (with a median of 14%, in contrast to cancerous tissue with 6% abundance), while *C. fetus* is overexpressed in healthy tissues but also present, albeit minimally, in non-PCC tissues and absent in PCC tissues (with half as much, 45%, in PCC tissues as in the others, 90%). When a lower p-value of 0.05 is applied, three additional bacterial strains emerge as noteworthy. One of them is *F. gonidiaformans*, which shows a similar expression pattern to *C. fetus* in both healthy and non-PCC tissues. In contrast, two bacterial strains from the Neisseria genus (*N. elongata* and *N. meningitidis*) are overexpressed in cancerous tissues, particularly in PCC tissues (with an abundance of 26% and only 2% for healthy tissues) (**Fig. 6C-D -S15B**). It was confirmed by looking at the bacterioscores examples shown in **Fig. 6F**. Interestingly, we discovered that the ratio of *L. gasseri* to *N. elongata* holds potential as a diagnosis marker. Specifically, a *L. gasseri*/*N. elongata* ratio below 0.2 is strongly associated with cancerous tissue, while a ratio exceeding 0.4 suggests healthy tissue. Notably, when this ratio is particularly low (approaching 0), it may serve as an accurate predictor of the presence of a PCC subtype (**Fig. 6E**). In addition, although not statistically significant, the expression of various bacterial strains across different tissue types revealed some interesting patterns. For instance, *A. odontolitycus* and *L. lactis* were overexpressed in cancerous tissues, while *F. periodonticum* showed higher expression specifically in PCC tissues (**Fig. S15C-D-E-F**).

**Figure 6.**
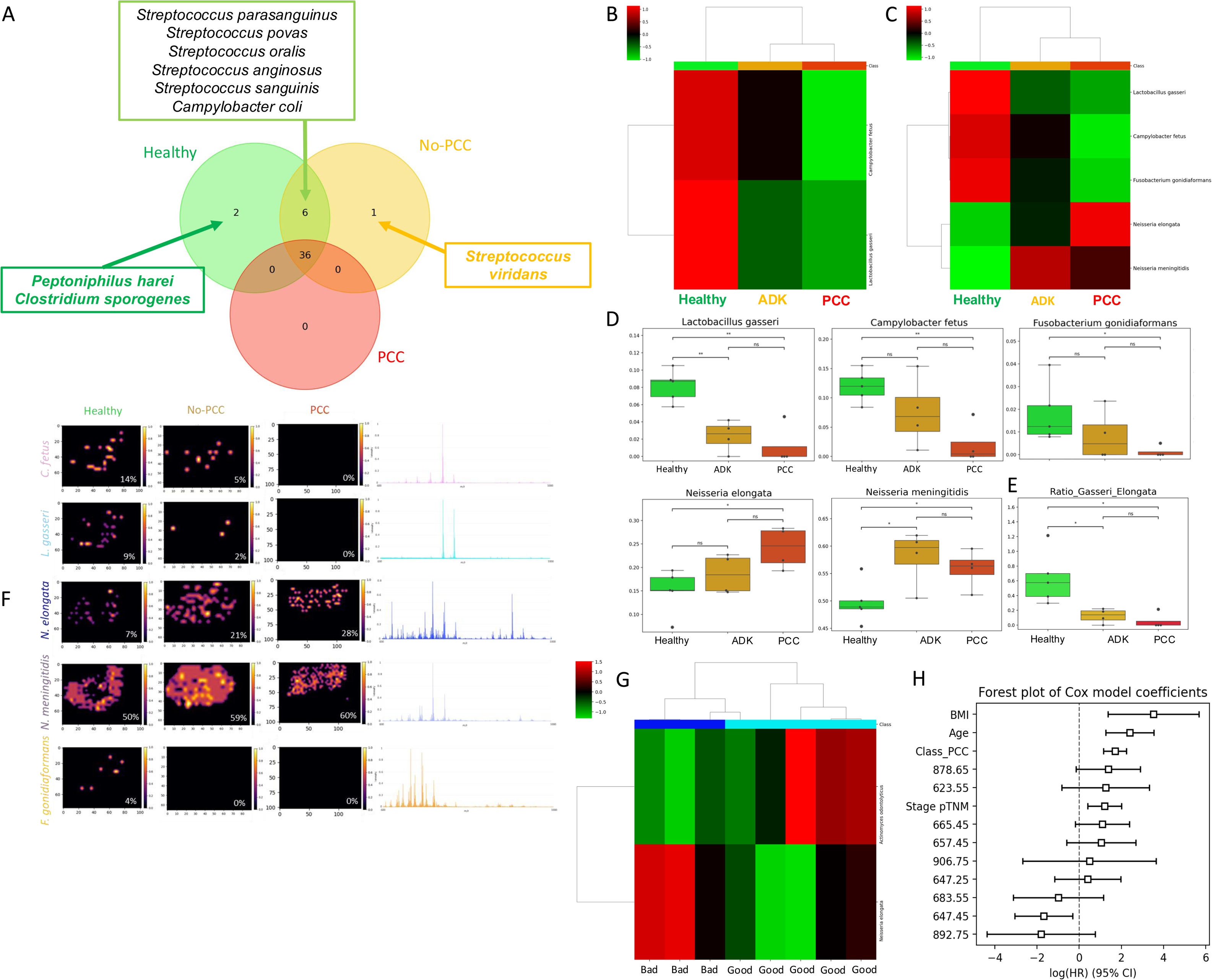
Diagnosis and prognosis of esogastric cancer tissues based on SpiderMass signature of bacterial strains. (**A**) Diagram de Venn. **(B-C)** Heatmap following ANOVA analysis of average tissues with a p-value of 0.01 and 0.05 respectively. **(D)** Corresponding boxplot of 5 bacterial strains specific of either cancerous tissues or healthy. **(E)** *L. gasseri**/**N. elongata* ratio boxplot in different type of EC tissues. **(F)** Bacterioscoring representation of 5 interesting bacterial strains in 3 fresh frozen tissues examples of each type; healthy, PCC and no-PCC. For each bacterial strain, the abundance in percentage is indicated and their corresponding mean spectra was added. **(G)** Heatmap following ANOVA analysis of bad prognosis vs good prognosis. **(H)** Forest plot of Cox model coefficients made with combined significant clinical data and interesting bacteria strains lipid biomarkers. Related to Fig. S12- 13-14-15-16.

#### Correlating EC prognosis with SpiderMass image bacterioscoring

Since we demonstrated that certain bacteria were differentially abundant according to the type of tissues, we also wanted to examine the hypothesis that certain bacteria could be linked to the OS. We performed the bacterioscore from SpiderMass images obtained from the tissues of patients with different OS (two patients with a bad prognosis lower than 1 year and four patients with a good prognosis over 3 years). The bacterioscore calculated at the image level shows a correlation to the OS for only two bacterial strains, *N. elongata* and *A. odontolitycus*. *N. elongata* is overexpressed in tissues from patients with poor survival (less than one year), whereas *A. odontolyticus* is more abundantly present in tissues from patients with longer survival (**Fig. 6G**). These findings provide direct evidence of the possibility to use microbiota analysis, through *in vivo* bacterioscoring as a predictive feature for patients with EC. Furthermore, using a Cox proportional hazards model combining clinical data and marker ions for bacteria in negative ion mode from all the patient data, it was also shown that two ions are strongly correlated with the patient OS. Indeed, *m/z* 878.65, specific to *N. elongata*, exerts a significant positive effect on risk (exp(coef) = 4, p = 0.07), associated with a significant decrease in the OS. In contrast, one other ion, *m/z* 647.45, specific to *F. gonidiaformans*, show a marked negative effect on risk (p=0.01), indicating a protective effect that allows for increased patient survival (**Fig. 6H -S16**).

Thus, we constructed a bacterial spectral library representing gastric microbiota and developed a bacterioscore quantifying microbial signatures directly from tissue spectra. The ratio of *Lactobacillus gasseri* to *Neisseria elongata* emerged as a powerful indicator distinguishing healthy tissues from tumors, especially aggressive PCC. However, patient-specific factors such as recent antibiotic use represent confounders necessitating controlled future validations to confirm clinical utility. We confirm in the present study, that PPI has no effect on the diagnosis and prognosis based on the integration of SpiderMass data with clinical data.

## Discussion

This study presents an in-depth investigation into the possibility of improving the management of patients with EC and reducing relapse through more accurate surgery using the MS-based SpiderMass technology. The versatility of SpiderMass was demonstrated by its ability to perform accurate histological classification, cancer subtyping, and bacterioscoring in the context of EC.

Firstly, the optimal classification model for discriminating cancerous tissue from healthy tissue achieved accuracy rates of 93% and 90%, respectively, using negative and positive ionisation modes. In blind predictions performed on an independent validation cohort of 32 samples, only 3 samples were misclassified in the negative mode, equivalent to a 91% correct classification rate. A three-class model distinguishing between healthy, PCC, and non-PCC samples was trained, achieving strong classification rates of 93% in negative mode and 96% in positive mode. When applied to a blinded dataset, the model demonstrated a robust classification accuracy of 84%. In addition, we were further able to clearly distinguish the two ADK subtypes, PCC and non-PCC, using a two-class classification model that achieved an accuracy rate of 98% and 97% in each ionization mode. However, this blind model was only able to classify 76% of tissues correctly. Both strategies highlight the complexity of *in vivo* identification of EC subtypes.

To provide a biological meaning and give an interpretation to our molecular-based histological classification, we achieved the identification of the lipid markers. Variations in the phospholipid classes were observed depending on the tissue type (cancer versus healthy) and subtypes (PCCS versus non-PCCs). Healthy tissues are characterised by the presence of PIs, PAs and PCs, while cancer tissues show a significant concentration of DGs, PSs, PEs and Cer. The overexpression of PEs in cancer tissue is intriguing due to findings by Luo *et al*. [37] indicating the significance of the PE domain within the PEBP4 protein. Their results propose that this domain plays a crucial role, as PEBP4 can potentially boost cancer cell resistance to apoptosis by hindering the activation of ERK signalling pathways. In addition, increased PS exposure of tumour cells in the TME promotes innate immunosuppression, aiding tumour growth and metastasis [38]. Current research explores agents targeting PS, some undergoing clinical trials as combination therapies for diverse cancers. Interestingly, some molecular markers are shared between healthy and non-PCC tissues, non-PCC being less aggressive than PCC and associated with better survival. In addition, specific ions to lymphocyte and M1 macrophage cell populations are over-expressed in non-PCC tissues.; while one ion specific to M2-type macrophages is over-expressed in PCC tissues. This finding is highly interesting because it links M2-type macrophages which have anti-inflammatory properties and promote tumour growth (unlike M1-type macrophages), to the particularly aggressive PCC subtype. These results suggest significant differences between the two cancer subtypes in terms of molecular markers and the involvement of specific immune cells in tumour progression.

In addition, SpiderMass technology was applied to mixed cases of EC tissues, presenting both a tumoral and peritumoral region, to evaluate its potential for real-time margin delineation during surgery. A percentage-based classification was developed for margin delineation because the analysed area comprises a mixed population of tumoral, healthy cells and other cell types, such as immune cells. Indeed, the 2-class cancer vs. non-cancer classification model cannot account for the complexity of the tumour boundary as demonstrated by the calculated percentages of tumour versus non-tumour cell populations. This method, which integrates for more precise boundary identification, demonstrated strong agreement with histopathological annotations and highlighted its ability to refine margin delineation, even in complex tissues, confirmed also by t-SNE analysis. Some discrepancies between our results and histopathological findings underscored the strength of SpiderMass as a margin delineation tool, given its high concordance with MALDI-MSI segmentation outcomes.

SpiderMass lipid molecular fingerprints were also tested against patient outcome prediction. The built classification model showed 77% and 76% accuracies in cross-validation for overall survival (< or > 2 years), in positive and negative ion modes, respectively. Key lipid markers, such as PE 40:5 (correlated with better outcomes, HR = 0.02) and PA 36:1 (risk factor, HR = 22.58), were identified alongside clinical factors like age and pTNM stage as significant predictors of survival. Combining metadata with SpiderMass biomarkers revealed insights into the interplay between clinical and molecular factors, highlighting potential targets for further research to improve patient therapy.

Another meaningful aspect of this study is the use of SpiderMass to diagnose EC based on the presence of certain bacterial strains. We tested 52 bacteria strains known to be present in the esogastric area. The individually grown bacterial strains were analysed by SpiderMass or REIMS, in this study [36], to get their specific lipid molecular profiles and use these profiles to predict their presence in the EC tissues. Currently, pathologists are limited to detecting *Helicobacter pylori*, overlooking other potentially relevant bacteria, from preserved tissue samples, but this detection is only possible after surgery. This highlights the need for earlier and more comprehensive methods for bacterial detection in EC. The creation of a bacterioscore adapted to the SpiderMass approach could make it possible to translate the presence of bacteria *in vivo* and predict the results during surgery, which could help surgeons personalise patient management and shift to precision surgery. Thanks to this approach we could demonstrate that *L. gasseri* is less abundant in cancer tissues, particularly in PCCs, than in healthy tissues, where it is predominant. Numerous studies have demonstrated the close link between *L. gasseri* and *H. pylori*. About half the world’s population is infected with *H. pylori*, an infection causing inflammation of the gastric mucosa and gradually leading to the disappearance of the parietal cells in the stomach, responsible for secreting hydrochloric acid. This ultimately leads to a condition known as atrophic gastritis, characterised by chronic inflammation and a reduction in gastric acidity, thus considerably increasing the risk of developing gastric cancer [39]. However, it is important to note that *L. gasseri*, one of the most renowned probiotics, appears effective against *H. pylori*. The main antibacterial mechanisms of probiotics include competition with *H. pylori* for binding sites on gastric epithelial cells, reinforcement of the mucosal barrier and production of organic acids with bactericidal properties [40]. In vitro studies revealed that *L. gasseri* caused a significant decrease in the expression of certain oncogenes, including Bcl-2, β-catenin, integrin α5 and integrin β1. Overall, current evidence suggests that certain strains of *Lactobacillus* may be beneficial to human gastric health due to their ability to eliminate pathogens, maintain gastric barrier integrity, reduce inflammation and exert protective effects against cancer [41,42]. However, the opposite trend was observed for two *Neisseria* strains, with a higher presence in cancer than in healthy tissues. Indeed, it was already shown that several Neisseria species were enriched in gastric cancer and are known as oral cavity commensals and opportunistic pathogens [43]. These species are associated with pathways related to LPS and ubiquinol biosynthesis, which promote inflammation and tumorigenesis, warranting further research on their potential impact on gastric cancer [44]. Finally, the analysis of human microbiota in esogastric tissues, using SpiderMass, reveals intriguing connections with the survival of cancer patients. Examining fresh frozen tissues from individuals without esogastric cancer recurrence, a correlation was observed between bacterial presence and overall patient survival. Specifically, *A. odontolyticus* is predicted to be more abundant in tissues from patients with longer survivals when compared with those who die earlier. The inverse was found for *N. elongata* strain. Such associations underscore the potential of microbiota analysis as a predictive marker for gastric cancer patients, offering insights for personalised treatment strategies. Considering that our study demonstrates that changes in the bacteria populations in the esogastric area could be a predictive factor of cancer developments and patient outcomes, this predictive quantitative approach offers new insights into the microbial landscape of EC tissues, supporting the development of bacterioscoring-based diagnosis and prognosis tools. It is important to note that in our study we had the opportunity to obtain healthy tissue from the same patients, but at a distance from the cancer site. However, it is unclear whether patients have a general dysbiosis of the gut that affects the bacterial microbiota even in distant healthy tissues. In this respect, we would probably measure a more important difference in the bacterial population if we could compare with people without cancer. However, obtaining such tissues is clearly a bottleneck.

Besides, it would be interesting in the future to explore SpiderMass for pre-surgery steps and even during routine diagnosis by endoscopy procedures to evaluate the possibility offered by the technology to detect EC at early stages of developments which would considerably improve the patient outcomes. Additionally, one of the next steps we wish to address would be the demonstration of the technology through a clinical trial intraoperatively on patient-resected tissue first, by placing our SpiderMass prototype in a room next to the operating theatre and then moving the prototype in the surgery room for direct evaluation on patient. In this context, the miniaturisation of the SpiderMass technology for its integration into the endoscope would allow to offer an earlier detection of the microbiota changes as an indicator of the first stage of cancer development. This will be an added value, as unfortunately the detection of EC is at stage III or IV, giving a poor OS at 5 years.

## Acknowledgments

This work was supported by grants from Excellence Initiative of Lille University (ISite ULNE Sustain program, IF), Canceropôle Nord-Ouest (emergence project, GP and IF) and Institut Universitaire de France (IF). L.L. PhD was co-founded by University of Lille Excellence Initiative, Région Haut de France (EU Feder funds) and OCR company. Y.Z. PhD is funded by ANR CE29 2023 “Click&Detect”. FREGAT was funded with support from the Institut National du Cancer (INCa), and CHU de Lille is the sponsor of the FREGAT study. We thank the data management department of the SEED unit from CHU de Lille for the administration of FREGAT database and the provision of data. We thank the FREGAT working group and in particular the team of CHU-Lille: Guillaume Piessen, Anthony Turpin, Emmanuelle Leteurtre and Michael Hisbergues. We thank the Tumorothèque ALLIANCE-CANCER de Lille for the contribution and technical support on the tissue samples.

## Authors contribution

I.F. and M.S. designed the experiments. L.L. performed the experiments. C.L. and A.G. designed the synthesis of the Tags for the MALDI IHC. A.G. performed the synthesis of the Tags and the MALDI-IHC experiments. G.P. and J.V. are promotors of the clinical study and gathered patient clinical data. C.D. and F.R. performed the histopathology examination. Y.Z. designed the data analysis workflow and developed the bacterioscore. I.F., L.L., Y.Z. and M.S. analysed and interpreted the data. L.L. and I.F. wrote the manuscript’s original draft. G.P., I.F., J.V., L.L., M.S. and Y.Z. corrected the manuscript. I.F. supervised the project. I.F., C.L. and G.P. provided the funding.

## Competing interest

L.L., Y.Z., G.P., C.D., F.R., A.G., C.L. and J.V. declare they have no competing interests.

M.S. and I.F. are inventors on a patent (priority number WO2015IB57301 20150922) related to part of the described protocol.

## Material and methods

### Esogastric cancer tissues cohort

Tissue samples and associated data were obtained from the tumour tissue bank “Tumorothèque ALLIANCE-CANCER de Lille” and the French EsoGAstric Tumors, FREGAT Database (fregat-database.org) [45]. Tissue and clinical data were collected in compliance to the French legislation with getting patient informed consent. Patient tissues were snap-frozen and stored at -80°C. 164 fresh-frozen tissue samples from 108 patients, of which 77 are healthy and 84 are adenocarcinomas, with 60 no-PCC and 27 PCC subtypes (**Table S1-S2**).

### Bacterial strains

Several bacterial strains (Escherichia Coli, Lactococcus lactis, Lactobacillus gasseri, Streptococcus bovis, Streptococcus anginosus and Streptococcus salivarius) were obtained from the American Type Culture Collection (ATCC).

### Sample preparation

All the excised tissues were sectioned in a precise order on a Leica CM1510S cryostat (Leica Biosystems, Nanterre, France). Three sections were collected, one (5 µm) for the HPS (Hematoxyline Phloxine Saffron) staining, the second (20 µm) for SpiderMass analysis and the last one (12 µm) for the MALDI-MSI analysis. The different sections were mounted on a regular glass slide and on ITO (Indium Tin Oxide) conductive glass slide respectively.

### Histological staining and annotations

The 5 µm thick tissue slices were further processed for staining. First, the tissue was exposed to Hemalum solution for 1 minute, followed by rinsing using tap water. Subsequently, the tissue section was placed for 10 s in a 0.1% phloxine solution and then gradually dehydrated by 70% and then 100% ethanol bath for 1 min. After staining, the section underwent a cleaning process in xylene, followed by two rounds of alcohol cleaning. It was then briefly immersed in saffron for 5 seconds before being fixed with coverslips and EUKITT slide mounting media. The stained slides were then digitised using the Pannoramic MIDI slide scanner from 3DHISTECH LTD. (Budapest, Hungary). The resulting images were visualised and exported using QuPath 0.2.3. All annotations were made from the stained tissues.

### SpiderMass analysis

The comprehensive configuration of the instrument arrangement has been previously discussed elsewhere [21]. Briefly, the system consists of three main components: a mass spectrometer, a laser system designed for remote micro-sampling of tissues, and a transfer line for transporting the desorbed material up to the mass spectrometer. The laser is an OPO (optical parametric oscillator) system emitting at 2.94 µm with a pulse duration of 6 ns and a repetition rate of 20 Hz. A biocompatible laser fiber, 1-1.5 m in length of 450 µm core diameter, is connected to the laser system output and has a handpiece affixed to its end, including a 4 cm CaF2 focusing lens. The laser intensity was adjusted to 4 mJ per pulse, maintaining a constant irradiation duration of 10 seconds. This yielded a laser fluence of about 3 J/cm². The second key component of the setup is the transfer line made out from a 2- meter Tygon ND 100-65 tubing (sourced from Akron in Ohio, USA), boasting internal dimensions of 2.4 mm diameter and 4 mm outer diameter. This transfer line sucks the aerosol generated by the laser shot, up to the mass spectrometer (Xevo G2-XS, Waters) through a REIMS prototype interface which replaces the conventional ESI source of the instrument. For each acquisition, an infusion of isopropanol at 200 µL/min flow rate was introduced, supplemented by 200 µg/mL of leucine enkephalin to serve as a lock mass. The sampling position was determined based on the histopathological annotations. The data acquisition process encompassed 10 laser shots in a burst each yielding to an individual spectrum. Spectral acquisition was conducted in both positive and negative ion modes, choosing the sensitivity mode of the instrument, with a scan duration of 1 second. The targeted mass range spanned from *m/z* 50 to 2000.

### WALDI-MSI analysis

The SpiderMass setup was described in the previous section. For MS imaging, the Spider- Mass microprobe was coupled to a stiff robotic arm described in detail in a previous work [46]. The spatial step size was set to 250 µm to achieve 250 µm spatial resolution in oversampling. The mass range was fixed between *m/z* 50-2000. The acquisition sequence was composed of 3 consecutive laser shots and 3 seconds between each step. The laser bursts and the spectrometer acquisition were automatically triggered through a MATLAB in-house user interface developed for the robotic WALDI-MSI [46]. The data was acquired in negative and sensitivity ion mode.

### MALDI-MSI analysis

The Norharmane matrix at 7 mg/mL in CHCl_3_:CH_3_OH (2:1, *v/v*), was used for dual polarity imaging. The matrix was sprayed using the automated HTX TM-Sprayer (HTX Technologies, LLC). The parameters were set as follows: 30°C nozzle temperature, 12 passes with a CC pattern, 0.1 mL/min as flow rate, the nozzle velocity at 1200 mm/min, the track width spacing at 2 mm and 10 psi for the gas pressure. The MALDI-MSI analysis was performed in both ion polarities on a MALDI-TOF (Rapiflex TissueTyper, Bruker) using red phosphorus as an external calibrant. MSI data were acquired by using a 50 × 50 μm^2^ raster and a 20 × 20 μm^2^ beam scan area for dual polarity imaging with a mass range of *m/z* 400-1500.

### MALDI-IHC analysis

For MALDI-IHC, 12 µm sections on ITO conductive slides were used. Prior to analysis, FF slides underwent several tissue preparation steps, including: 1-minute thawing in a desiccator, and delipidation to make protein sites more accessible. The delipidation process involved immersing the slides in a series of consecutive baths as follows: 30 seconds in 70% EtOH, 30 seconds in 100% EtOH, 2 minutes in Carnoy’s solution (3:6:1 CHCl_3_:EtOH:Acetic Acid), 30 seconds in 100% EtOH, 30 seconds in H2O, and 30 seconds in 100% EtOH. Tissues were then dried under vacuum. A hydrophobic barrier was drawn around the tissue for the next steps. A blocking buffer (5% serum, 2% BSA, 0.3% Triton X-100 in PBS) was then applied to the tissue for at least 2 hours at room temperature. After the blocking buffer was aspirated, primary antibodies (conjugated with the TAG probe) in the blocking buffer were added to the tissue. After overnight incubation at 4°C in a humid chamber, the antibody solution was removed from the tissue before proceeding with a series of washes. The slides were immersed consecutively in two baths of PBS for 1 minute, followed by three 5-minute baths in milliQ water. The slides were then dried under vacuum for at least 1 hour, followed by UV exposure at a wavelength of 365 nm for 20 minutes to activate the photo-cleavage of the probes. To perform MALDI MSI, 2,5-DHB matrix at 20 mg/mL, dissolved in MeOH:TFA 0.1% (70:30, *v:v*), was sprayed onto the tissues using the HTX M5 automated system. The parameters for matrix deposition included a nebulization temperature of 75°C, a plate temperature of 55°C, a pressure of 10 psi, and a flow rate of 0.1 mL/min for 12 passes. Subsequently, the slides were analyzed using a Rapiflex MALDI TOF mass spectrometer for mass spectrometry imaging. Spectra were acquired in positive reflectron mode, in a *m/z* range 140-400, with 500 laser shots per pixel, providing a spatial resolution of 10 µm and continuous scanning over a 10 µm area. The laser energy was set around 45 % and the voltages of the ion source were 20 kV and 11 kV for the lens. The image data were processed using FlexImaging software, applying total ion count (TIC) normalization.

### Bacterial strain analysis

The bacterial strains used in this study are Escherichia Coli (ATCC 12435), Lactococcus lactis (ATCC 19435), Lactobacillus gasseri (ATCC 33323), Streptococcus bovis (ATCC 33317), Streptococcus anginosus (ATCC 700231) and Streptococcus salivarius (ATCC 7073). All the strains were obtained from the American Type Culture Collection (ATCC). E. coli was cultured on Tryptone and NaCl agar plates, Lactococcus lactis and Streptococcus salivarius were cultured on Brain Heart infusion (BD 211065) agar plates, Streptococcus anginosus and bovis were cultured on Tryptic soy (BD 236950) agar plates and Lactobacillus gasseri was cultured on Lactobacilli MRS (BD 288210) agar plates, at 37°C for 24 hours. The SpiderMass analysis was made directly on the agar plates.

### MS/MS analysis

MS/MS spectra were recorded by subjecting to collision-induced dissociation (CID) the selected precursor ion. The collision energy in the collision cell was ramps between 30 and 40 eV. Lipid annotations were executed manually, adhering to fragmentation spectra guidelines, and were cross-referenced against the LipidMaps database, Alex123, Metlin database, and relevant literature.

### Data processing and analysis

#### SpiderMass data

The SpiderMass data consists of .raw files in Waters format. To read and process these files, a data importation pipeline was developed. This pipeline includes executing the msconvert command from the ProteoWizard software (version 3.0.24) directly within Python using the subprocess python module. The raw data was converted to the mzML format and aligned based on the lock mass (*m/z* 554.2 in negative ion mode and *m/z* 556.2 in positive ion mode). To import the data into Python, the pyopenms package was used. Peak detection was performed using the find_peaks function from the scipy package (version 1.7.3) to extract individual spectra from the chromatogram peaks. Each peak corresponds to a laser shot from SpiderMass, containing spectral information. In the final step, the spectra were structured as a Pandas DataFrame (version 1.4.2) , where the first four columns correspond to metadata: "Class" (The target class of the spectrum) "File" (The file name from which the spectrum originates), "RT" (The retention time in the chromatogram, corresponding to the acquisition time), “Sum” (The total ion count (TIC) registered in the spectrum), The remaining columns correspond to *m/z* feature values, where each row contains intensity values for those *m/z* features. The data was then binned to 0.1 Da to reduce the number of data points, and the mass range was set between 600 and 1100 *m/z*. Finally, total ion current (TIC) normalization was applied to remove intensity variations between spectra. The final dataset contains 5000 *m/z* data points.

#### Classification model based on SpiderMass data and biomarkers discovery

Protocol to analyse 1D-mass spectrometry data [28,34] was used to build, train and test multiple models from scikit-learn (24 classifiers). The optimal model was individually reconstructed in scikit-learn, fine-tuning its parameters. To address class imbalance, the dataset was augmented using SMOTE (Synthetic Minority Over-sampling Technique) from the imbalanced-learn package (version 0.10.1), increasing the total number of spectra to the highest (healthy most of the time). 5 and 20-fold cross-validation was performed, and a classification report was generated. The model was saved using joblib (version 1.4.2) in PKL format for blind predictions. Local Interpretable Model-agnostic Explanations (LIME) [47] explained the model using positive/negative feature contributions, visualized in a LIME table via ELI5 (version 0.13.0). A peak picking algorithm (findpeaks function) from SciPy (version 1.10.1) removed instrument noise, and Kruskal-Wallis with Bonferroni correction assessed feature significance. Only p≤0.05 features were kept, and mono-isotopic peaks were filtered. Box plots were created with seaborn. To identify confident lipid biomarkers, supervised and unsupervised analyses were combined, requiring significance, correct expression, and contribution alignment within an ROI.

#### Margin delineation

For the margin delineation, the previously trained cancer vs. healthy model was used. In this case, the predict_proba function was employed to generate the probability for each class at each spot analyzed by SpiderMass across the tissue. These probabilities were then visualized using a color-coded scheme to represent the percentages.

#### MALDI-MSI segmentation

The raw MALDI MSI data for lipids in both ionization modes were initially converted into the imzML format using SCiLS lab software. Subsequently, the imzML converter, version 1.3.3, was employed to import these datasets into MATLAB R2023b. SVD (Singular Value Decomposition) data compression was implemented as a preprocessing step before segmentation. For the segmentation process, the *k*-means++ algorithm was used, implemented as the ’k-means’ function in the MATLAB Statistics Toolbox. The cosine distance metric was employed to calculate the cosine angle between two spectra for quantifying the similarity. For visualization, each cluster’s pixels are uniformly assigned a specific color, facilitating the creation of a segmentation map. To estimate the right numbers of clusters, the Silhouette criterion was used. The silhouette plot displays a measure of the proximity of each point in a cluster [48]. Silhouette plot was calculated using the function silhouette in Matlab.

#### Bacterioscore

The BacterioScore model was developed using the Random Forest algorithm from the sklearn.ensemble module of the scikit-learn library (version 1.2.2). This model used all 52 bacterial strains spectra within the *m/z* range of 600–1000 in negative ion mode. To address class imbalance, the dataset was augmented using SMOTE (Synthetic Minority Over- sampling Technique) from the imbalanced-learn package (version 0.10.1), increasing the total number of spectra to over 2860 (55 spectra for each bacterial strain). To ensure that the predicted probability scores accurately reflect the true likelihood of bacterial presence, the model was calibrated using isotonic regression. Isotonic regression is a non-parametric method that fits a piecewise constant non-decreasing function to the predicted probabilities, ensuring that they are monotonically increasing with respect to the true probabilities. To assess the generalization of the model and validate its robustness while minimizing overfitting, 20-fold cross-validation was applied. For predicting bacterial presence in SpiderMass images derived from tissue samples, the predict_proba function was used to generate probability estimates at the pixel level in each image. These probability scores were then transformed into log-abundance values to enhance interpretability. The transformed abundances were subsequently scaled to a 0–1 range and encoded into a color scale for visualization, generating detailed bacterial distribution maps within the tissue samples. For inter-patient comparisons, the mean abundance of each bacterial strain was computed from the image data. To enable standardized comparisons across different samples, these abundances were normalized relative to the most abundant bacterial strain within each sample, ensuring consistency in cross-sample analyses. For ease of comparison, heatmap clustering from seaborn package (version 0.11.2) was combined with an Anova test to retain the desired significant differences (eg., p<= 0.05, p<= 0.01, p<=0.001).

#### Kaplan-Meier Survival Analysis

Kaplan-Meier survival analysis was performed using the Kaplan-MeierFitter class from the lifelines library to estimate the survival function of patients. Survival times and event indicators were provided as inputs as well as the related group stratification for each analysis. The Kaplan-Meier curve was plotted using matplotlib.pyplot to visualize the survival probability over time.

#### Cox Proportional Hazards Model

The Cox Proportional Hazards model was applied to assess the effect of various covariates on survival. The model was fitted using the CoxPHFitter class from the lifelines library (version 0.27.8). The proportional hazards assumption was tested using the proportional_hazard_test function from lifelines.statistics. A forest plot of the Cox model coefficients was generated using the plot method of the CoxPHFitter class, with visualization provided by matplotlib.pyplot. The forest plot included the coefficients and their 95% confidence intervals, offering insights into the relative risk of each factor.

## Supplemental figures legends

**Table S1.** Overview of clinical data of the EC cohort. Related to Fig. 2-3-4-5-6.

**Table S2**. Demographic and clinical data for all the EC cohort (169 tissues). Related to Fig. 2- 3-4-5-6.

**Figure S1.** H&E scans of the 27 PCC esogastric tissues from the total EC prospective cohort. Related to Fig. 2.

**Figure S2.** H&E scans of the 60 non-PCC esogastric tissues from the total EC prospective cohort. Related to Fig. 2.

**Figure S3.** H&E scans of the 77 healthy esogastric tissues from the total EC prospective cohort. Related to Fig. 2.

**Figure S4.** (**A**) Classification report and confusion matrix after 5- and 20-fold cross-validation of the classification model built from the negative mode and positive mode. SpiderMass using the RidgeClassifier algorithm to discriminate cancer from healthy tissues from 132 tissues of the cohort. (**B**) Results from the blind interrogation of this model based on the 32 remaining tissues of the cohort (summary table and classification report). Related to Fig. 2.

**Figure S5.** (**A**) Classification report and confusion matrix after 5- and 20-fold cross-validation of the classification model built from the negative mode and positive mode. SpiderMass using the RidgeClassifier algorithm to discriminate PCC vs non-PCC vs healthy from 132 tissues of the cohort. (**B**) Results from the blind interrogation of this model based on the 32 remaining tissues of the cohort (summary table and classification report). Related to Fig. 2.

**Figure S6.** (**A**) Classification report and confusion matrix after 5- and 20-fold cross-validation of the classification model built from the negative mode and positive mode. SpiderMass using the RidgeClassifier algorithm to discriminate PCC vs the other subtypes of GC (non-PCCs) from 132 tissues of the cohort. (**B**) Results from the blind interrogation of this model based on the 32 remaining tissues of the cohort (summary table and classification report). Related to Fig. 2.

**Table S3.** Comprehensive list of reliable biomarkers for healthy and cancerous tissues, as well as for PCC and no-PCC cancer subtypes in both ion modes. Related to Fig. 2.

**Figure S7.** Margin delineation using SpiderMass technology of 4 additional mixte, healthy and cancerous tissues. Related to Fig. 3.

**Figure S8.** Boxplots of 20 ions overexpressed in healthy or cancerous tissue but here between healthy, margin and cancerous areas. Their putative annotations can be found in the Table S3. Related to Fig. 3.

**Figure S9.** Boxplots of 4 ions overexpressed in cancerous tissue but here between healthy, peri-healthy margin, peri-cancer margin and cancerous areas. Their putative annotations can be found in the Table S3. Related to Fig. 3.

**Table S4.** Comprehensive list of reliable biomarkers for the global survival in both ion modes. Related to Fig. 4.

**Figure S10.** Survival curves (Kaplan Meier analysis) of all patients according to the gastric reflux, the BMI, the sex, the tumor localization, the recurrence, the adjuvant use and the neoadjuvant use. Related to Fig. 4.

**Figure S11.** Summary tables of cox proportional hazard model coefficients made with (**A**) all clinical data available and with combined significant metadata and lipids biomarkers in (**B**) negative ion mode and (**C**) positive ion mode. Related to Fig. 4.

**Figure S12.** Mean spectra of all 52 bacterial strains in negative ion mode. Related to Fig. 5.

**Figure S13.** Classification report of bacterial strains models before oversampling, after 20- fold cross-validation in negative ion mode. Related to Fig. 5.

**Figure S14.** Mean intensity of each of the 82 lipid biomarkers in all the 52 bacterial strains. Related to Fig. 5.

**Figure S15.** Heatmap following ANOVA analysis of: (**A**) tissues by tissues with a p-value of 0.01, (**B**) tissues by tissues with a p-value of 0.05, (**C-D**) average and tissues by tissues with a p-value of 0.1, (**E-F**) average and tissues by tissues with a p-value of 1, illustrating the predicted abundance of over-expressed and under-expressed bacteria in healthy, PCC, and non-PCC tissues. Related to Fig. 6.

**Figure S16.** Summary table of Cox proportional hazard model coefficients made with both significant clinical data and specific bacteria lipids biomarkers. Related to Fig. 6.

