## Supplementary material for "Transforming Esogastric Cancer Surgery Integrating SpiderMass Mass Spectrometry with Clinical and Microbiome Data for Margin Delineation and Prognosis": supplemtary figures and tables

**Table S1.** Overview of clinical data of the EC cohort. Related to Fig. 2-3-4-5-6.

|  |  |  | **Patients (108)** |
| --- | --- | --- | --- |
| Age (in years), *mean (min-max)* | | | 63 (24-86) |
| Organ | Esophagus, *n* | | 13 |
|  | EGJ, *n* | | 39 |
|  | Stomach, *n* | | 55 |
| Gender | Woman, *n* | | 27 |
|  | Man, *n* | | 80 |
| Gastric reflux | Yes, *n* | | 21 |
|  | No, *n* | | 87 |
| Body mass index (BMI) | BMI < 18.5 | | 5 |
|  | 18.5 < BMI < 25 | | 49 |
|  | 25 < BMI < 30 | | 30 |
|  | 30 < BMI > 35 | | 7 |
|  | 35 < BMI > 40 | | 1 |
|  | BMI > 40 | | 1 |
| Neoadjuvant treatment | Yes, *n* | Chemotherapy | 50 |
|  |  | Radiochemotherapy | 10 |
|  | No, *n* | | 47 |
| Adjuvant treatment | Yes, *n* | Chemotherapy | 76 |
|  |  | Radiochemotherapy | 0 |
|  | No, *n* | | 31 |
| Death | Yes, *n* | | 70 |
|  | No, *n* | | 37 |
| Relapse | Yes, *n* | | 49 |
|  | No, *n* | | 58 |
| Overall survival (in months), *median (min-max)* | | | 40 (0-160) |

**Table S2.** Demographic and clinical data for all the EC cohort (169 tissues). Related to Fig. 2-3-4-5-6.

| **N°** | **F/H** | **pTN M** | **ASA** | **Age** | **Localisation** | **BMI** | **Gastric reflux** | **Neo-**  **adjuvant** | **Adju**  **vant** | **Rela**  **pse** | **Death** | **Survival (month)** |
| --- | --- | --- | --- | --- | --- | --- | --- | --- | --- | --- | --- | --- |
| **Other GC subtypes (Non PCCs)** | | | | | | | | | | | | |
| **1** | F | 3 | 1 | 74 | EGJ | 30.5 | No |  | Yes | Yes | Yes | 18 |
| **3** | H | 4 | 3 | 53 | Sto | 28.9 | No |  | No | No | Yes | 2 |
| **5.1** | H | 4 | 3 | 78 | Sto | 24.5 | No |  | Yes | Yes | Yes | 27 |
| **24-1** | H | 4 | 2 | 69 | EGJ | 22.2 | No |  | Yes | No | No | 126 |
| **24-2** | H | 4 | 2 | 69 | EGJ | 22.2 | No |  | Yes | No | No | 126 |
| **27** |  |  |  |  |  | 19.2 | No |  |  |  |  |  |
| **30** | F | 2 | 1 | 41 | Sto | 20.4 | No | Chemo | Yes | Yes | Yes | 22 |
| **44** | H | 3 | 2 | 80 | EGJ | 18.7 | No | Chemo | No | No | Yes | 2 |
| **45** | H | 1 | 3 | 39 | Sto | 27.7 | No |  | Yes | No | No | 35 |
| **46** | F | 3 | 3 | 70 | Sto | 28.7 | Yes | Chemo | Yes | Yes | No | 41 |
| **48** | H | 3 | 2 | 67 | Sto | 17.3 | Yes | Chemo | Yes | No | No | 35 |
| **49** | H | 3 | 2 | 41 | EGJ | 24.4 | No | Radiochem | Yes | Yes | Yes | 7 |
| **50** | H | 3 | 2 | 82 | Sto | 31.6 | No | Chemo | Yes | No | Yes | 15 |
| **52.1** | F | 3 | 2 | 78 | EGJ | 27.1 | No | Chemo | No | Yes | No | 63 |
| **52.2** | F | 3 | 2 | 78 | EGJ | 27.1 | No | Chemo | No | Yes | No | 63 |
| **55** | H | 3 | 2 | 83 | EGJ | 22.3 | No | Chemo | No | No | No | 81 |
| **56** | H | 2 | 4 | 60 | EGJ | 19.2 | No |  | Yes | No | Yes | 12 |
| **57** | F | 1 | 3 | 82 | Sto | 28.6 | No | Chemo | Yes | No | No | 49 |
| **58** | H | 2 | 3 | 55 | Sto | 30.9 | Yes |  | No | No | No | 100 |
| **59** | H | 3 | 1 | 54 | Sto | 16.3 | No | Chemo | Yes | No | No | 56 |
| **60** | H | 4 | 2 | 63 | EGJ | 32.6 | No | Chemo | Yes | Yes | Yes | 50 |
| **61** | H | 1 | 3 | 62 | EGJ | 22.3 | No |  | No | No | No | 119 |
| **62** | H | 4 | 3 | 83 | Sto | 35.6 | No |  | Yes | No | Yes | 16 |
| **63** | F | 2 | 1 | 71 | Sto | 25.5 | No | Chemo | Yes | No | No | 55 |
| **64** | H | 1 | 3 | 63 | Sto | 19.4 | No | Chemo | Yes | No | No | 40 |
| **65** | H | 3 | 1 | 57 | Sto | 28.3 | No | Chemo | Yes | Yes | Yes | 16 |
| **67** | H | 3 | 2 | 68 | Sto | 28.4 | No | Chemo | Yes | Yes | Yes | 34 |
| **69** | H | 3 | 2 | 68 | Sto | 22.5 | No | Chemo | Yes | Yes | Yes | 24 |
| **72** | H | 3 | 2 | 56 | EGJ | 25.5 | No | Chemo | Yes | Yes | Yes | 19 |
| **73** | H | 4 | 2 | 68 | Sto | 26.8 | No | Radiochem | Yes | Yes | Yes | 25 |
| **74** | H | 4 | 1 | 67 | Sto | 20.2 | Yes | Chemo | Yes | Yes | Yes | 31 |
| **75** | H | 3 | 2 | 53 | EGJ | 32.6 | No | Chemo | Yes | Yes | No | 27 |
| **78** | H | 2 | 2 | 74 | EGJ | 30.9 | No | Chemo | Yes | No | No | 46 |
| **79** | F | 4 | 3 | 57 | Sto | 26.1 | No |  | Yes | Yes | Yes | 35 |
| **81** | H | 2 | 2 | 44 | EGJ | 28.4 | No | Radiochem | No | Yes | No | 59 |
| **83** | H | 4 | 2 | 84 | EGJ | 26.8 | No | Chemo | Yes | No | Yes | 15 |
| **84** | H | 4 | 2 | 49 | EGJ | 23.7 | No |  | No | Yes | Yes | 31 |
| **85** | H | 4 | 3 | 84 | Eso | 32.2 | Yes |  | Yes | No | Yes | 32 |
| **86** | H | 3 | 2 | 75 | Sto | NK | No | Chemo | Yes | Yes | Yes | 3 |
| **88** | H | 1 | 2 | 68 | EGJ | 28.4 | No |  | No | No | No | 100 |
| **90.1** | H | 4 | 1 | 68 | EGJ | 33.9 | No |  | Yes | Yes | Yes | 119 |
| **90.2** | H | 4 | 1 | 68 | EGJ | 33.9 | No |  | Yes | Yes | Yes | 119 |
| **91** | F | 3 | 2 | 58 | EGJ | 32.6 | Yes | Radiochem | No | Yes | Yes | 23 |
| **92** | H | 3 | 2 | 60 | EGJ | NK | Yes |  | No | No | Yes | 1 |
| **93** | H | 3 | 1 | 85 | EGJ | 21.1 | Yes | Chemo | Yes | Yes | Yes | 30 |
| **94** | H | 3 | 2 | 75 | EGJ | 26.6 | Yes | Chemo | Yes | Yes | Yes | 22 |
| **95** | H | 2 | 3 | 69 | Eso | 30.1 | No |  | No | Yes | Yes | 22 |
| **96** | H | 1 | 2 | 61 | Eso | 23 | Yes |  | No | No | No | 63 |
| **97** | H | 3 | 2 | 64 | Eso | 33 | No |  | Yes | No | Yes | 21 |
| **98** | H | 3 | 2 | 73 | Eso | 29.9 | No | Chemo | Yes | No | No | 67 |
| **102** | H | 3 | 2 | 77 | EGJ | 24 | Yes | Chemo | Yes | No | No | 64 |
| **104** | F | 3 | 2 | 32 | Eso | 22 | No | Chemo | Yes | No | No | 82 |
| **105** | H | 1 | 2 | 65 | Eso | 23.9 | Yes | Chemo | Yes | No | Yes | 59 |
| **106** | H | 4 | 2 | 73 | Eso | 23.3 | Yes | Chemo | Yes | No | Yes | 4 |
| **107** | H | 1 | 3 | 73 | EGJ | 30.1 | Yes | Chemo | Yes | No | No | 88 |
| **108** | H | 4 | 2 | 66 | Eso | 22.9 | Yes | Chemo | Yes | No | Yes | 7 |
| **109** | H | 4 | 3 | 77 | Eso | 30.7 | No | Chemo | No | Yes | Yes | 23 |
| **112** | H | 4 | 3 | 66 | EGJ | 32.2 | No | Chemo | Yes | No | Yes | 11 |
| **113** | H | 4 | 1 | 63 | EGJ | 27.7 | No | Chemo | Yes | Yes | Yes | 69 |
| **116** | H | 3 | 1 | 60 | EGJ | 27.8 | No | Chemo | Yes | Yes | No | 89 |
| **PCCs** | | | | | | | | | | | | |
| **2** | F | 4 | 2 | 47 | Sto | 24.3 | No |  | Yes | No | Yes | 5 |
| **6** | H | 4 | 2 | 78 | Sto | 19.8 | No |  | No | No | Yes | 3 |
| **10** | F | 4 | 1 | 42 | Sto | 23.4 | No |  | Yes | Yes | Yes | 11 |
| **11** | F | 4 | 1 | 86 | Sto | 21.1 | No |  | No | No | Yes | 1 |
| **12** | F | 4 | 3 | 51 | Sto | 21.2 | No |  | Yes | Yes | Yes | 39 |
| **15** | H | 1 | 2 | 62 | EGJ | 22.8 | Yes |  | No | No | Yes | 13 |
| **17** | F | 4 | 0 | 41 | EGJ | 26 | No |  | Yes | No | Yes | 26 |
| **18** | H | 4 | 3 | 51 | Sto | 57 | No |  | No | No | Yes | 0 |
| **20** | F | 3 | 2 | 70 | EGJ | 32 | Yes |  | Yes | Yes | Yes | 20 |
| **22** | H | 4 | 1 | 47 | Eso | NK | No | Chemo | Yes | Yes | Yes | 7 |
| **23** | H | 4 | 2 | 83 | Sto | 22.6 | No |  | No | No | Yes | 2 |
| **28** | F | 2 | 2 | 66 | Sto | 18.6 | No |  | No | Yes | Yes | 12 |
| **29** | H | 3 | 1 | 36 | Sto | 26.7 | No |  | Yes | Yes | Yes | 19 |
| **31** | F | 2 | 2 | 50 | Sto | 22 | No | Chemo | Yes | No | No | 55 |
| **32** | H | 3 | 1 | 24 | Sto | 12.5 | No |  | Yes | Yes | No | 59 |
| **37** | H | 4 | 2 | 68 | Sto | 18.3 | No |  | Yes | No | Yes | 11 |
| **38** | H | 2 | 3 | 67 | Sto | 20.5 | No |  | Yes | Yes | Yes | 35 |
| **39** | H | 4 | 2 | 71 | Sto | 23.1 | No | Chemo | Yes | Yes | Yes | 35 |
| **40** | F | 4 | 2 | 43 | Sto | 18.4 | No | Chemo | Yes | Yes | Yes | 40 |
| **42** | H | 4 | 3 | 67 | EGJ | 29.3 | No | Radiochem | No | No | Yes | 1 |
| **47** | H | 4 | 2 | 57 | EGJ | 28 | No | Chemo | No | Yes | No | 55 |
| **51** | H | 3 | 3 | 71 | Sto | 22 | No | Radiochem | No | No | Yes | 4 |
| **76** | H | 2 | 3 | 72 | Eso | 21.5 | Yes | Radiochem | No | No | Yes | 1 |
| **77** | H | 3 | 2 | 60 | Sto | 23.1 | Yes | Radiochem | Yes | No | No | 40 |
| **80** | F | 4 | 2 | 46 | Sto | 18.8 | No |  | Yes | Yes | Yes | 41 |
| **87** | F | 4 | 1 | 60 | Sto | 14 | No | Chemo | Yes | No | No | 55 |
| **114** | H | 3 | 2 | 51 | EGJ | NK | No | Chemo | Yes | Yes | Yes | 12 |
| **Healthy** | | | | | | | | | | | | |
| **1** | F | 3 | 1 | 74 | EGJ | 30.5 | No |  | Yes | Yes | Yes | 18 |
| **2** | F | 4 | 2 | 47 | Sto | 24.3 | No |  | Yes | No | Yes | 5 |
| **3** | H | 4 | 3 | 53 | Sto | 28.9 | No |  | No | No | Yes | 2 |
| **4** | H | 1 | 1 | 49 | Sto | 33 | No | Chemo | Yes | No | No | 63 |
| **7** | F | 1 | 1 | 54 | Sto | 32.4 | No |  | No | No | No | 159 |
| **8-1** | F | 3 | 1 | 36 | EGJ | 21.7 | No |  | Yes | Yes | Yes | 25 |
| **9** | H | 3 | 2 | 65 | Sto | 25.1 | No |  | Yes | Yes | Yes | 8 |
| **10** | F | 4 | 1 | 42 | Sto | 23.4 | No |  | Yes | Yes | Yes | 11 |
| **12** | F | 4 | 3 | 51 | Sto | 21.2 | No |  | Yes | Yes | Yes | 39 |
| **13** | H | 3 | 2 | 68 | Sto | 24.9 | No |  | Yes | No | No | 160 |
| **14** | H | 3 | 2 | 48 | Sto | 23.6 | No | Chemo | Yes | No | No | 87 |
| **17** | F | 4 | 0 | 41 | EGJ | 26 | No |  | Yes | No | Yes | 26 |
| **18** | H | 4 | 3 | 51 | Sto | 57 | No |  | No | No | Yes | 0 |
| **19** | H | 3 | 2 | 76 | Sto | 28.4 | No | Chemo | Yes | Yes | Yes | 6 |
| **20** | F | 3 | 2 | 70 | EGJ | 32 | Yes |  | Yes | Yes | Yes | 20 |
| **21** | F | 3 | 2 | 56 | Sto | 28.8 | No |  | No | No | Yes | 2 |
| **22** | H | 4 | 1 | 47 | Eso | NK | No | Chemo | Yes | Yes | Yes | 7 |
| **23** | H | 4 | 2 | 83 | Sto | 22.6 | No |  | No | No | Yes | 2 |
| **25** | F | 3 | 1 | 50 | Sto | 20.4 | No |  | Yes | Yes | Yes | 16 |
| **26.1** | H | 3 | 1 | 34 | EGJ | 23.7 | No | Radiochem | No | No | No | 130 |
| **26.2** | H | 3 | 1 | 34 | EGJ | 23.7 | No | Radiochem | No | No | No | 130 |
| **27** |  |  |  |  |  | 19.2 | No |  |  |  |  |  |
| **28** | F | 2 | 2 | 66 | Sto | 18.6 | No |  | No | Yes | Yes | 12 |
| **29** | H | 3 | 1 | 36 | Sto | 26.7 | No |  | Yes | Yes | Yes | 19 |
| **31** | F | 2 | 2 | 50 | Sto | 22 | No | Chemo | Yes | No | No | 55 |
| **32** | H | 3 | 1 | 24 | Sto | 12.5 | No |  | Yes | Yes | No | 59 |
| **33.1** | H | 1 | 2 | 79 | Sto | 22.5 | No |  | No | No | Yes | 49 |
| **33.2** | H | 1 | 2 | 79 | Sto | 22.5 | No |  | No | No | Yes | 49 |
| **34** | H | 4 | 2 | 61 | Sto | 23 | No | Chemo | Yes | Yes | Yes | 9 |
| **35** | H | 3 | 2 | 59 | EGJ | 24.7 | No |  | Yes | Yes | Yes | 67 |
| **36.1** | F | 4 | 2 | 51 | Sto | 21 | No | Chemo | Yes | Yes | Yes | 13 |
| **36.2** | F | 4 | 2 | 51 | Sto | 21 | No | Chemo | Yes | Yes | Yes | 13 |
| **38** | H | 2 | 3 | 67 | Sto | 20.5 | No |  | Yes | Yes | Yes | 35 |
| **39** | H | 4 | 2 | 71 | Sto | 23.1 | No | Chemo | Yes | Yes | Yes | 35 |
| **40** | F | 4 | 2 | 43 | Sto | 18.4 | No | Chemo | Yes | Yes | Yes | 40 |
| **41.1** | H | 1 | 4 | 68 | Sto | 18.9 | No | Chemo | Yes | No | No | 55 |
| **41.2** | H | 1 | 4 | 68 | Sto | 18.9 | No | Chemo | Yes | No | No | 55 |
| **43** | H | 2 | 3 | 71 | Sto | 26.2 | No | Chemo | Yes | No | No | 49 |
| **44** | H | 3 | 2 | 80 | EGJ | 18.7 | No | Chemo | No | No | Yes | 2 |
| **45** | H | 1 | 3 | 39 | Sto | 27.7 | No |  | Yes | No | No | 35 |
| **46** | F | 3 | 3 | 70 | Sto | 28.7 | Yes | Chemo | Yes | Yes | No | 41 |
| **47** | H | 4 | 2 | 57 | EGJ | 28 | No | Chemo | No | Yes | No | 55 |
| **48** | H | 3 | 2 | 67 | Sto | 17.3 | Yes | Chemo | Yes | No | No | 35 |
| **49** | H | 3 | 2 | 41 | EGJ | 24.4 | No | Radiochem | Yes | Yes | Yes | 7 |
| **50** | H | 3 | 2 | 82 | Sto | 31.6 | No | Chemo | Yes | No | Yes | 15 |
| **51** | H | 3 | 3 | 71 | Sto | 22 | No | Radiochem | No | No | Yes | 4 |
| **53** | H | 4 | 1 | 56 | EGJ | 24.1 | No | Chemo | No | Yes | Yes | 16 |
| **54.1** | H | 3 | 3 | 76 | Sto | 24.2 | No |  | Yes | Yes | Yes | 18 |
| **54.2** | H | 3 | 3 | 76 | Sto | 24.2 | No |  | Yes | Yes | Yes | 18 |
| **55** | H | 3 | 2 | 83 | EGJ | 22.3 | No | Chemo | No | No | No | 81 |
| **56** | H | 2 | 4 | 60 | EGJ | 19.2 | No |  | Yes | No | Yes | 12 |
| **57** | F | 1 | 3 | 82 | Sto | 28.6 | No | Chemo | Yes | No | No | 49 |
| **58** | H | 2 | 3 | 55 | Sto | 30.9 | Yes |  | No | No | No | 100 |
| **59** | H | 3 | 1 | 54 | Sto | 16.3 | No | Chemo | Yes | No | No | 56 |
| **60** | H | 4 | 2 | 63 | EGJ | 32.6 | No | Chemo | Yes | Yes | Yes | 50 |
| **61** | H | 1 | 3 | 62 | EGJ | 22.3 | No |  | No | No | No | 119 |
| **63** | F | 2 | 1 | 71 | Sto | 25.5 | No | Chemo | Yes | No | No | 55 |
| **64** | H | 1 | 3 | 63 | Sto | 19.4 | No | Chemo | Yes | No | No | 40 |
| **65** | H | 3 | 1 | 57 | Sto | 28.3 | No | Chemo | Yes | Yes | Yes | 16 |
| **66-1** | F | 3 | 2 | 50 | Sto | 23.2 | No | Chemo | Yes | No | No | 130 |
| **67** | H | 3 | 2 | 68 | Sto | 28.4 | No | Chemo | Yes | Yes | Yes | 34 |
| **68-1** | H | 2 | 1 | 60 | Sto | 27 | No | Chemo | Yes | No | No | 130 |
| **69** | H | 3 | 2 | 68 | Sto | 22.5 | No | Chemo | Yes | Yes | Yes | 24 |
| **71** | H | 1 | 2 | 81 | Sto | 25.3 | No |  | No | No | Yes | 53 |
| **72** | H | 3 | 2 | 56 | EGJ | 25.5 | No | Chemo | Yes | Yes | Yes | 19 |
| **74** | H | 4 | 1 | 67 | Sto | 20.2 | Yes | Chemo | Yes | Yes | Yes | 31 |
| **78** | H | 2 | 2 | 74 | EGJ | 30.9 | No | Chemo | Yes | No | No | 46 |
| **81** | H | 2 | 2 | 44 | EGJ | 28.4 | No | Radiochem | No | Yes | No | 59 |
| **83** | H | 4 | 2 | 84 | EGJ | 26.8 | No | Chemo | Yes | No | Yes | 15 |
| **84** | H | 4 | 2 | 49 | EGJ | 23.7 | No |  | No | Yes | Yes | 31 |
| **86** | H | 3 | 2 | 75 | Sto | NK | No | Chemo | Yes | Yes | Yes | 3 |
| **89** | H | 3 | 2 | 68 | EGJ | 29.1 | Yes | Radiochem | No | No | Yes | 46 |
| **96** | H | 1 | 2 | 61 | Eso | 23 | Yes |  | No | No | No | 63 |
| **98** | H | 3 | 2 | 73 | Eso | 29.9 | No | Chemo | Yes | No | No | 67 |
| **99** | H | 3 | 2 | 66 | EGJ | 24.2 | No | Chemo | Yes | Yes | Yes | 11 |
| **100** | H | 1 | 1 | 53 | Eso | 30.4 | Yes | Chemo | Yes | No | No | 65 |
| **101** | F | 3 | 2 | 74 | EGJ | 28.1 | No |  | Yes | No | No | 37 |
| **Mixte** | | | | | | | | | | | | |
| **8-2** | F | 3 | 1 | 36 | EGJ | 21.7 | No |  | Yes | Yes | Yes | 25 |
| **66-2** | F | 3 | 2 | 50 | Sto | 23.2 | No | Chemo | Yes | No | No | 130 |
| **68-2** | H | 2 | 1 | 60 | Sto | 27 | No | Chemo | Yes | No | No | 130 |
| **82** | H | 4 | 1 | 49 | Sto | 20.3 | No | Chemo | Yes | Yes | Yes | 22 |
| **103** | H | 4 | 2 | 82 | EGJ | 26.7 | Yes | Chemo | Yes | No | Yes | 13 |

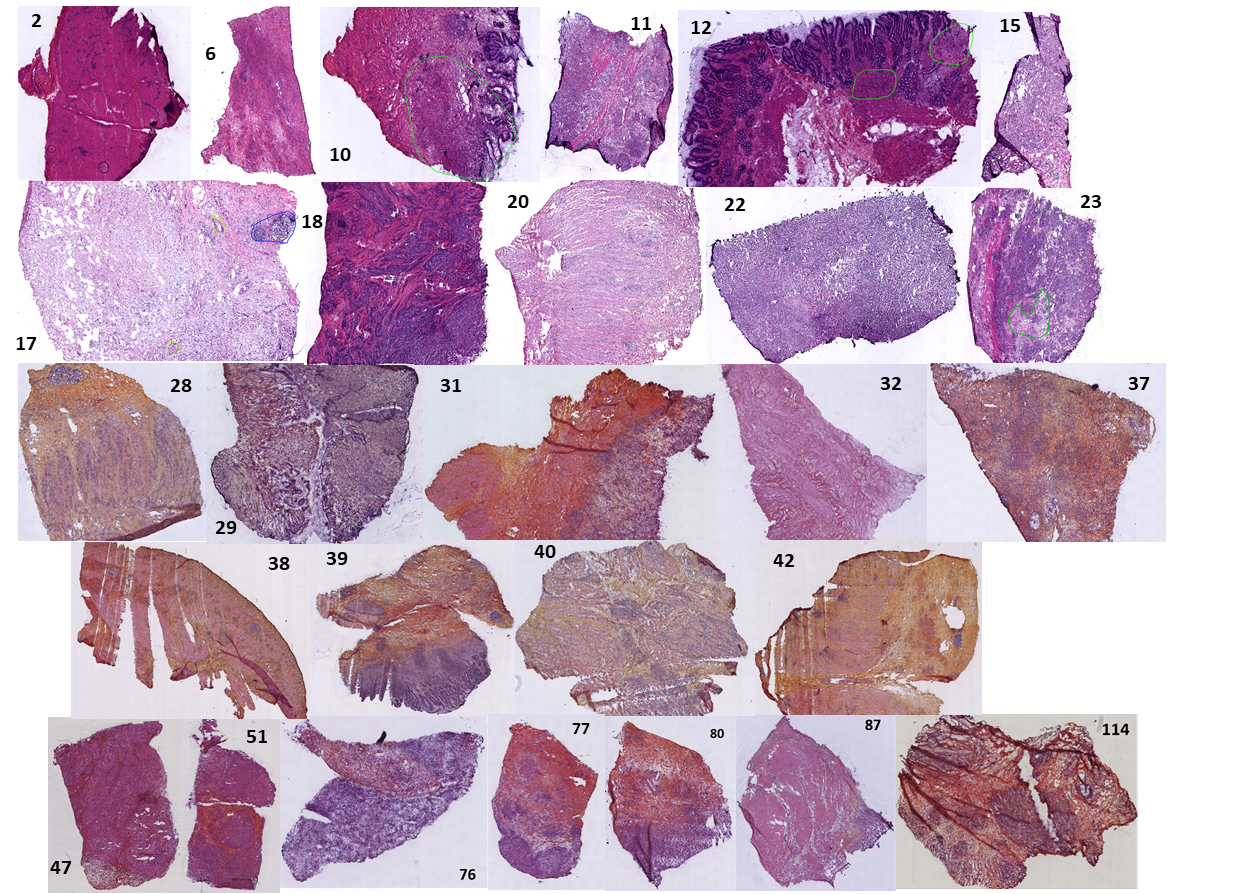

**Figure S1.** H&E scans of the 27 PCC esogastric tissues from the total EC prospective cohort. Related to Fig. 2.

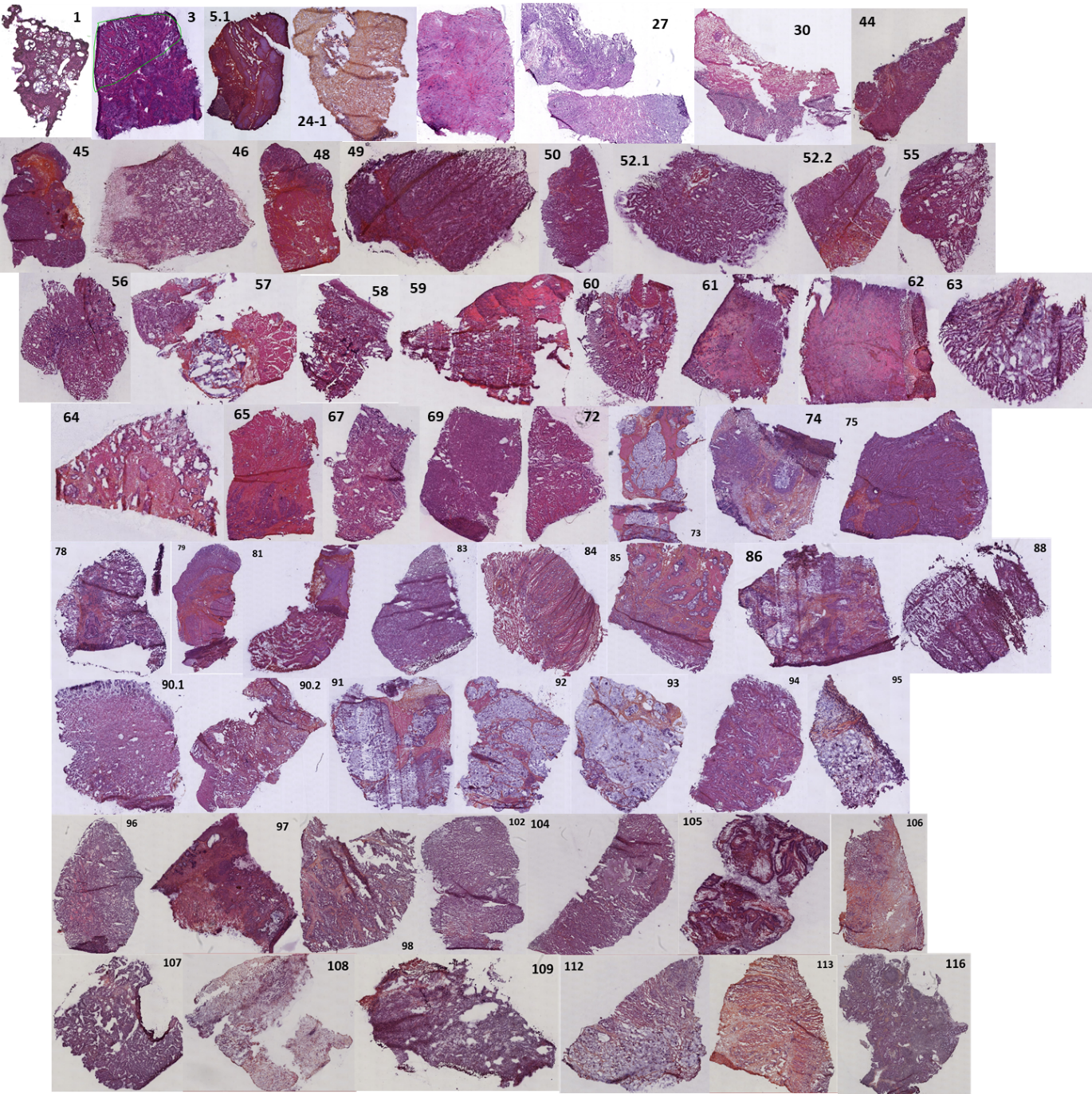

**Figure S2.** H&E scans of the 60 no-PCC esogastric tissues from the total EC prospective cohort. Related to Fig. 2.

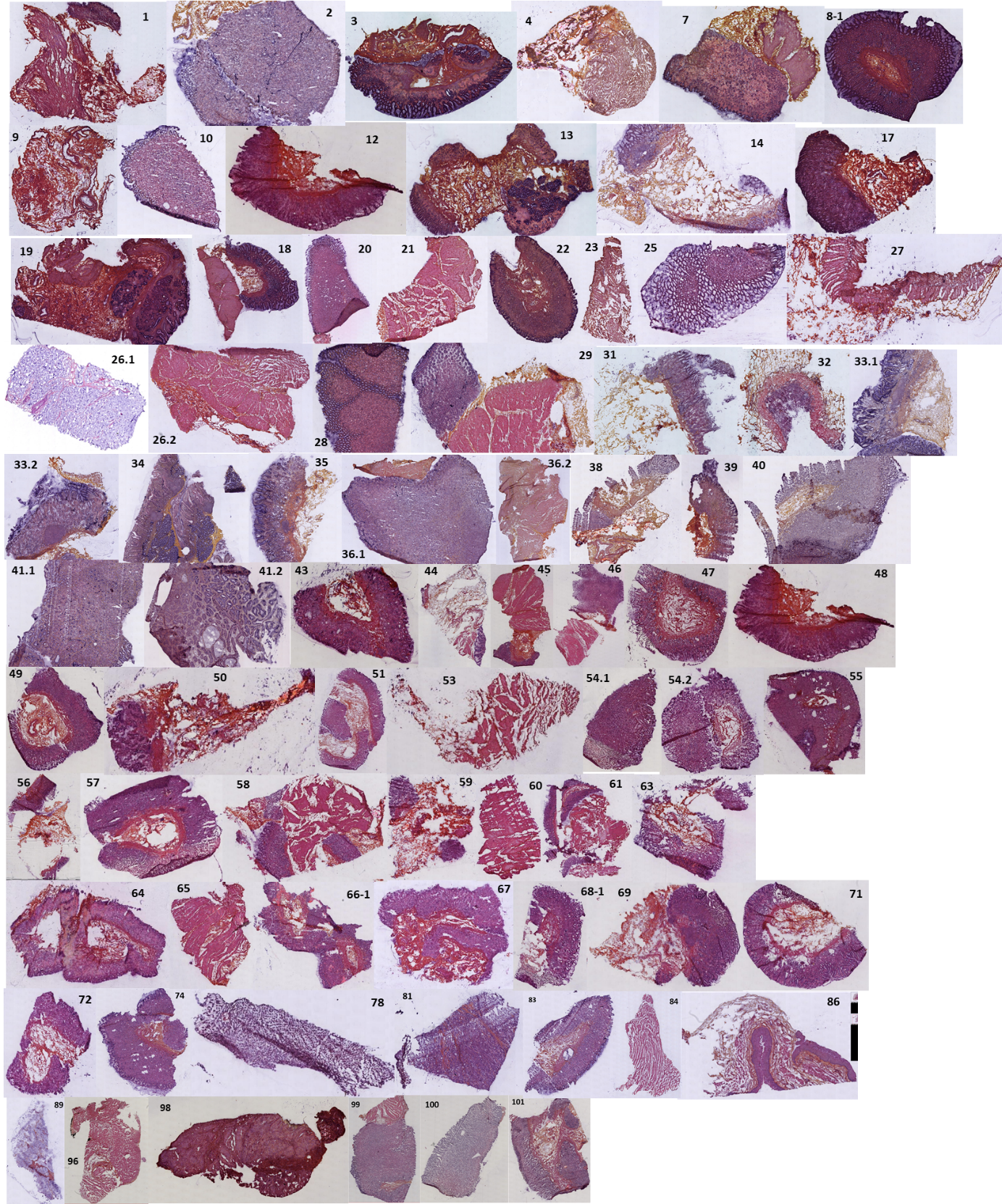

**Figure S3.** H&E scans of the 77 healthy esogastric tissues from the total EC prospective cohort. Related to Fig. 2.

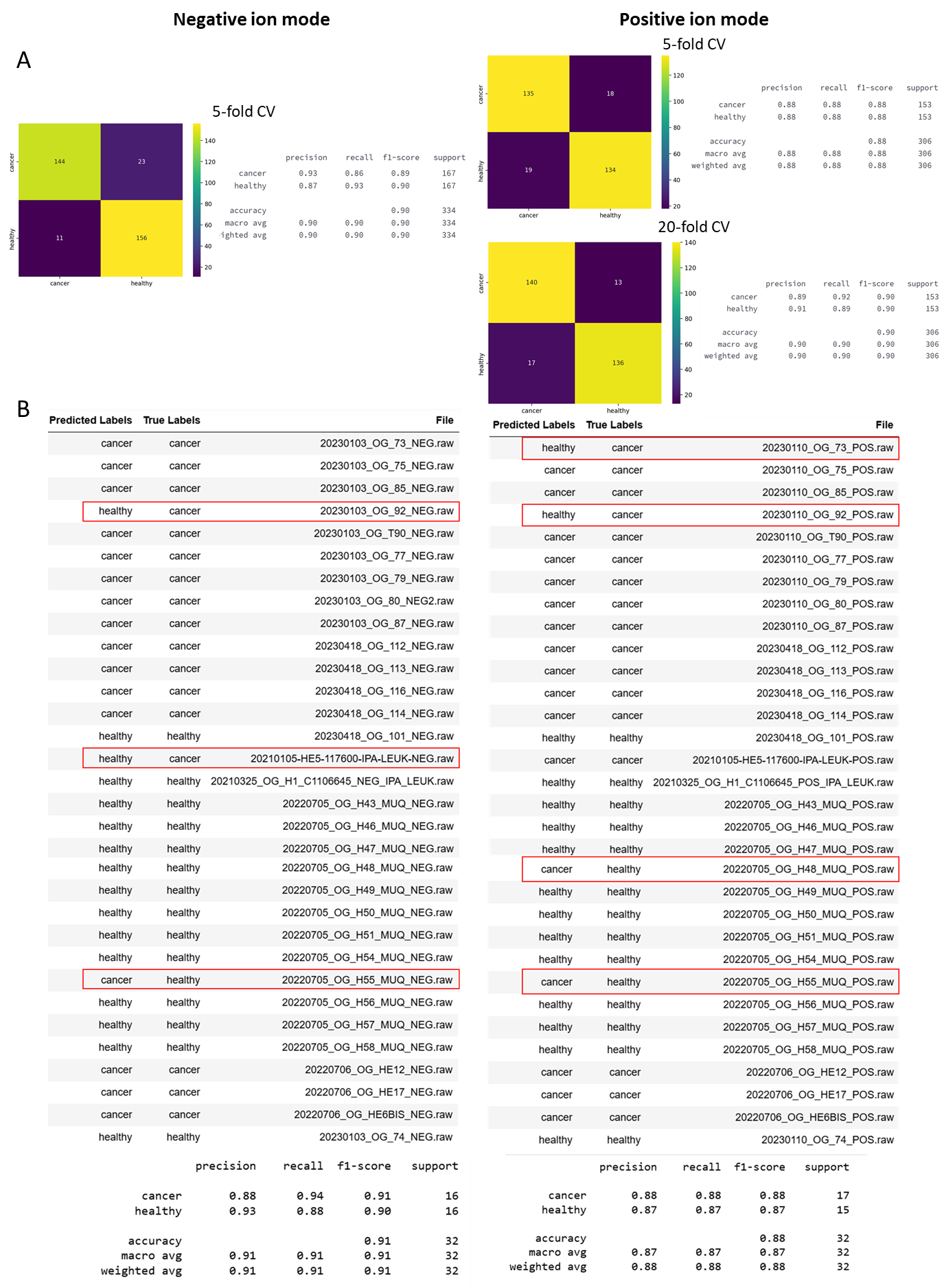

**Figure S4.** (A) Classification report and confusion matrix after 5- and 20-fold cross-validation of the classification model built from the negative mode and positive mode. SpiderMass using the RidgeClassifier algorithm to discriminate cancer from healthy tissues from 132 tissues of the cohort. (B) Results from the blind interrogation of this model based on the 32 remaining tissues of the cohort (summary table and classification report). Related to Fig. 2.

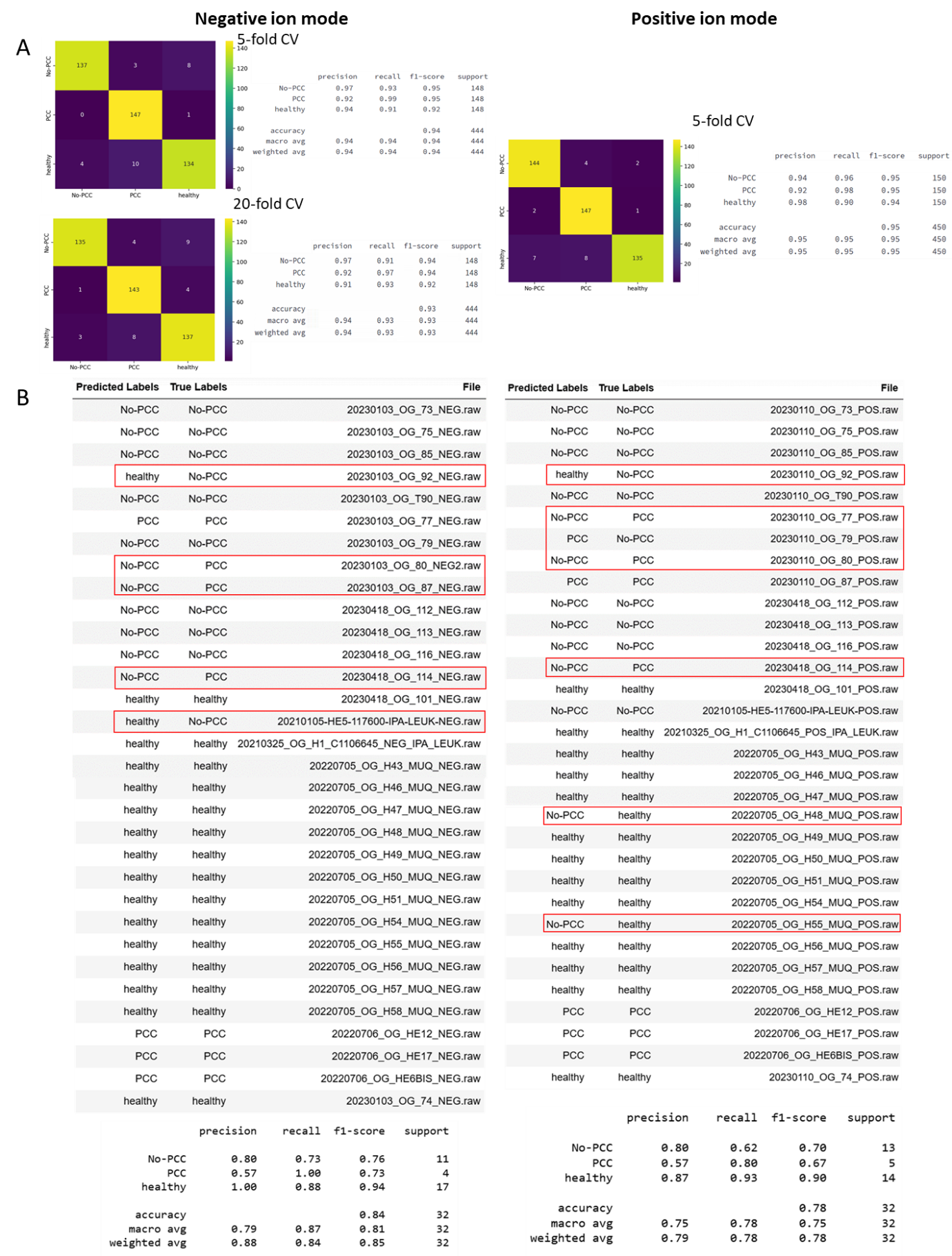

**Figure S5.** (A) Classification report and confusion matrix after 5- and 20-fold cross-validation of the classification model built from the negative mode and positive mode. SpiderMass using the RidgeClassifier algorithm to discriminate PCC vs non-PCC vs healthy from 132 tissues of the cohort. (B) Results from the blind interrogation of this model based on the 32 remaining tissues of the cohort (summary table and classification report). Related to Fig. 2.

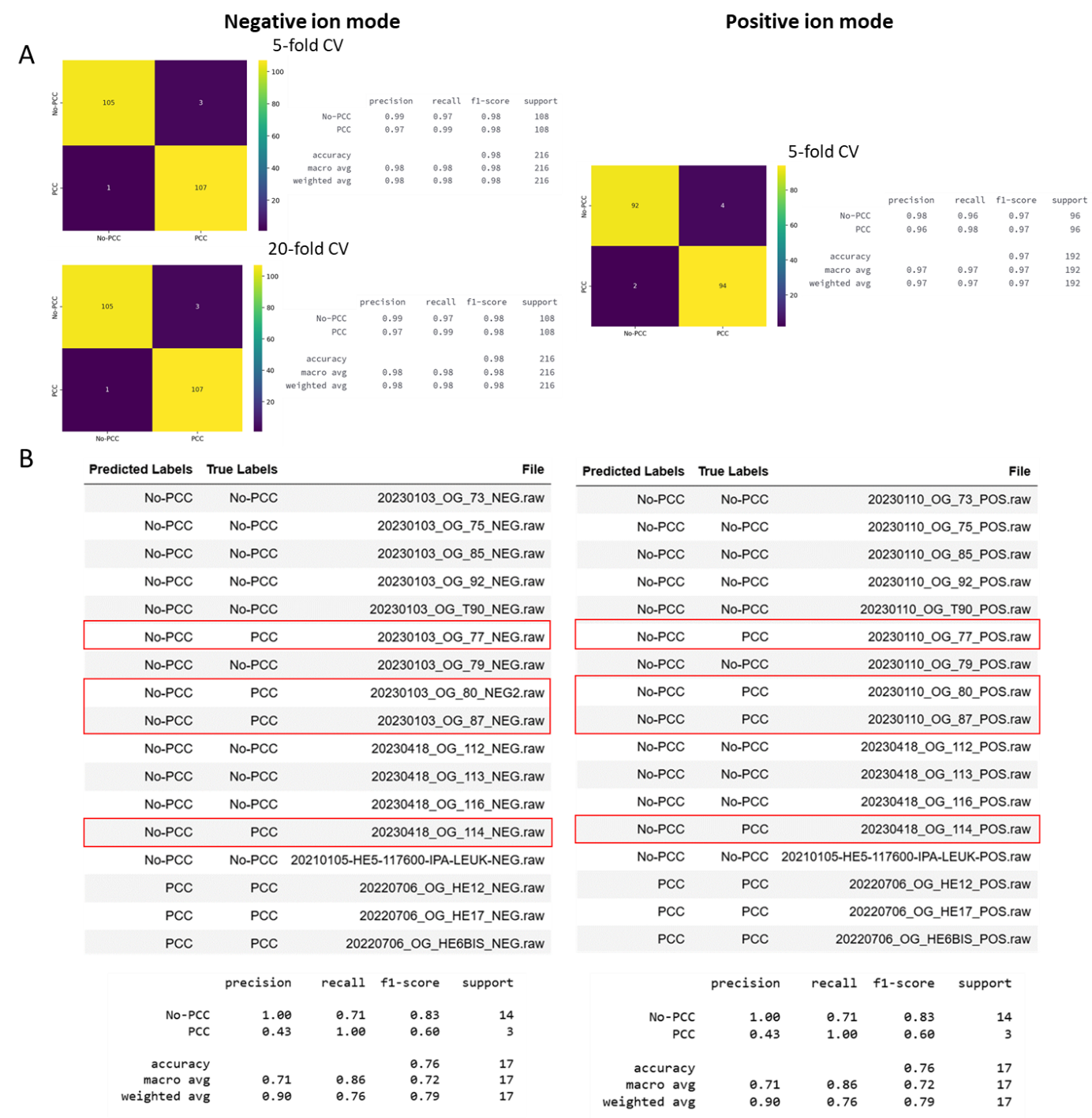

**Figure S6.** (A) Classification report and confusion matrix after 5- and 20-fold cross-validation of the classification model built from the negative mode and positive mode. SpiderMass using the RidgeClassifier algorithm to discriminate PCC vs the other subtypes of GC (non-PCCs) from 132 tissues of the cohort. (B) Results from the blind interrogation of this model based on the 32 remaining tissues of the cohort (summary table and classification report). Related to Fig. 2.

**Table S3.** Comprehensive list of reliable biomarkers for healthy and cancerous tissues. as well as for PCC and non-PCC cancer subtypes in both ion modes. Related to Fig. 2.

| **Negative ion mode** | | | | | |
| --- | --- | --- | --- | --- | --- |
| *m/z* | Cancer vs Healthy | | PCC vs non-PCC | | Putative annotation based on MS2 analysis |
|  | **Cancer** | **Healthy** | **PCC** | **Non-PCC** |  |
| 639.25 |  |  |  |  | [PA (26:7)-H]^-^ |
| 671.45 |  |  |  |  | [PA (34:2)-H]^-^ |
| 673.45 |  |  |  |  | [PA (34:1)-H]^-^ |
| 678.45 |  |  |  |  | [PE (32:6)-H]^-^ |
| 684.65 |  |  |  |  | [Cer d42:1+Cl]^-^ |
| 687.55 |  |  |  |  | [PA (O-36:1)-H]^-^ |
| 697.45 |  |  |  |  | [PA (18:1_18:2)-H]^-^ |
| 698.55 |  |  |  |  | [Cer (d18:2_24:0(2OH))+Cl]^-^ |
| 699.55 |  |  |  |  | [PA (36:2)-H]^-^ |
| 700.55 |  |  |  |  | [Cer (d18:1_24:0(2OH))+Cl]^-^ |
| 701.55 |  |  |  |  | [PA (36:1)-H]^-^ |
| 714.55 |  |  |  |  | [PE (16:0_18:2)-H]^-^ |
| 718.55 |  |  |  |  | [PE (16:0_18:0)-H]^-^ |
| 720.45 |  |  |  |  | [PE (32:2)+Cl]^-^ |
| 722.55 |  |  |  |  | [PE (P-16:0_20:4)-H]^-^ |
| 726.55 |  |  |  |  | [PE (P-18:0_18:2)-H]^-^ |
| 728.55 |  |  |  |  | [PE (P-18:1_18:0)-H]^-^ |
| 733.45 |  |  |  |  | [PA (36:3)+Cl]^-^ |
| 734.55 |  |  |  |  | [PE (16:2_20:4)-H]^-^ |
| 736.55 |  |  |  |  | [PE (36:5)-H]^-^ |
| 738.55 |  |  |  |  | [PE (16:0_20:4)-H]^-^ |
| 740.55 |  |  |  |  | [PE (36:3)-H]^-^ |
| 744.55 |  |  |  |  | [PE (18:0_18:1)-H]^-^ |
| 748.55 |  |  |  |  | [PS (O-34:0)-H]^-^ |
| 750.55 |  |  |  |  | [PE (P-18:0_20:4)-H]^-^ |
| 762.55 |  |  |  |  | [PE (38:6)-H]^-^ |
| 764.55 |  |  |  |  | [PE (18:1_20:4)-H]^-^ |
| 772.55 |  |  |  |  | [PE (18:1_20:0)-H]^-^ |
| 778.55 |  |  |  |  | [PS (36:6)-H]^-^ |
| 788.55 |  |  |  |  | [PS (18:0_18:1)-H]^-^ |
| 790.55 |  |  |  |  | [PS (18:0_18:0)-H]^-^ |
| 794.55 |  |  |  |  | [PE (40:4)-H]^-^ |
| 797.65 |  |  |  |  | [TG (48:4)-H]^-^ |
| 819.55 |  |  |  |  | [PG (18:1_22:6)-H]^-^ |
| 820.55 |  |  |  |  | [PE (42:5)-H]^-^ |
| 835.55 |  |  |  |  | [PI (18:1_16:0)-H]^-^ |
| 846.65 |  |  |  |  | [HexCer (d18:1/24:0)-H]^-^ |
| 864.65 |  |  |  |  | [PS (42:5)-H]^-^ |
| 883.55 |  |  |  |  | [PI (18:1_20:4)-H]^-^ |
| 981.25 |  |  |  |  | / |
| 1006.65 |  |  |  |  | [Hex2Cer (38:2)-H]^-^ |
| **Positive ion mode** | | | | | |
| *m/z* | Cancer vs Healthy | | PCC vs Non-PCC | | Putative annotation based on MS2 analysis |
|  | **Cancer** | **Healthy** | **PCC** | **Non-PCC** |  |
| 601.55 |  |  |  |  | / |
| 605.55 |  |  |  |  | [DG (36:1)+H]^+^ |
| 627.55 |  |  |  |  | [PA (32:11)+H]^+^ |
| 630.65 |  |  |  |  | / |
| 650.65 |  |  |  |  | [Cer (42:1)+H]^+^ |
| 663.45 |  |  |  |  | [PG (28:2)+H]^+^ |
| 666.45 |  |  |  |  | [PS (O-38:0)+H]^+^ |
| 685.45 |  |  |  |  | [PA (O-36:3)+H]^+^ |
| 689.55 |  |  |  |  | [PA (36:8)+H]^+^ |
| 702.55 |  |  |  |  | [PE (O-34:2)+H]^+^ |
| 703.55 |  |  |  |  | [SM (d16:1_18:0)+H]^+^ |
| 710.65 |  |  |  |  | [Cer (44:1)+H]^+^ |
| 712.65 |  |  |  |  | [Cer (t20:0_24:0 (2OH))+H]^+^ |
| 716.55 |  |  |  |  | [PE (18:1_16:1)+H]^+^ |
| 720.55 |  |  |  |  | [PS (P-32:0)+H]^+^ |
| 723.45 |  |  |  |  | [PA (38:5)+H]^+^ |
| 724.55 |  |  |  |  | [PC (32:5)+H]^+^ |
| 728.55 |  |  |  |  | [PE (O-36:3)+H]^+^ |
| 730.55 |  |  |  |  | [PE (O-36:2)+H]^+^ |
| 734.55 |  |  |  |  | [PE (36:7)+H]^+^ |
| 742.55 |  |  |  |  | [PE (36:3)+H]^+^ |
| 744.55 |  |  |  |  | [PE (36:2)+H]^+^ |
| 750.55 |  |  |  |  | [PE (O-38:6)+H]^+^ |
| 758.55 |  |  |  |  | [PE (34:2)+H]^+^ |
| 766.55 |  |  |  |  | [PC (O-36:5)+H]^+^ |
| 774.55 |  |  |  |  | [PS (O-36:2)+H]^+^ |
| 780.55 |  |  |  |  | [PC (22:4_14:1)+H]^+^ |
| 784.55 |  |  |  |  | [PC (36:3)+H]^+^ |
| 788.65 |  |  |  |  | [PC (36:1)+H]^+^ |
| 792.55 |  |  |  |  | [PE (40:6)+H]^+^ |
| 794.55 |  |  |  |  | [PE (40:5)+H]^+^ |
| 805.55 |  |  |  |  | [PA (44:6)+H]^+^ |
| 808.55 |  |  |  |  | [PC (38:5)+H]^+^ |
| 810.65 |  |  |  |  | [PE (O-42:4)+H]^+^ |
| 812.55 |  |  |  |  | [PC (38:3)+H]^+^ |
| 814.55 |  |  |  |  | [PE (42:9)+H]^+^ |
| 855.55 |  |  |  |  | [PI (36:6)+H]^+^ |
| 875.55 |  |  |  |  | [TG (54:8)+H]^+^ |
| 907.35 |  |  |  |  | [TG (56:6)+H]^+^ |

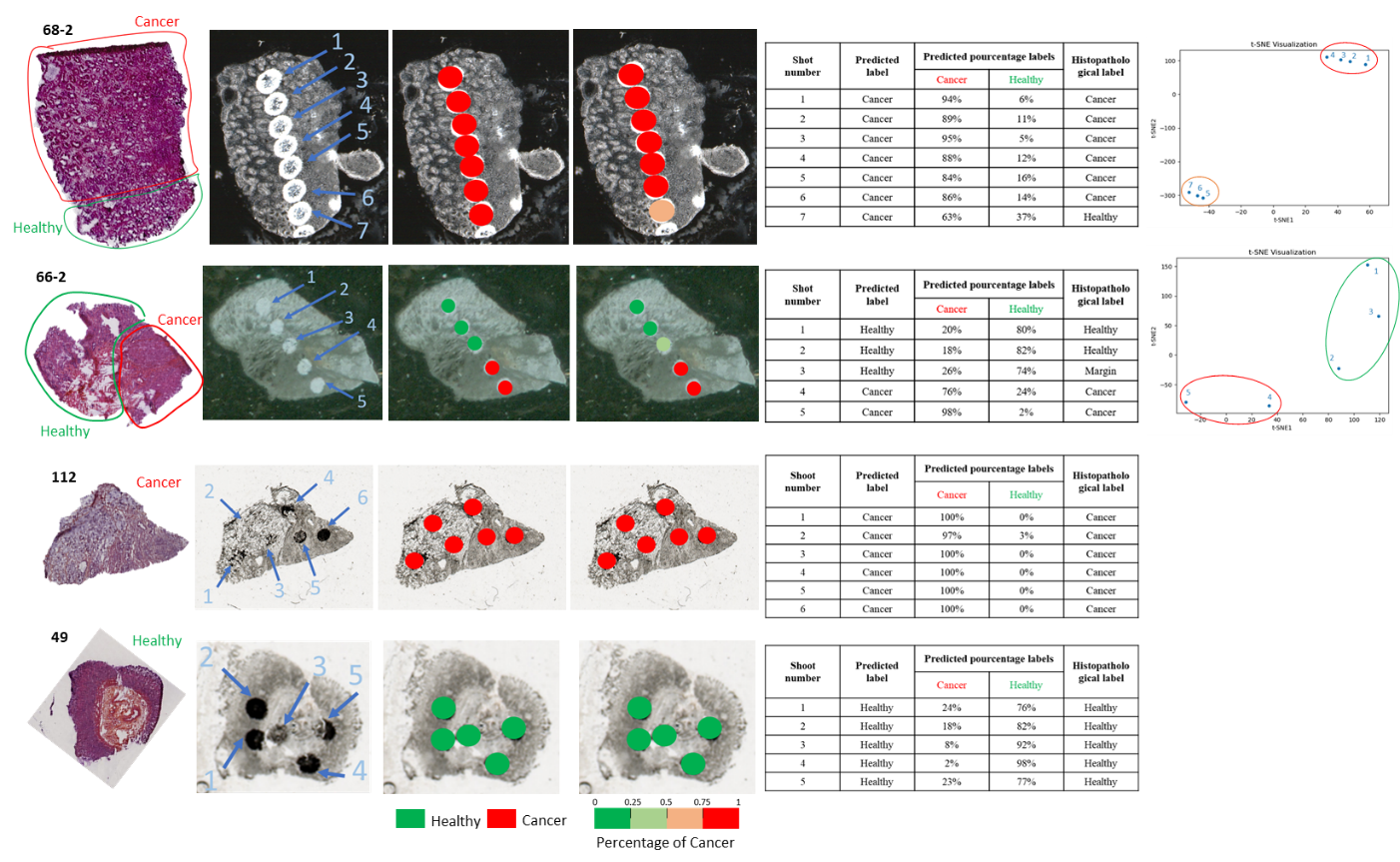

**Figure S7.** Margin delineation using SpiderMass technology of 4 additional mixed, healthy and cancerous tissues. Related to Fig. 3.

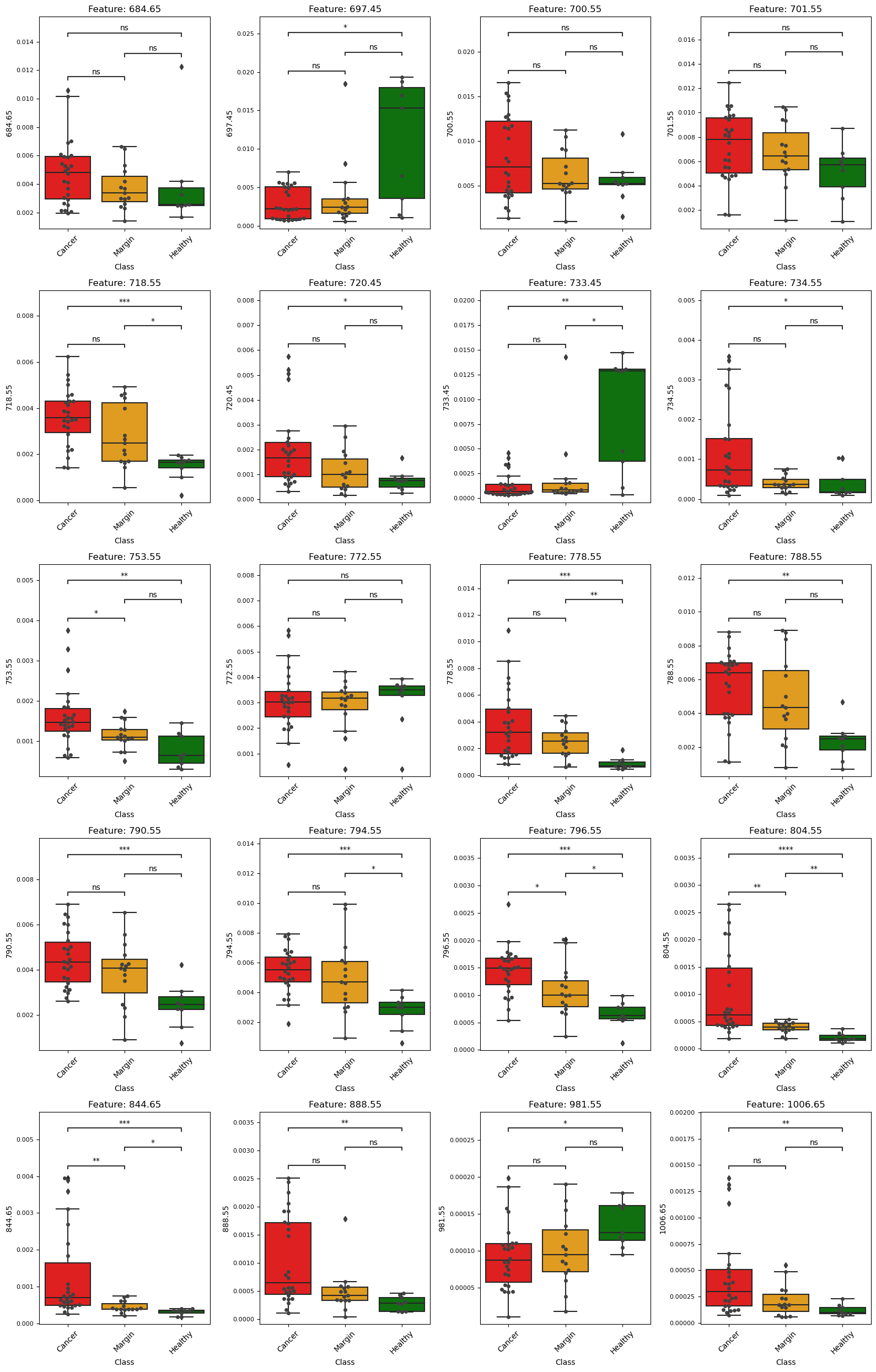

**Figure S8.** Boxplots of 20 ions overexpressed in healthy or cancerous tissue but here between healthy, margin and cancerous areas. Their putative annotations can be found in the Table S3. Related to Fig. 3.

Figure S9. Boxplots of 4 ions overexpressed in cancerous tissue but here between healthy, peri-healthy margin, peri-cancer margin and cancerous areas. Their putative annotations can be found in the Table S3. Related to Fig. 3.

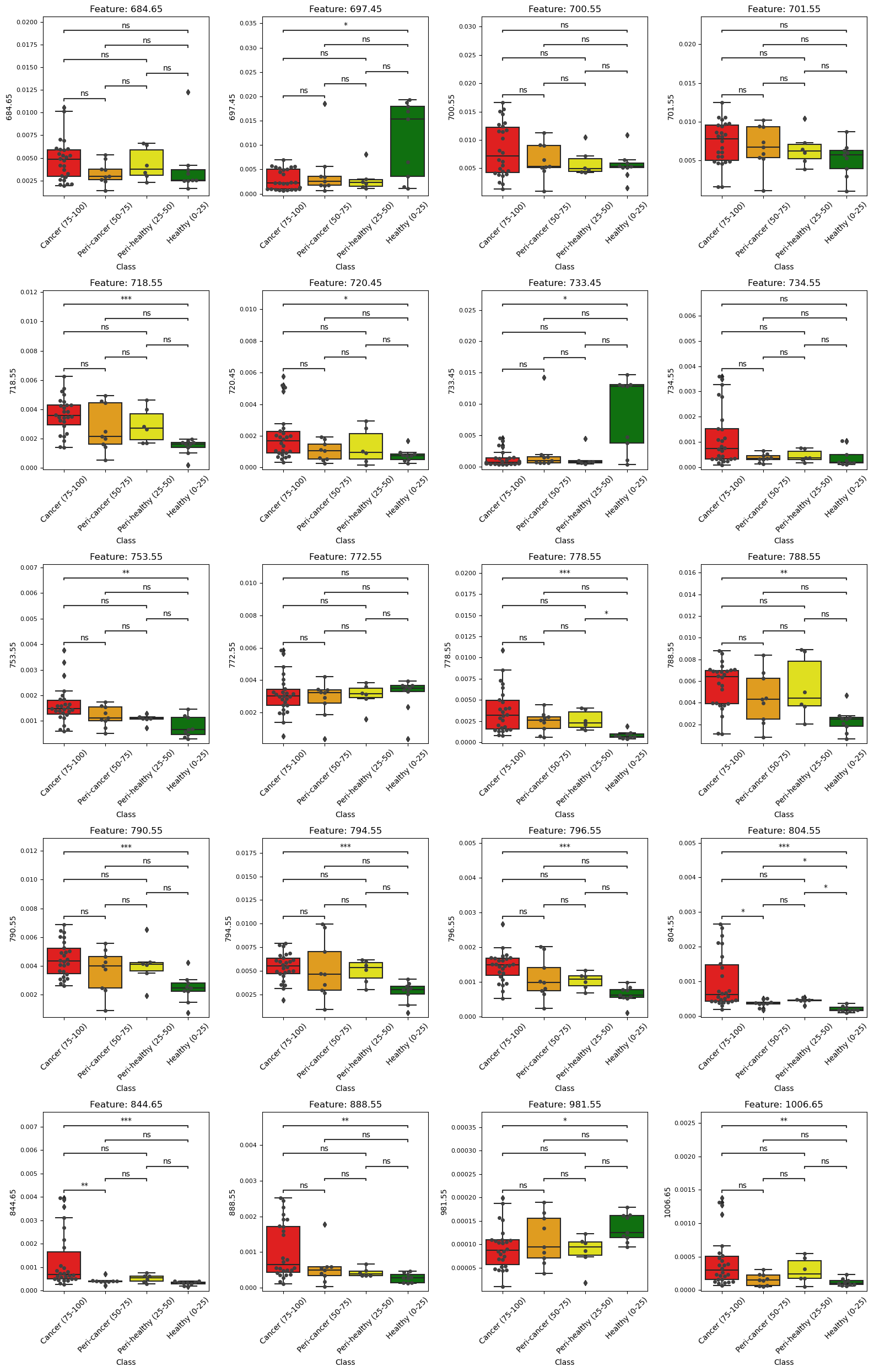

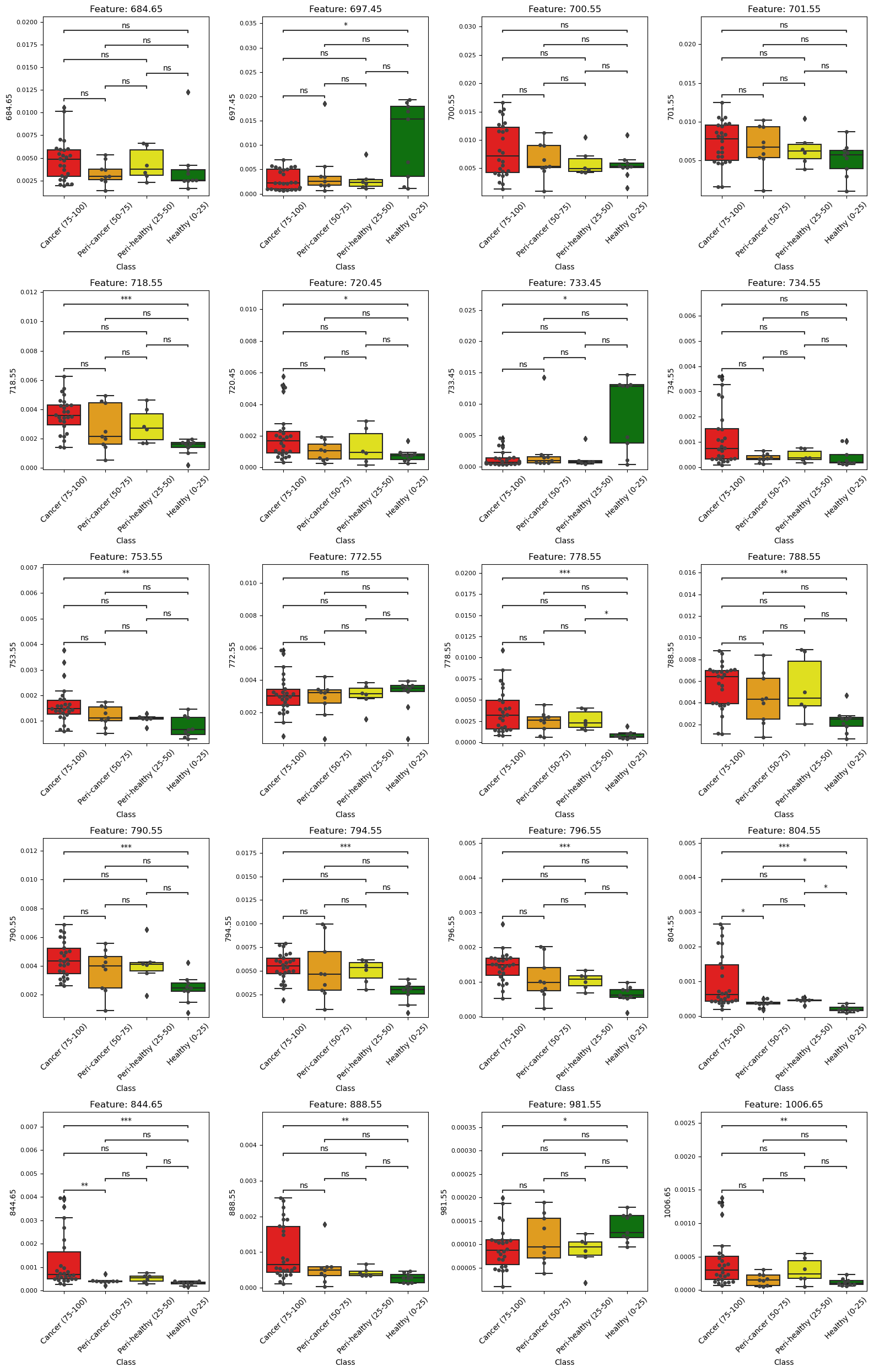

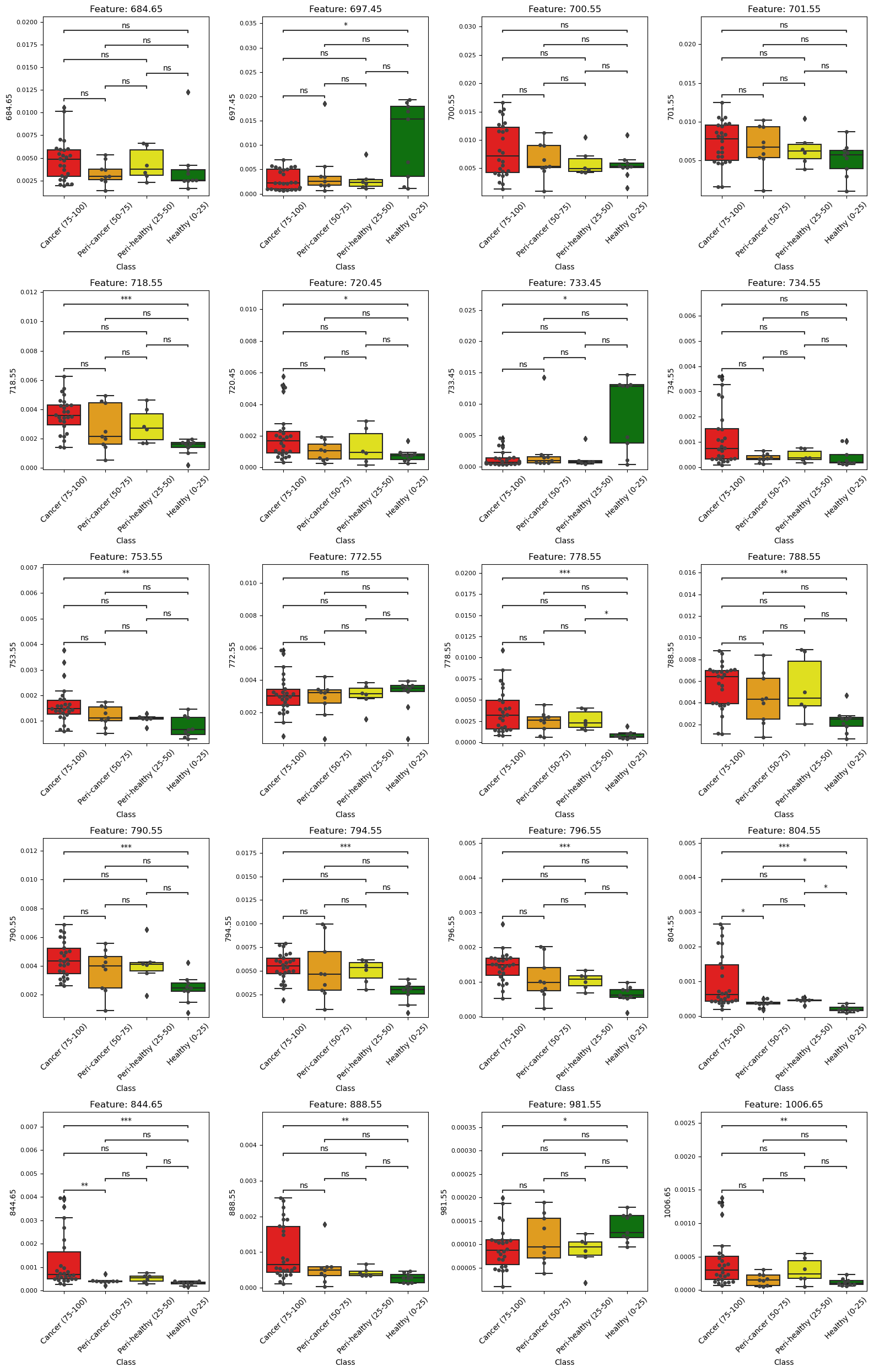

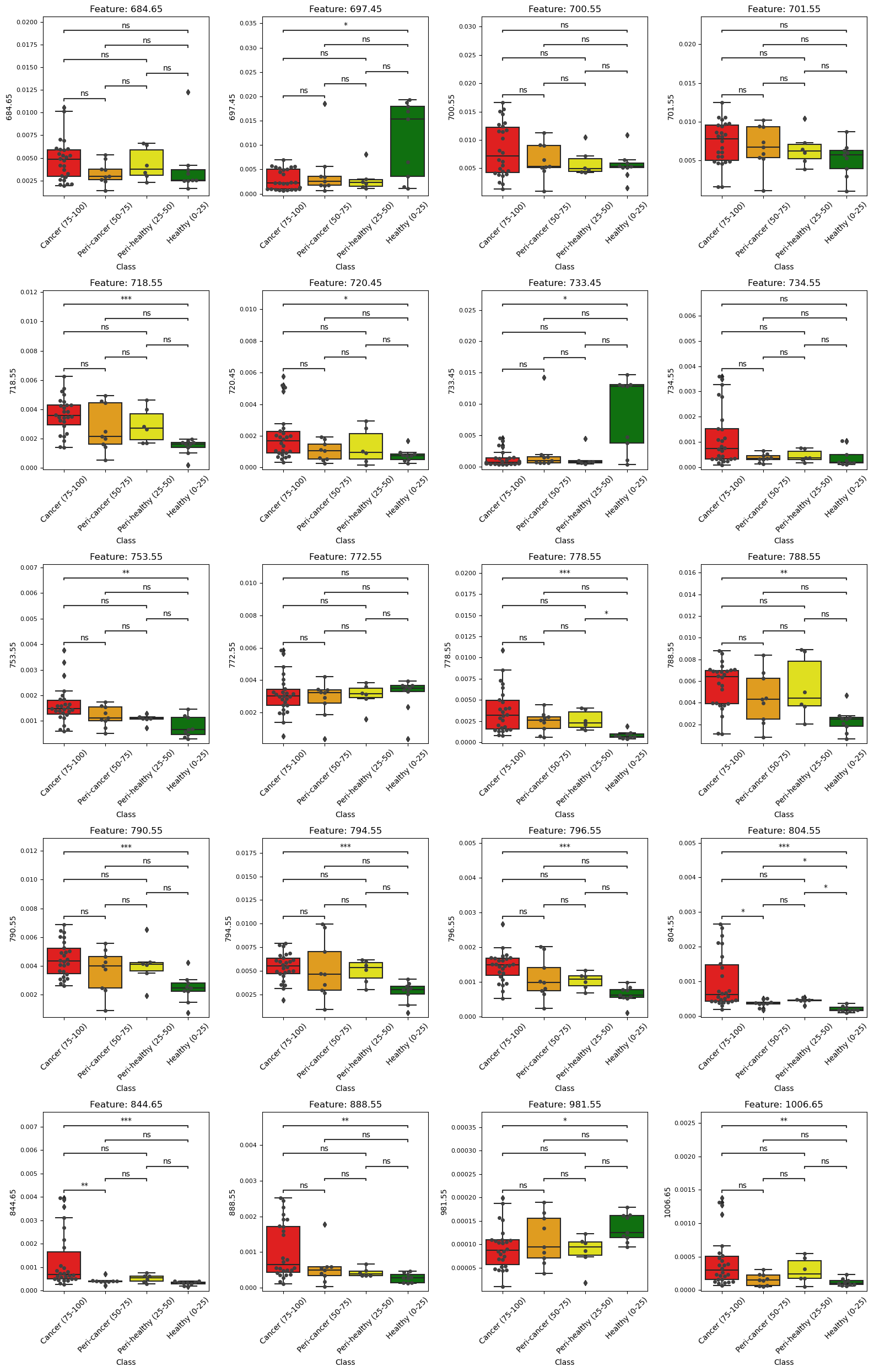

Table S4. Comprehensive list of reliable biomarkers for the global survival in both ion modes. Related to Fig. 4.

| *m/z* | **Survival outcome** | | **Possible annotation** |
| --- | --- | --- | --- |
|  | **< 2 years** | **> 2 years** |  |
| **Negative ion mode** | | | |
| 686.45 |  |  | / |
| 701.55 |  |  | [PA (36:1)-H]^-^ |
| 716.55 |  |  | [PE (34 :1)-H]^-^ |
| 728.55 |  |  | [PE (P-18:1_18:0)-H]^-^ |
| 736.55 |  |  | [PE (36:5)-H]^-^ |
| 738.55 |  |  | [PE (16:0_20:4)-H]^-^ |
| 744.55 |  |  | [PE (18:0_18:1)-H]^-^ |
| 748.55 |  |  | [PS (O-34:0)-H]^-^ |
| 764.55 |  |  | [PE (18:1_20:4)-H]^-^ |
| 766.55 |  |  | [PE (38:4)-H]^-^ |
| 772.55 |  |  | [PE (18:1_20:0)-H]^-^ |
| 792.55 |  |  | [PE (40:5)-H]^-^ |
| 819.55 |  |  | [PG (18:1_22:6)-H]^-^ |
| **Positive ion mode** | | | |
| 648.65 |  |  | / |
| 718.55 |  |  | / |
| 746.55 |  |  | / |
| 760.55 |  |  | [PS (34:2)+H]^+^ |
| 794.55 |  |  | [PE (40:5)+H]^+^ |
| 805.55 |  |  | [PA (44:6)+H]^+^ |
| 810.65 |  |  | [PE (O-42:4)+H]^+^ |
| 855.55 |  |  | [PI (36:6)+H]^+^ |
| 881.55 |  |  | / |
| 941.65 |  |  | [PI (42:5)+H]^+^ |
| 997.65 |  |  | [PI (46:5)+H]^+^ |

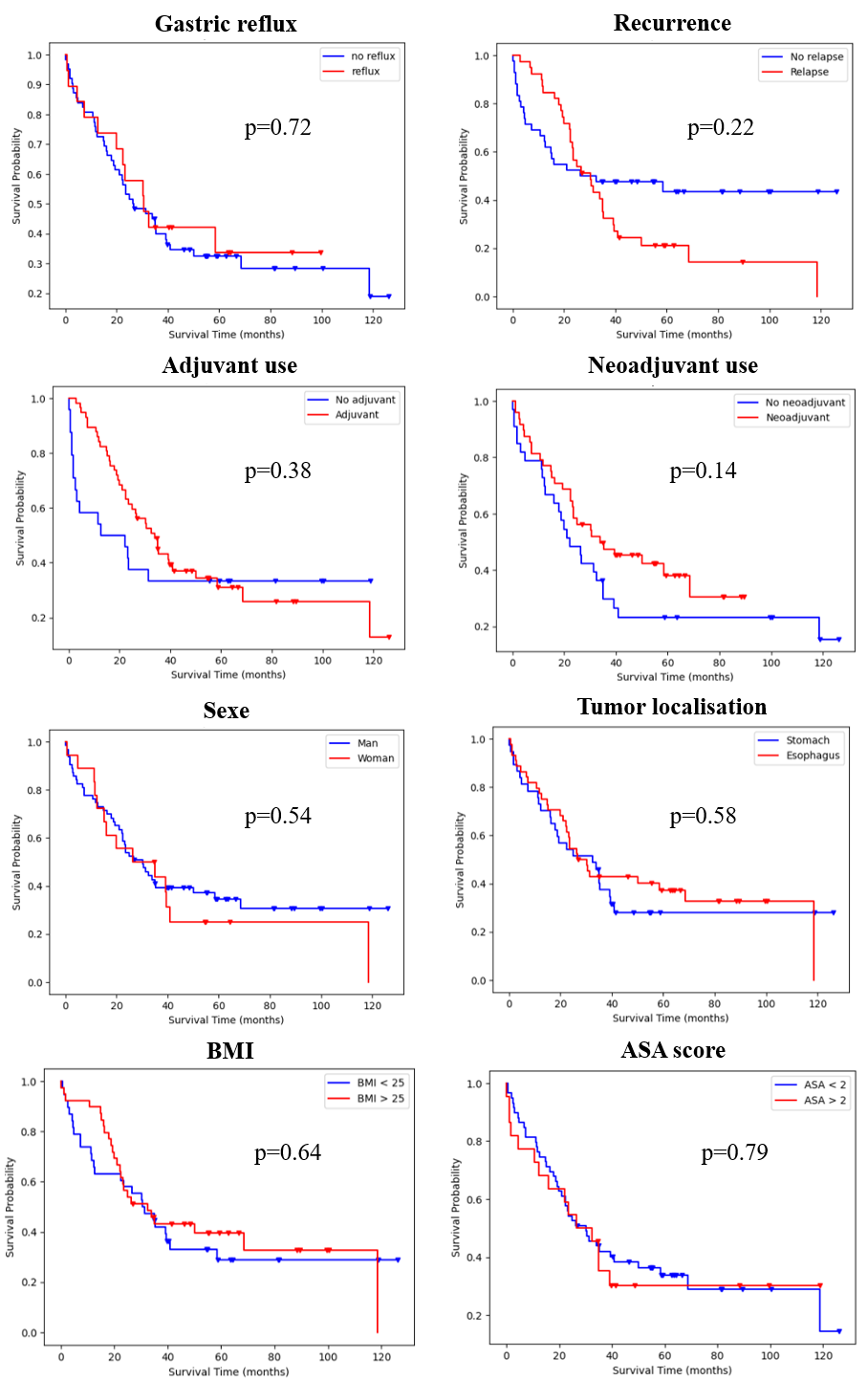

Figure S10. Survival curves (Kaplan Meier analysis) of all patients according to the gastric reflux, the BMI, the sexe, the tumor’s localisation, the recurrence, the adjuvant use and the neadjuvant use. Related to Fig. 4.

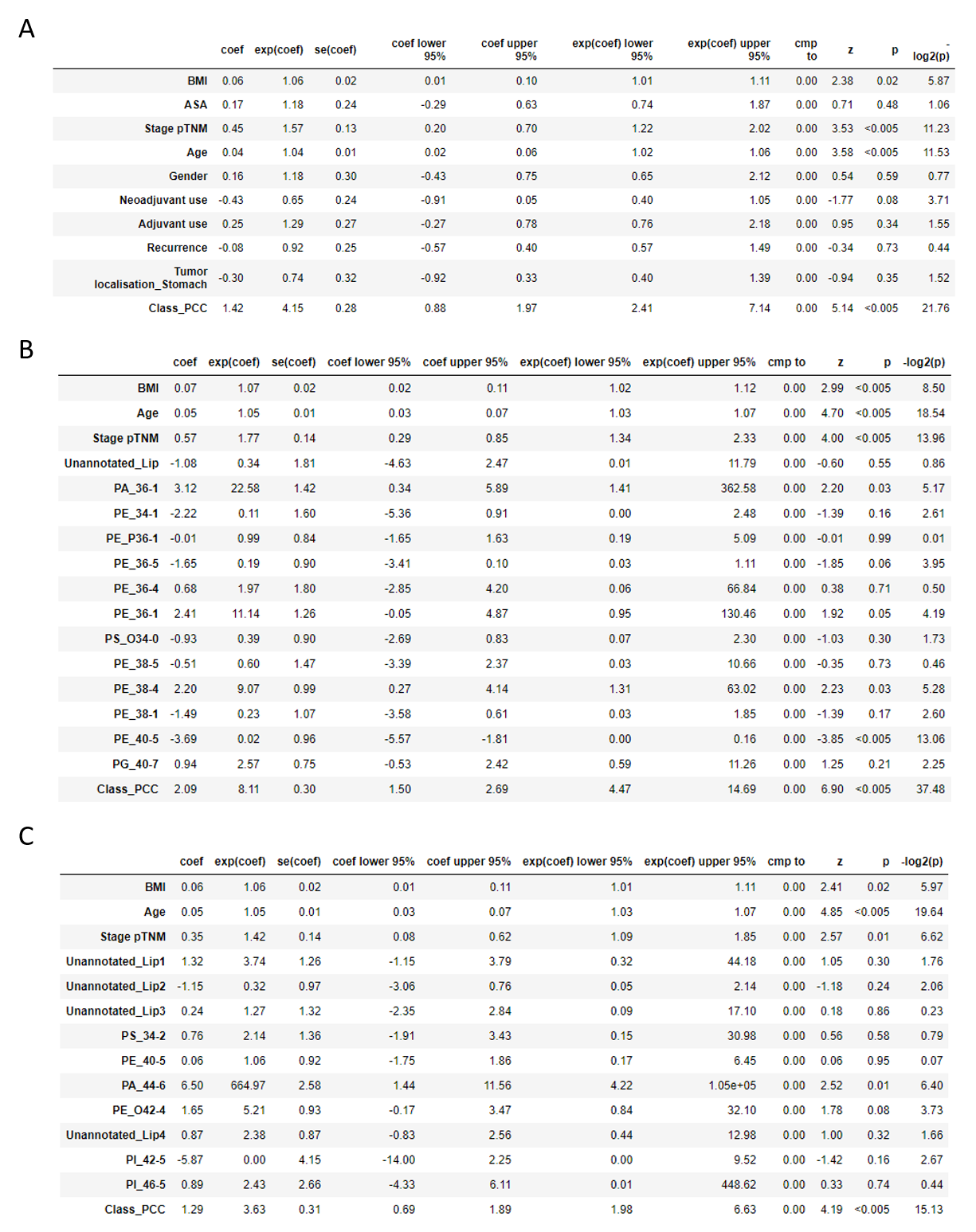

Figure S11. Summary tables of cox proportional hazard model coefficients made with (**A**) all clinical data available and with combined significant metadata and lipidsbiomarkers in (**B**) negative ion mode and (**C**) positive ion mode. Related to Fig. 4.

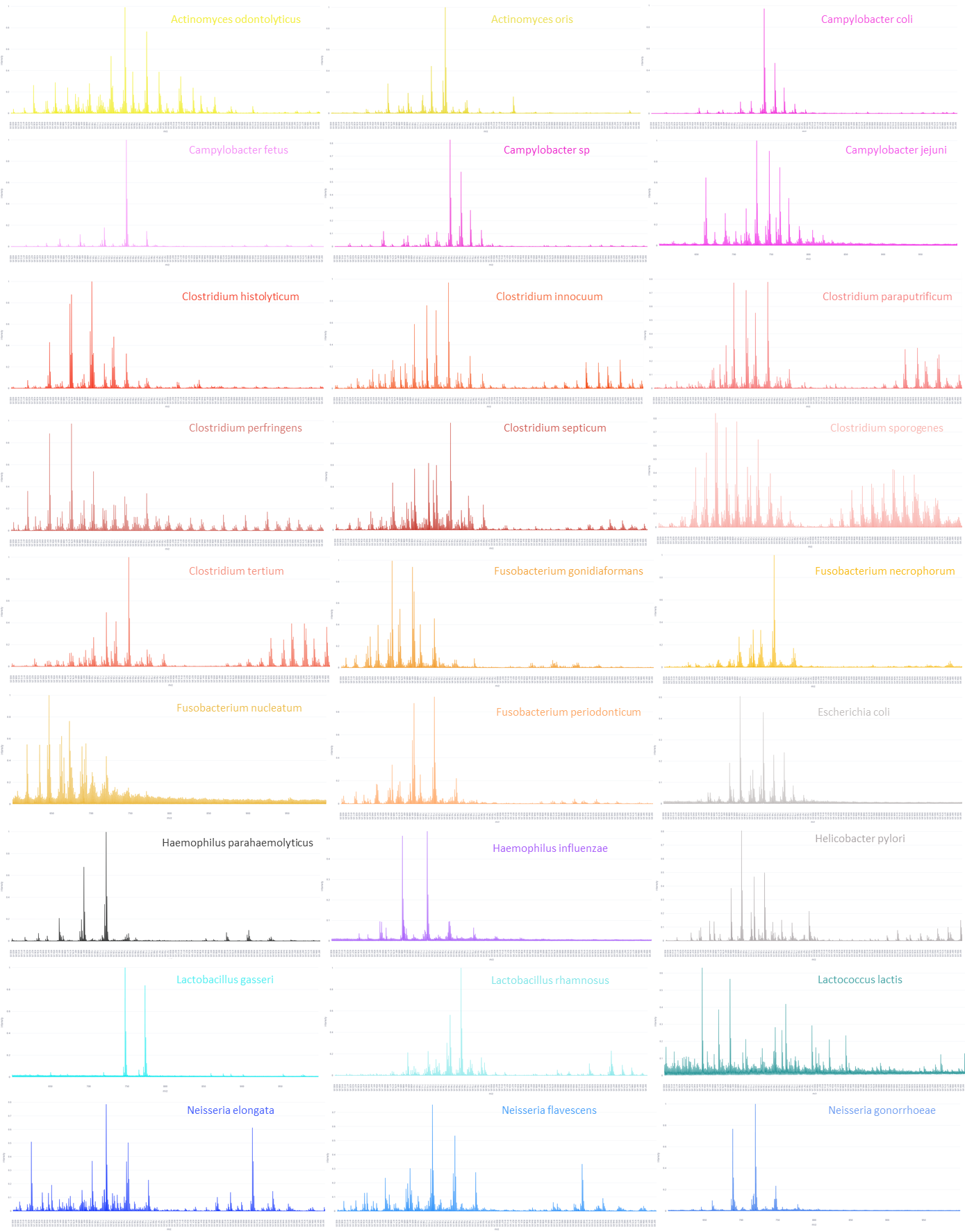

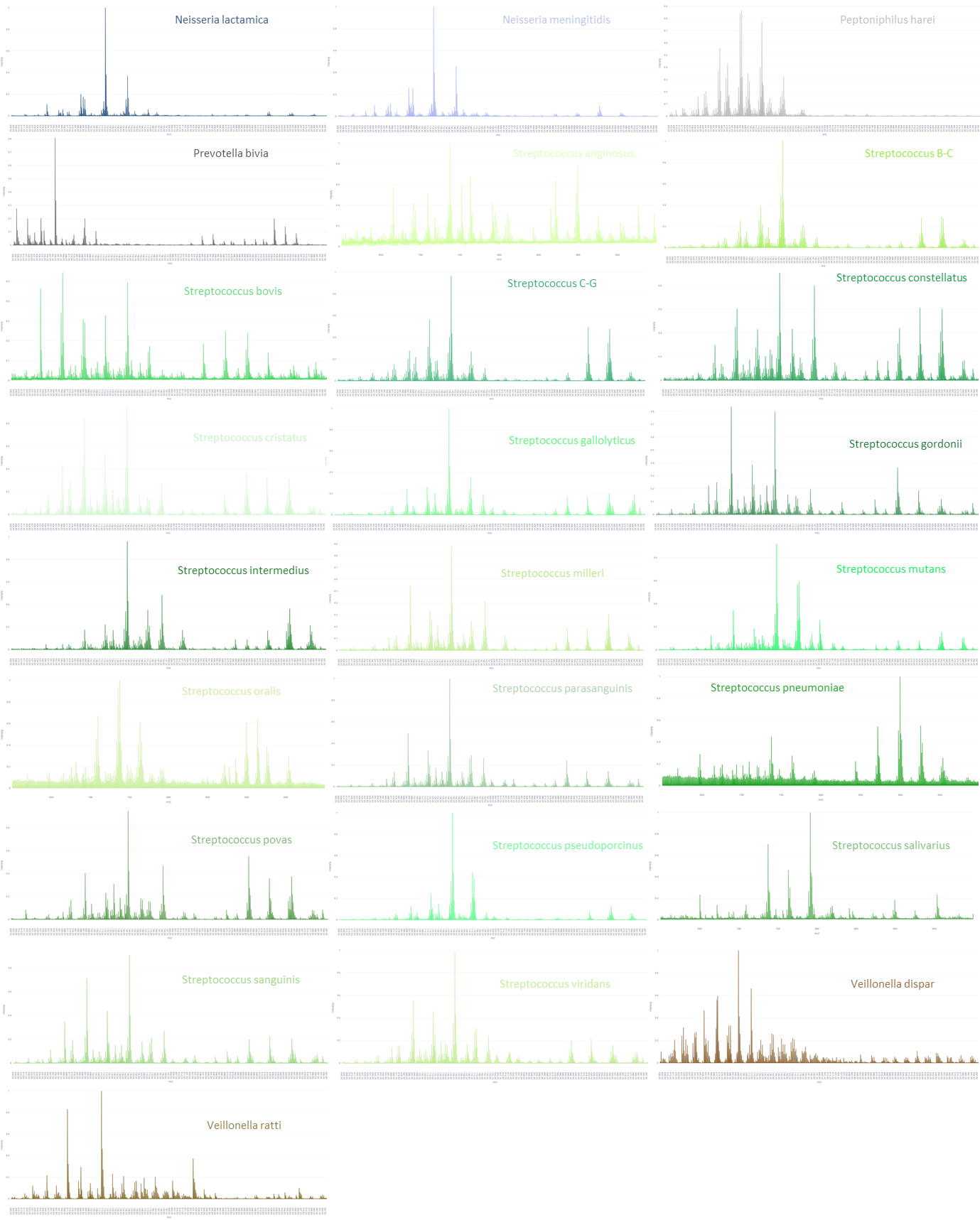

Figure S12. Mean spectra of all 52 bacterial strains in negative ion mode. Related to Fig. 5.

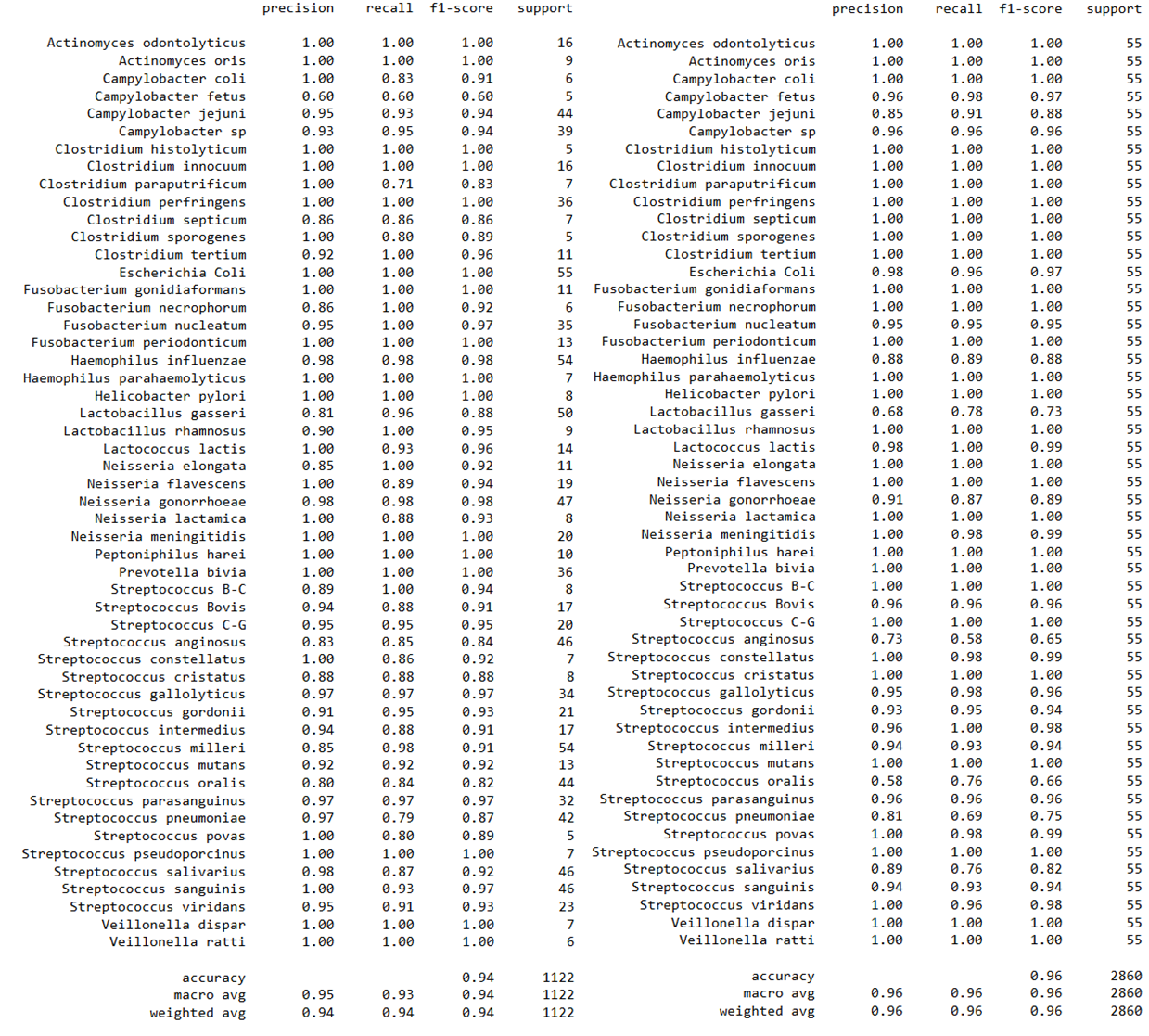
Figure S13. Classification report of bacterial strains models before oversampling, after 20-fold cross-validation in negative ion mode. Related to Fig. 5.

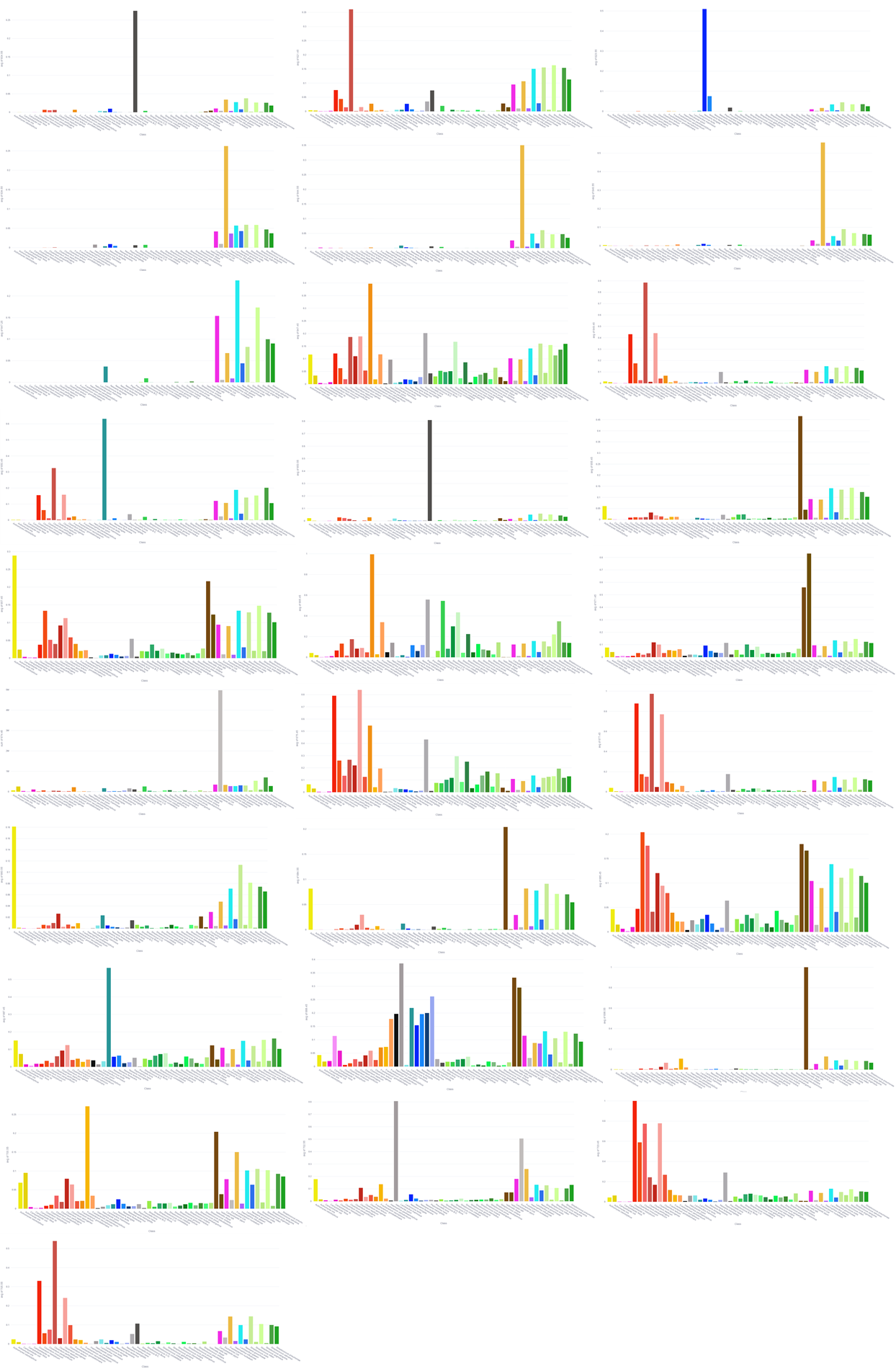

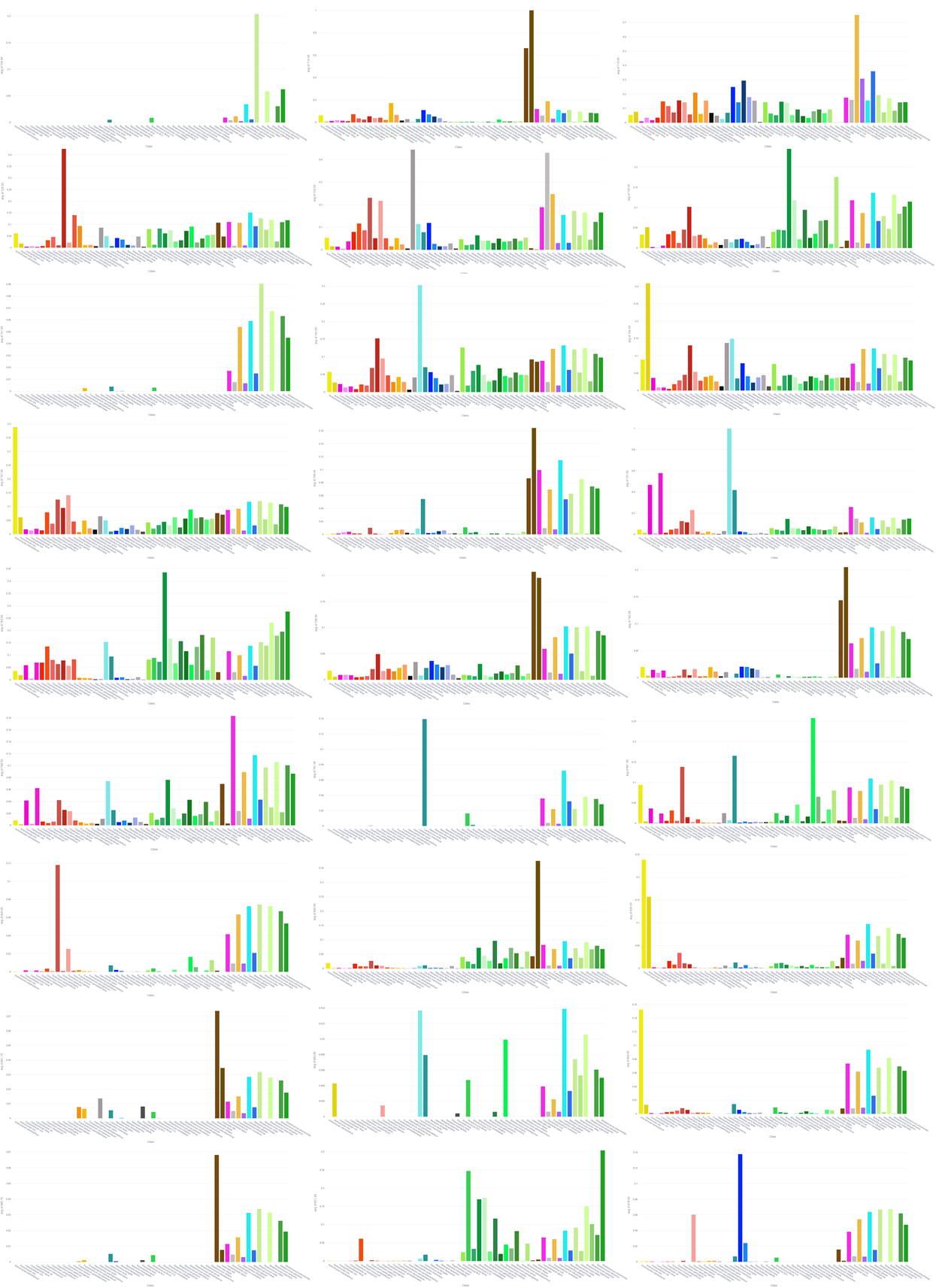

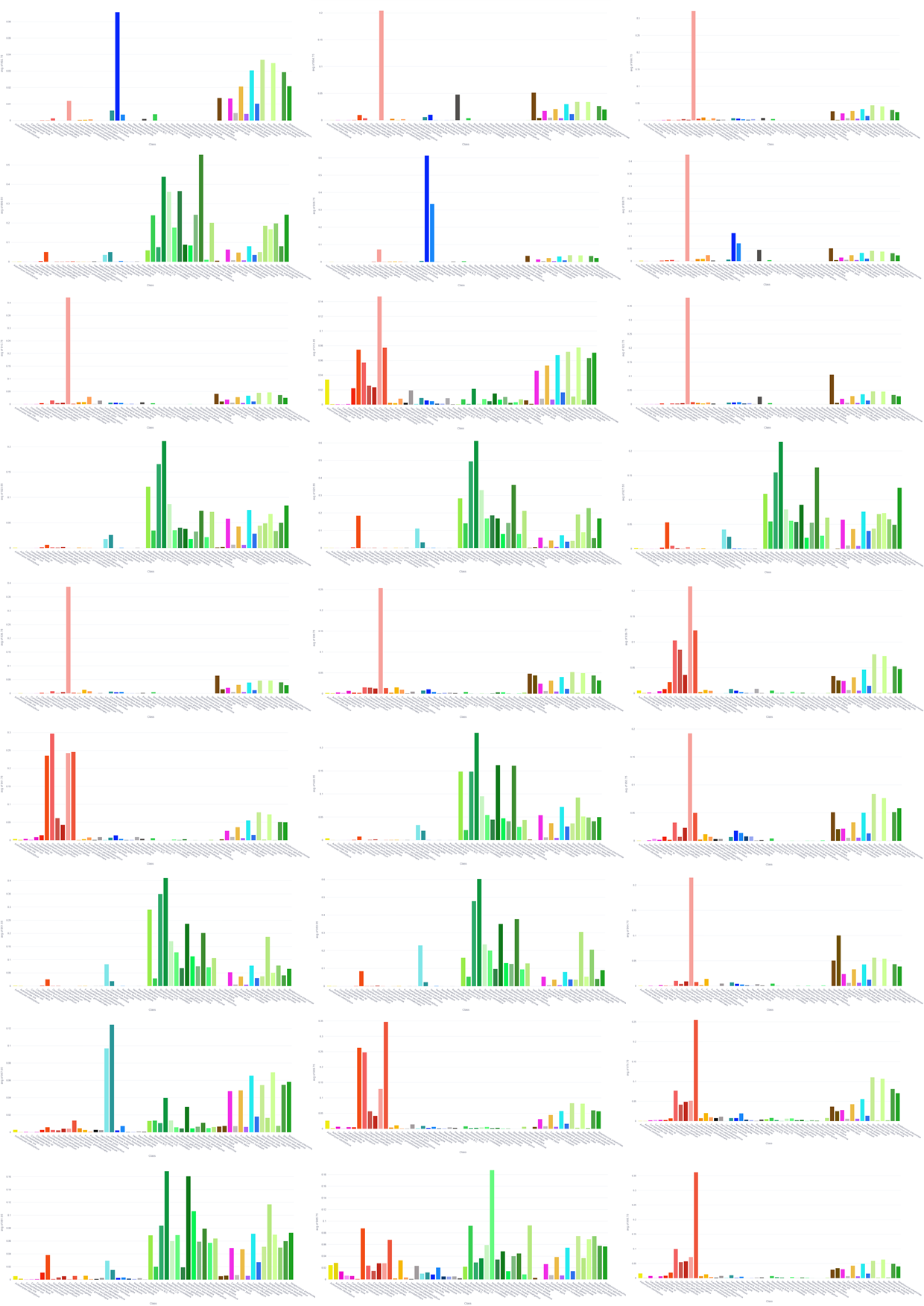

Figure S14. Mean intensity of each of the 82 lipid biomarkers in all the 52 bacterial strains. Related to Fig. 5.

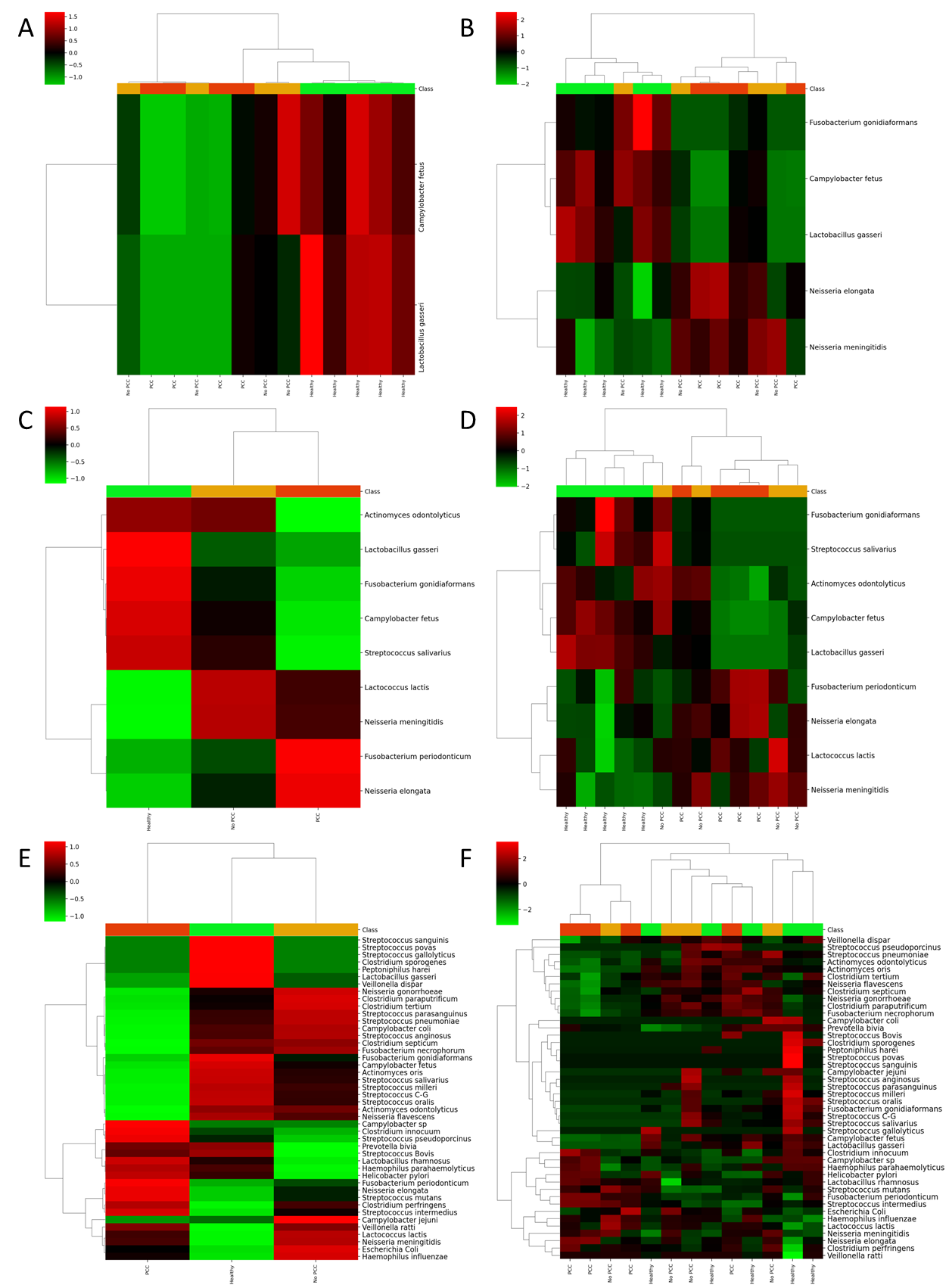

Figure S15. Heatmap following ANOVA analysis of: (**A**) tissues by tissues with a p-value of 0.01, (**B**) tissues by tissues with a p-value of 0.05, (**C-D**) average and tissues by tissues with a p-value of 0.1, (**E-F**) average and tissues by tissues with a p-value of 1, illustrating the predicted abundance of over-expressed and under-expressed bacteria in healthy, PCC, and non-PCC tissues. Related to Fig. 6.

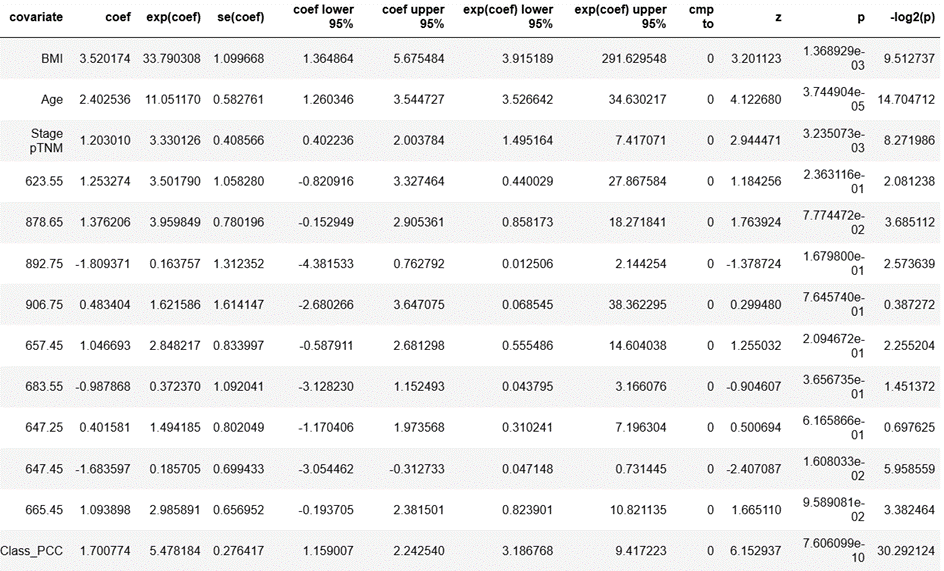

Figure S16. Summary table of Cox proportional hazard model coefficients made with both significant clinical data and specific bacteria lipids biomarkers. Related to Fig. 6.
